# Structure-based identification of naphthoquinones and derivatives as novel inhibitors of main protease Mpro and papain-like protease PLpro of SARS-CoV-2

**DOI:** 10.1101/2022.01.05.475095

**Authors:** Lucianna H. Santos, Thales Kronenberger, Renata G. Almeida, Elany B. Silva, Rafael E. O. Rocha, Joyce C. Oliveira, Luiza V. Barreto, Danielle Skinner, Pavla Fajtová, Miriam A. Giardini, Brendon Woodworth, Conner Bardine, André Luiz Lourenço, Charles S. Craik, Antti Poso, Larissa M. Podust, James H. McKerrow, Jair L. Siqueira-Neto, Anthony J. O’Donoghue, Eufrânio N. da Silva Júnior, Rafaela S. Ferreira

## Abstract

The worldwide COVID-19 pandemic caused by the coronavirus SARS-CoV-2 urgently demands novel direct antiviral treatments. The main protease (Mpro) and papain-like protease (PLpro) are attractive drug targets among coronaviruses due to their essential role in processing the polyproteins translated from the viral RNA. In the present work, we virtually screened 688 naphthoquinoidal compounds and derivatives against Mpro of SARS-CoV-2. Twenty-four derivatives were selected and evaluated in biochemical assays against Mpro using a novel fluorogenic substrate. In parallel, these compounds were also assayed with SARS-CoV-2 PLpro. Four compounds inhibited Mpro with half-maximal inhibitory concentration (IC_50_) values between 0.41 µM and 66 µM. In addition, eight compounds inhibited PLpro with IC_50_ ranging from 1.7 µM to 46 µM. Molecular dynamics simulations suggest stable binding modes for Mpro inhibitors with frequent interactions with residues in the S1 and S2 pockets of the active site. For two PLpro inhibitors, interactions occur in the S3 and S4 pockets. In summary, our structure-based computational and biochemical approach identified novel naphthoquinonal scaffolds that can be further explored as SARS-CoV-2 antivirals.

## 1. Introduction

COVID-19 is caused by a β-coronavirus that is related to the virus that was responsible for the severe acute respiratory syndrome (SARS) in 2003, and therefore designated SARS-CoV-2 [1]. In December 2019, the first cases of COVID-19 were reported in Wuhan, the capital of Hubei Province, China [2]. The new coronavirus showed a rapid geographical spread, associated with a high infection rate, and the World Health Organization (WHO) declared it as a pandemic on March 11, 2020 [3, 4]. The rapid transmission from human to human is undoubtedly the main source of contagion, which occurs mainly through droplets, hand contact, or contact with contaminated surfaces [5]. To control the spread of this pandemic virus, biosecurity and hygiene measures are now worldwide applied [6]. Despite the rapid development and emergency authorization of vaccines, viral escape mutants have emerged, and SARS-CoV-2 infections remain a concern for the global community. Therefore, there is a continuing need to discover structural frameworks for drugs that can be employed against COVID-19 [7].

Drug development efforts have targeted the SARS-CoV-2 main protease (Mpro) also known as 3-chymotrypsin-like protease (3CLpro) or non-structural protein 5 (nsp5) [8, 9]. Mpro is an essential cysteine protease that cleaves the precursor replicase polyprotein in a coordinated manner [10], to generate at least 11 non-structural proteins [11]. As a target, Mpro is conserved among other coronaviruses, and has no closely related human homolog [12–14]. Therefore, it has been intensively investigated as a drug target for SARS and Middle East Respiratory Syndrome (MERS) [15–18]. Several Mpro inhibitors with *in vitro* antiviral activity against SARS-CoV-2 have been reported [19–25], including peptidomimetic aldehydes (best IC_50_ values ranging ∼0.03-0.05 µM [19,21,23]), α-ketoamides (best IC_50_ values ranging ∼0.04-0.67 µM [20, 22]), calpain inhibitors (best IC_50_ values ranging ∼0.45-0.97 µM [20, 23]), nonpeptidic inhibitors (best IC_50_ values ranging ∼0.17-0.25 µM [24–26]). The binding modes of dozens of these inhibitors have been determined by crystallography [20,22–29]. Recently, a covalent reversible nitrile was reported as an orally bioavailable Mpro inhibitor with *in vitro* and *in vivo* antiviral activity [30], and shown to reduce hospitalizations in COVID-19 patients by 89% [31]. Coronaviruses also encode a second cysteine protease, PLpro, that plays an essential role in suppression of the host immune system [32–34]. PLpro can hydrolyze ubiquitin and interferon-induced gene 15 (ISG15) from host proteins which allows the virus to evade the host innate immune responses [10, 35]. This enzyme also cleaves the viral polypeptide to release the nsp1, nsp2 and nsp3 proteins [36]. SARS-CoV-2 PLpro inhibitors with antiviral efficacy have been described [37–40], including naphthalene-based (EC_50_ values of 1.4 to 21 µM for antiviral activity and IC_50_ values of 0.18 to 43.2 µM against the enzyme [37–39]) and 2-phenylthiophene-based (EC_50_ values of 2.5 to 11.3 µM for antiviral activity and IC_50_ values of 0.11 to 0.56 µM against the enzyme [40]) non-covalent compounds, and 21 crystallographic structures of this protease complexed with a ligand are available [8,37–42]. The crystal structures of Mpro and PLpro with bound ligands provided us with a structural basis to identify novel inhibitors.

Repurposing existing chemical libraries is a promising strategy to quickly discover novel therapies [43, 44]. Several newly discovered therapies for treatment of COVID-19 infection are derived from approved drugs, clinical candidates, and other pharmacologically active compounds that were originally developed for other indications [45–48]. In addition, knowledge gained from previous outbreaks of SARS, MERS, and bat coronavirus (BatCoV-RaTG13) have facilitated the rapid discovery of SARS-CoV-2 drugs [2,6,49]. Remdesivir, a broad-spectrum viral RNA-dependent RNA polymerase (RdRp) inhibitor [50, 51], was rapidly approved for treatment of hospitalized patients with COVID-19 [52], which has resulted in a more rapid recovery of patients and lower levels of airway infection [53]. Drugs that provide either symptom relief for patients or have not been scientifically proven to be effective are also being widely studied by the scientific community [54].

Embelin, a natural product with a quinone core, has antiviral activity against influenza and hepatitis B [55, 56]. Recently, it was shown that Embelin may inhibit Mpro and therefore have potential to be used as a treatment of SARS-CoV-2 [57]. In addition to Embelin, other studies showed that molecules containing a quinoidal framework also had inhibitory activity against SARS-CoV-2 Mpro. These included celastrol, pristimerin, tingenone and iguesterin [58]. We have experience working with naphthoquinones and therefore searched for structures with potential activity against SARS-CoV-2. In this report, we outline an *in silico* screening of a library of 688 quinonoid compounds and derivatives against SARS-CoV-2 Mpro, from which 24 compounds were selected and tested against this protease. Based on this strategy, and on experimental screening against PLpro as well, we report novel naphthoquinoidal inhibitors of both SARS-CoV-2 proteases. In addition to biochemical validation, molecular dynamics (MD) simulations indicated the stability of the Mpro and PLpro quinoidal complexes binding modes, mediated by interactions that were also frequently found in crystallographic complexes of the proteases. The quinones are promising COVID-19 drug candidates to be further explored, while also offering valuable insights into Mpro and PLpro inhibition.

## 2. Results

### 2.1. Assembly of a chemical library for virtual screening against Mpro

To search for potential Mpro inhibitors, we retrieved a library of quinones and their derivatives, as detailed in **Figures 1** and **2** (See the Supporting Information **Figures S1-S40**, for more structural information). Six hundred and eighty-eight compounds were considered for virtual screening by molecular docking. We divided the molecules into eight different groups as described in **Figure 1** (Groups 1-4) and **Figure 2** (Groups 5-8). The compounds listed in Group 1 are *ortho*-quinones with different substitution patterns. In general, we evaluated compounds containing arylamino [59–64], alcohol [60] and alkoxy groups [61, 62], selenium and sulfur [65–67], the basic chalcone framework [68], among simple *ortho*-quinones [61,68–71].

**Figure 1.**
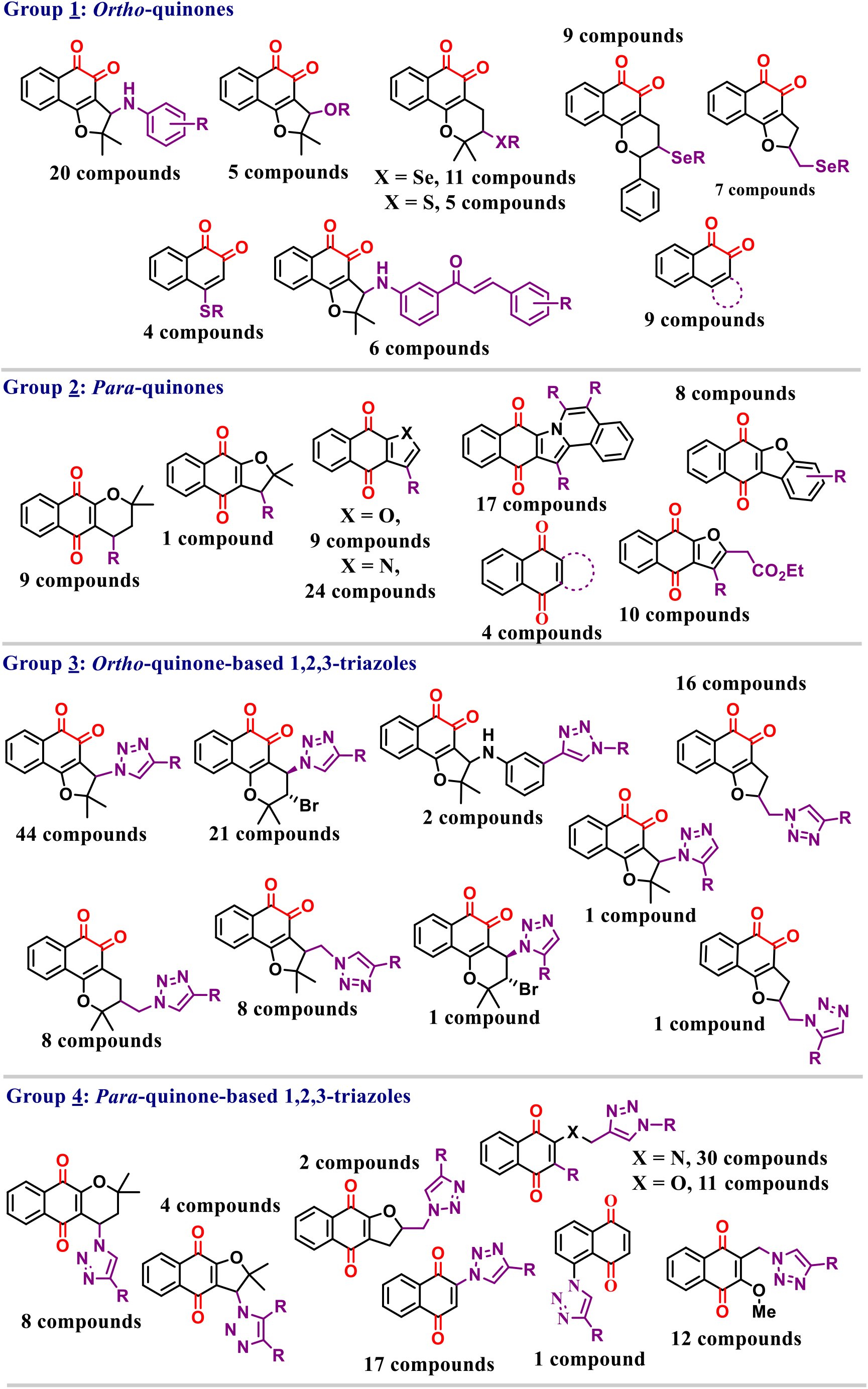
The basic structural framework of the compounds listed in Groups 1-4.

**Figure 2.**
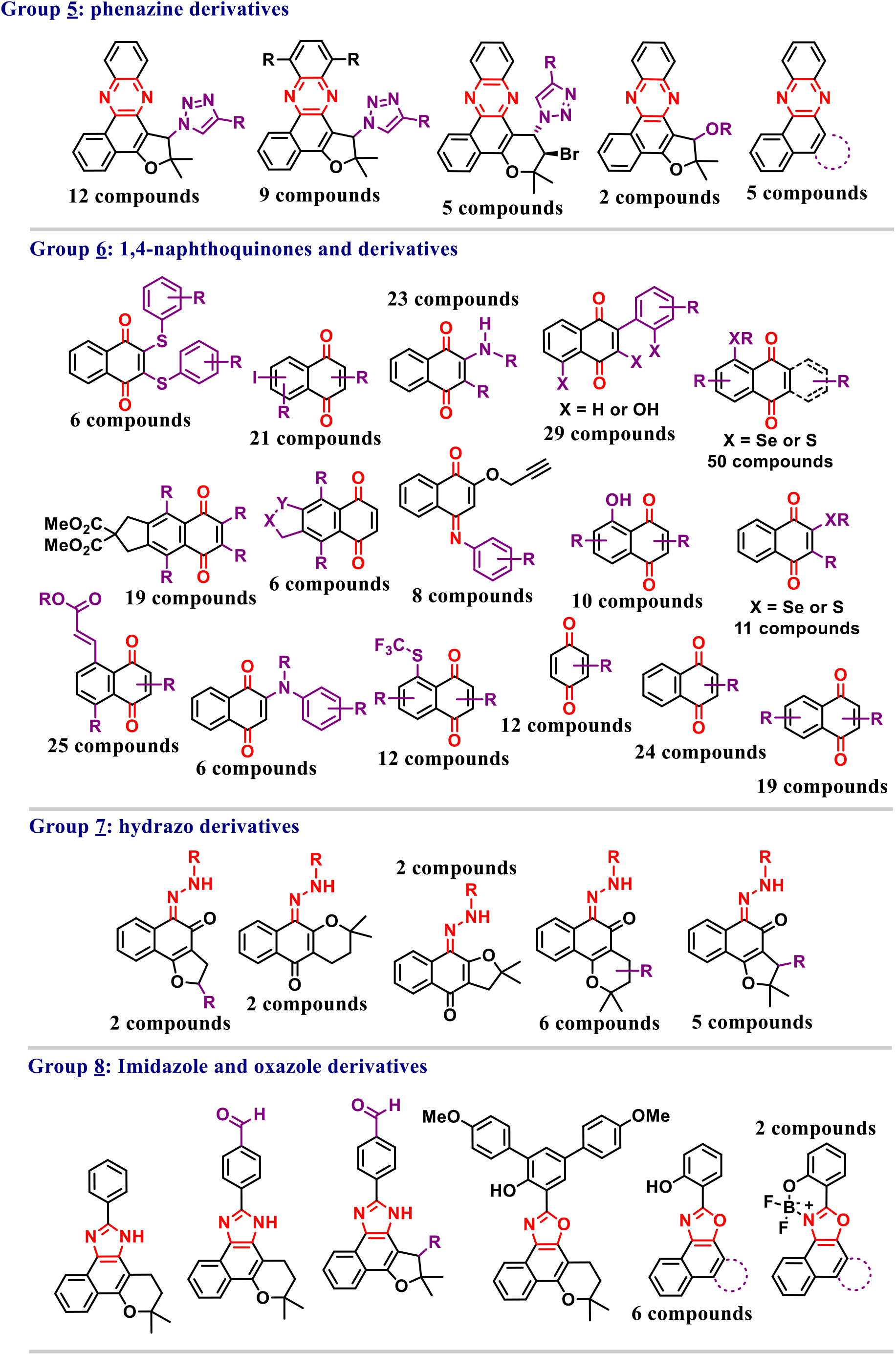
The basic structural framework of the compounds listed in Groups 5-8.

Group 2 is composed of *para*-quinones. We studied compounds such as α*-* lapachones [60, 72], arylamino derivatives [60,64,71], furanonaphthoquinones [73–75], and pyrrolonaphthoquinones [73, 76], in addition to other derivatives based on *para*-quinones [64, 75]. The selected compounds for this group exhibit a broad substitution pattern but, in general, arylamino and aryl groups are often observed. Compounds with antiviral activity containing the *para*-quinone core are frequently described in the literature [77–79].

Groups 3 and 4 consist of *ortho*- and *para*-quinones with a 1,2,3-triazole nucleus. Lapachone-based 1,2,3-triazoles have been studied because of their broad spectrum of biological activities. We studied compounds with aromatic and aliphatic substituents [61,71,80–86], the presence of selenium [64, 87], BODIPY [88, 89], and sugars [71], among other substituent groups in the present quinoid structure [45,90–93].

The phenazine form of the triazole compounds and quinones described in groups 1 and 3 were also evaluated in group 5 [94–97]. Group 6 is the most complete and diverse group addressed in this study, containing approximately two hundred 1,4-naphthoquinones with broad substitution patterns in the benzenoid A-ring and B-ring. Compounds containing sulfur, as sulfoxides and sulfones [98, 99], selenium [100], iodine [47], amines, bromine, hydroxyls, alkenes, among other substituent groups [46,101–107] were studied and evaluated according to their potential to act as anti-SARS-CoV-2. Imine derivatives were also targeted in our studies and were placed in group 6 [67].

Finally, groups 7 and 8 are formed by hydrazo, imidazole, and oxazole derivatives [108–111]. The compounds in these groups were prepared from the quinones described above and represent our attempt to study quinone-derived heterocyclic compounds with biological activity against various microorganisms and their effectiveness against the virus that causes COVID-19.

### 2.2. Available Mpro structures show conserved conformation, protein-ligand interactions, and location of waters molecules

As an initial investigation to support the virtual screening of the quinoidal library, we analyzed 72 Mpro structures with bound ligands that have a resolution of 1.3 Å to 2.5 Å (Supporting Information **Figure S41B**). Active Mpro forms a homodimer comprising two protomers [19], while its monomer is inactive [112]. Each protomer is formed by domains I, II, and III, binding to each other by an *N*-terminal finger between domains II and III [24] (**Figure 3**). The substrate-binding site is located in a cleft between domains I and II and covered by a loop connecting them. Also crucial to the formation of the active dimer, the *N*-terminal finger of one monomer extends to the other monomer, shaping and forming the active substrate-binding site [22]. The substrate-binding cleft is composed of four subsites S1ʹ, S1, S2, and S4 [14, 19], which features a non-canonical Cys-His catalytic dyad [19, 24] (**Figure 3**).

**Figure 3.**
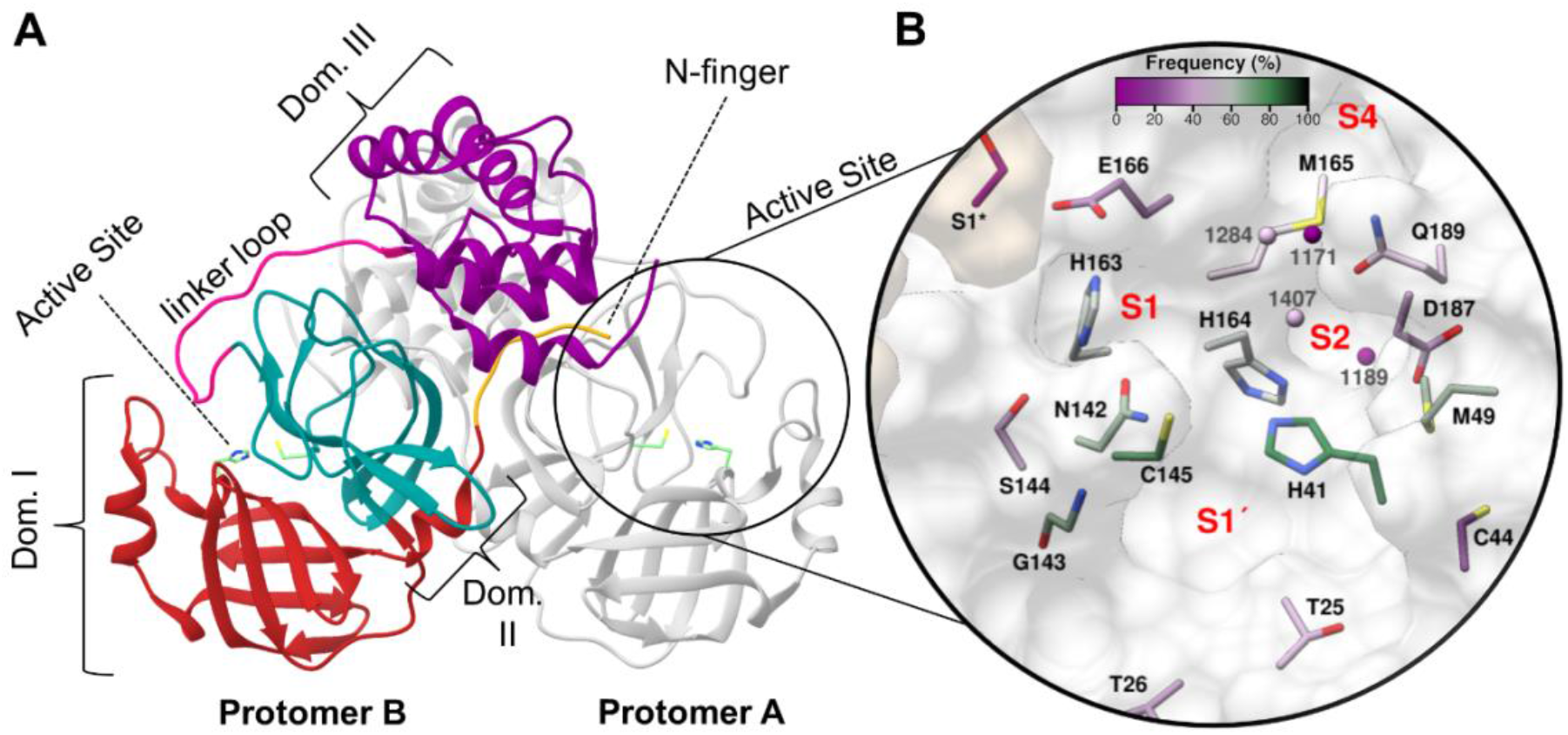
Three-dimensional structure of Mpro dimer (PDB code: 5R82 [117]) and surface view of the active site. Protomer A is shown in gray, and protomer B is colored according to three domains: red for domain I, blue for domain II, and purple for domain III. The loop linking domains II and III, critical for the protein dimerization, is colored in pink and the *N*-terminal finger colored orange (A). The substrate-binding cleft is highlighted with the catalytic residues H_41_ and C_145_ displayed as sticks (B). In the close-up view of the active site, the residues and conserved water molecules are colored by the frequency they are involved in interactions with 72 crystallography ligands, according to an analysis using the program LUNA (https://github.com/keiserlab/LUNA).

Using principal component analysis (PCA) to assess conformational differences among the structures, we found a high similarity among the Mpro structures evaluated. Even for the four most divergent structures (PDB codes: 6M2N [113], 6W63 [114], 6LU7 [24], and 7BQY [24], Supporting Information **Figure S41A**), carbon alpha (Cα) root-mean-square deviation (RMSD) between the protease structures is less than 1.0 Å (Supporting Information **Figure S41B**), suggesting high overall conservation of the quaternary structure.

On the other hand, the Mpro active site is known for its high flexibility with conformational changes induced by ligand binding [24,115,116]. Thus, to evaluate possible differences in active site residue conformations, we superimposed six high-resolution Mpro structures (1.31 Å to 1.51 Å, PDB codes 5R82, 5RFW, 5RF6, 5RFE, 5RFV, and 5RF3 [117]) with four structures that were discovered to have lower structural similarity by PCA and had resolutions between 2.10 Å and 2.20 Å. The superposition of these structures reveals that most residues in the ligand-binding site adopt similar conformations (Supporting Information **Figure S42**), except for M_49_, N_142_, M_165_, and Q_189_, which were the most flexible among the other binding site residues.

To better understand the molecular recognition between Mpro and inhibitors, we assessed the common protein-ligand interactions found in the 72 experimentally determined crystal structures, of which 49 displayed covalent and 23 non-covalent ligands, using the program LUNA (https://github.com/keiserlab/LUNA). Within this comprehensive set, ligands interacted most frequently with the catalytic dyad and residues in the S1 and S2 pockets (**Figure 3B**). For the catalytic dyad, C_145_ interacts with 78% of the ligands, forming hydrophobic and hydrogen bond interactions, while H_41_ binds to 82% of the inhibitors, mostly through aromatic stacking, hydrophobic, cation-π, and weak hydrogen bond interactions (**Figure 3B** and **4A**). Within the S1 pocket, polar protein-ligand interactions were enriched such as hydrogen bonds and hydrophobic interactions with G_143_ (68% interaction frequency); hydrophobic interactions and weak hydrogen bonds with N_142_ (65% interaction frequency); and cation-nucleophile, cation-π, and hydrogen bond interactions with H_163_ (58% interaction frequency). S_144_ (35%) and E_166_ (29%) in the S1 pocket and D_187_ (33%), and C_44_ (21%) in S2 had lower frequency of interactions (**Figure 3B** and **4A**). On the other hand, the S2 subsite is more hydrophobic. The two residues with the highest interaction frequencies from this pocket were M_49_ (65%) and H_164_ (58%) which formed hydrophobic and weak hydrogen bond interactions, while M_165_ (47%) and Q_189_ (43%) interacted mainly by hydrophobic contacts with the ligands (**Figure 3B** and **4A**). The high frequency of interactions with S1 and S2 residues showed that most of the ligands fill one or both pockets, conserving a more polar profile for S1, whereas the S2 retained a more aromatic and aliphatic profile as observed previously with the SARS-CoV Mpro [118] and in other studies with SARS-CoV-2 Mpro [119, 120].

**Figure 4.**
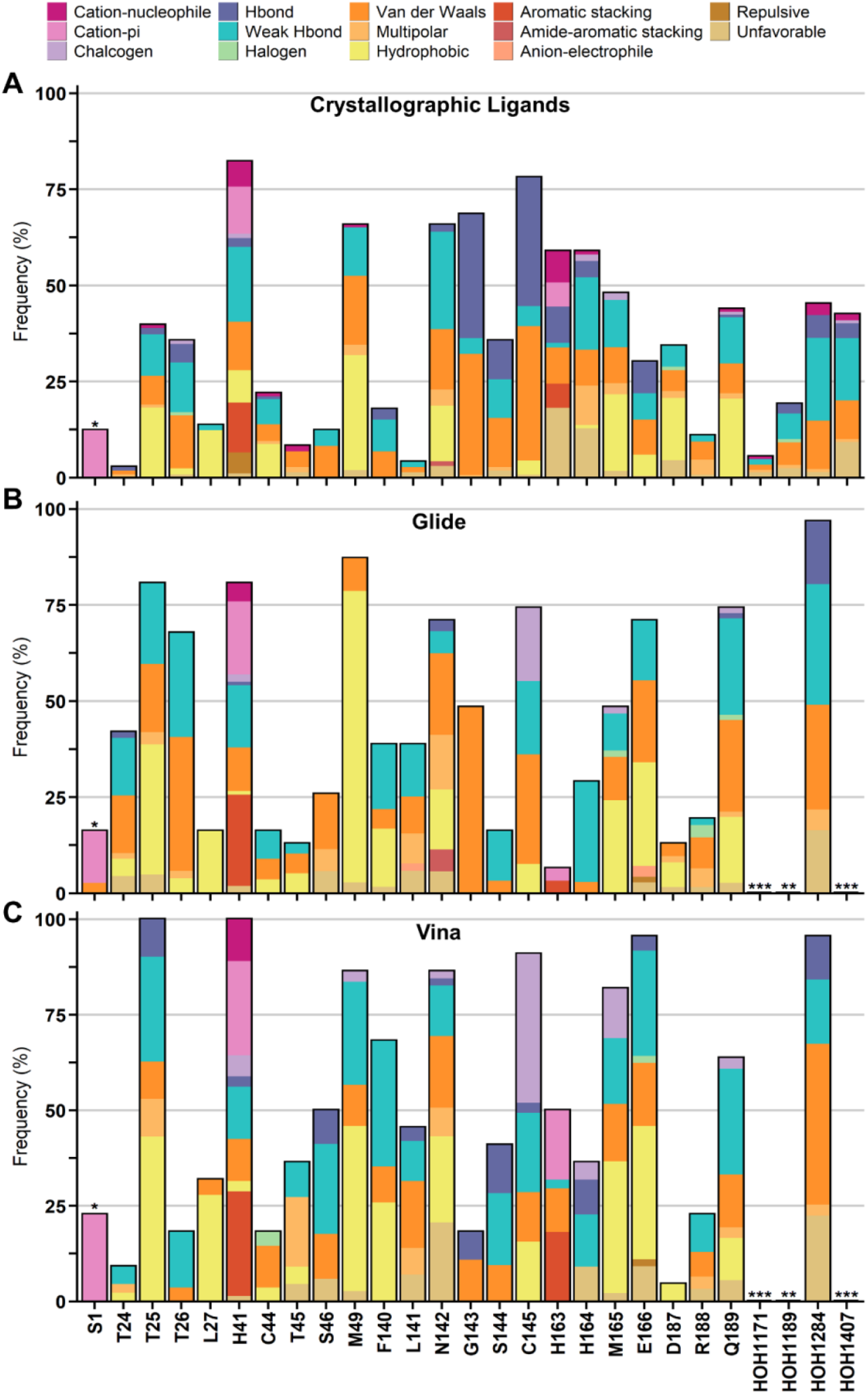
Interaction analysis between Mpro binding site residues and ligands from crystallographic structures and docking with Glide and Vina. Frequency and interaction types between residues and 72 crystallographic ligands (A), the final 24 selected compounds for biochemical assays from both Glide (B) and Vina (C). Residues with (*) are from the other protomer. From docking results, the (**) highlight residues with no interactions, while (***) are residues that were not considered in the docking calculations.

Additionally, two S1’ residues, T_25_ (39%) and T_26_ (35%), displayed frequent hydrophobic and weak hydrogen bond interactions. Amino acids in more solvent-exposed pockets, such as S1’ residues L_27_ (13%) and T_24_ (3%), and S4 residues retained few or no interactions (**Figure 4A**). Several of the hydrogen bond interactions found by Luna were mediated by water, meaning ligand and protein residue are bridged by a solvent molecule. In Mpro, water molecules contribute to ligand stabilization by forming water-mediated hydrogen bonds [19,20,24] and act as a possible third element to the catalytic dyad [9,121,122]. Therefore, we investigated which waters are conserved among the chosen Mpro structures using the ProBiS H2O plugin [123]. We found four conserved water molecules (present in over 50% of the structures, **Figure 3** and Supporting Information **Table S1**), that interacted with 20-45% of the ligands, displaying van der Waals, hydrogen bond, and weak hydrogen bond interactions (**Figure 4A**). Thus, these crystallographic conserved and buried water molecules might be important for ligand recognition.

### 2.3. Virtual screening of naphthoquinoidal compounds against SARS-CoV-2 main protease

Considering the high conservation within the Mpro crystal structure conformations, we performed initial molecular docking experiments with the highest-resolution structure (1.31 Å – PDB Code 5R82 [117]) from the most populated of the structural clusters, followed by a second round of flexible docking with compounds prioritized from rigid docking. Due to the importance of water molecules in the ligand binding site, we retained two of the four conserved water molecules, that may mediate hydrogen bonds with H_41_, C_145_, E_166_, and L_167_, for molecular docking (**Figure 4A** and Supporting Information **Table S1**). To account for the possibility of water displacement by ligands, a second Mpro preparation was also performed in the absence of water molecules. Both preparations were submitted to two distinct docking algorithms, Glide [124] and Autodock Vina [125].

Docking results were visually inspected and relevant poses were selected according to their overall binding site complementarity and specific protein-ligand interactions. Thus, we prioritized 70 compounds that interacted with the previously established high frequent residues, H_41_, M_49_, N_142_, G_143_, C_145_, M_165_, Q_189_, and water molecules for flexible docking approaches. Overall, Glide and Vina docking modes established contacts with S1’, S1, and S2 residues. However, a slight shift in interaction patterns was found. Compounds from the quinoidal library did not establish as many hydrogen bond interactions as the crystallographic ligands, giving a more hydrophobic nature to the interactions (**Figures 4B/C** and Supporting Information **Figure S43**).

In the second round of docking, we treated M_49_, N_142_, M_165_, and E_189_ as flexible residues, as these were most flexible within crystal structures analyzed and interacted with a high number of ligands (40-65%). Based on these results, we selected 24 (out of 70) compounds that matched the desired residue interactions (**Figure 4B** and **C**) and maintained good complementarity to the binding site (Supporting Information **Figures S44** to **S47**), for experimental validation in biochemical assays. The compounds selected represent diverse scaffolds from our library, comprising *ortho*-quinone-based 1,2,3 triazoles (group 3), *para*-quinone-based 1,2,3 triazoles (group 4), 1,4-naphthoquinones (group 6), and hydrazo derivatives (group 7).

### 2.4. Design and validation of Mpro substrate

Prior to biochemically evaluating the compounds against Mpro, we designed a fluorescent-quenched peptide substrate with the sequence ATLQAIAS that corresponds to the P4 to P4ʹ amino acids of the nsp7-nsp8 cleavage site and the dash representing the scissile bond. This substrate was chosen because the sequence most closely matches the consensus sequence for all 11 viral polypeptide cleavage sites (**Figure 5A** and **B**) [126]. ATLQAIAS was flanked by 7-methoxycoumarin-4-acetyl-L-lysine on the N-terminus, dinitrophenyl-L-lysine on the C-terminus. The peptide contains several non-polar amino acid residues and therefore two d-Arginine residues were added on the N-terminus to increase solubility. Using a concentration range of 3 µM to 250 µM, the K_M_ for this substrate was calculated to be 52.1 µM ± 14.4 µM.

**Figure 5.**
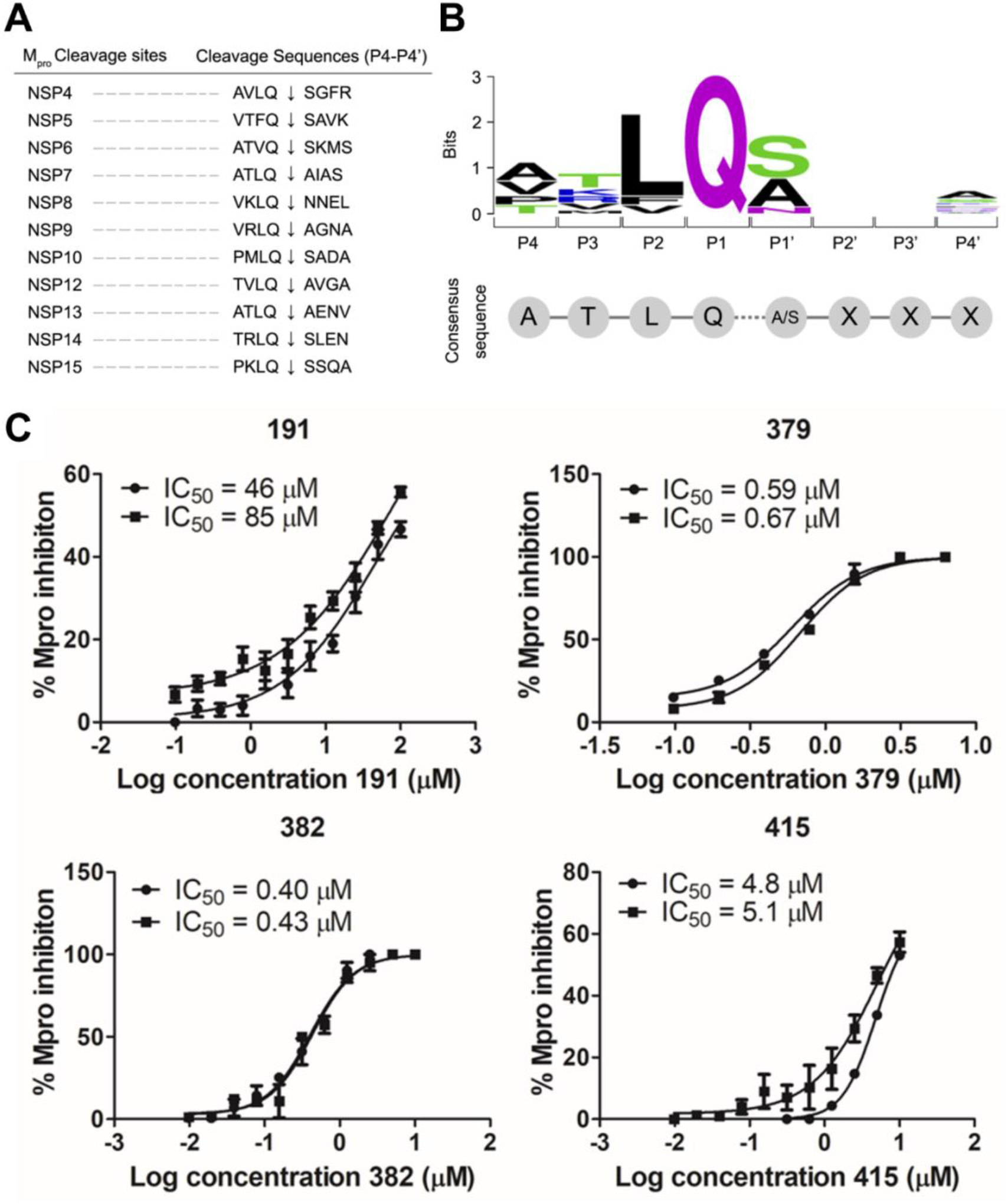
Validation of SARS-CoV-2 Mpro inhibitors in enzyme assays. List of the 11 Mpro cleavage sites (A) and the design of a fluorescent-quenched peptide substrate (B). The ATLQAIAS substrate was chosen since it closely matches the consensus sequence of all 11 viral polypeptide cleavage sites. IC_50_ curves for SARS-CoV-2 Mpro inhibitors (C). For each compound, two IC_50_ curves are shown, corresponding to two independent experiments (data shown as spheres or squares for each experiment), in which the compounds were pre-incubated with Mpro prior to substrate addition. Each curve was determined based on at least 7 compound concentrations in triplicate.

### 2.5. Validation of novel Mpro inhibitors

We evaluated the 24 hit compounds from our virtual screen in a biochemical assay using recombinant SARS-CoV-2 Mpro. The enzyme was pre-incubated with each compound at 10 µM and then assayed with the fluorogenic peptide substrate. To avoid detecting aggregators as false positives [127, 128], our assay was performed in the presence of 0.01% Tween 20. Additionally, we evaluated the absorbance of MCA fluorescence by the compounds, to make sure the observed enzyme inhibition was not an artifact of fluorescence, another common cause of false positives in enzyme assays [129]. From this screen, three 1,4-naphthoquinones derivatives, **379**, **382**, and **415**, fully inhibited Mpro, while two quinone-based 1,2,3 triazoles, **191** and **194**, had 50% or more inhibition. **668** was insoluble in assay buffer and was therefore eliminated from further analysis, while the remaining compounds had inhibition profiles of less than 50% (**Table 1**). The most potent compounds were subsequently evaluated at a concentration range of 10 µM to 9.7 nM and the half-maximal inhibitory concentration (IC_50_) was calculated to be 66 µM ± 22 for **191**, an *ortho*-quinone-based 1,2,3 triazole, 5 µM ± 0.15 for **415**, 0.63 µM ± 0.04 for **379**, and 0.41 µM ± 0.015 for **382** (**Table 1** and **Figure 5**).

**Table 1.** Percentage of inhibition at 10 µM and IC_50_ for naphtoquinoidal compounds against SARS-CoV-2 Mpro and PLpro.

| Compound<sup>a</sup> | Mpro |  | PLpro |  |
| --- | --- | --- | --- | --- |
| | % inhibition at 10 $\mu\text{M}$<sup>b</sup> | $\text{IC}_{50}$ ($\mu\text{M}$)<sup>c</sup> | % inhibition at 10 $\mu\text{M}$<sup>b</sup> | $\text{IC}_{50}$ ($\mu\text{M}$)<sup>c</sup> |
| 159 | 35 ± 1 | ND | 100 ± 1 | 1.7 ± 0.1 |
| 189 | 25 ± 2 | ND | 100 ± 0 | 2.2 ± 0.2 |
| 191 | 55 ± 3 | 66 ± 20 | 93 ± 0.7 | 3.1 ± 0.9 |
| 193 | 32 ± 1 | ND | 76 ± 1 | 46.5 ± 5 |
| 194 | 50 ± 7 | ND | 78 ± 5 | 29 ± 4 |
| 195 | 40 ± 2 | ND | 80 ± 2 | 33.5 ± 4 |
| 196 | 29 ± 2 | ND | 89 ± 2 | 22.5 ± 5 |
| 197 | 18 ± 4 | ND | 80 ± 2 | 20.5 ± 2 |
| 314 | 36 ± 2 | ND | 55 ± 7 | ND |
| 318 | 45 ± 13 | ND | 9 ± 5 | ND |
| 319 | 5 ± 2 | ND | 69 ± 3 | ND |
| 320 | 11 ± 6 | ND | 31 ± 6 | ND |
| 321 | 40 ± 8 | ND | 17 ± 0.2 | ND |
| 379 | 100 ± 0 | 0.63 ± 0.04 | 0 ± 0 | ND |
| 380 | 12 ± 3 | ND | 12 ± 6 | ND |
| 382 | 100 ± 0 | 0.41 ± 0.02 | 0 ± 0 | ND |
| 414 | 29 ± 2 | ND | 38 ± 2 | ND |
| 415 | 100 ± 0 | 5 ± 0.2 | 63 ± 9 | ND |
| 465 | 3 ± 5 | ND | 0 ± 0 | ND |
| 470 | 1 ± 2 | ND | 6 ± 0 | ND |
| 477 | 5 ± 2 | ND | 0 ± 0 | ND |
| 666 | 5 ± 5 | ND | 0 ± 0 | ND |
| 668 | NT | NT | NT | NT |
| 673 | 8 ± 3 | ND | 4 ± 4 | ND |

<sup>a</sup>See Supporting Information File for all structures. <sup>b</sup>Percentages of inhibition are reported as averages and standard deviation of the mean calculated from one experiment performed in triplicate. Compounds were pre-incubated with enzymes for 15 min before addition of the substrate. <sup>c</sup> $\text{IC}_{50}$ values are reported as the averages and standard deviation of the mean, based on two independent experiments. Each $\text{IC}_{50}$ curve was determined based on at least 7 compound concentrations in triplicate. ND: not determined. NT: not tested.

To better understand the mechanism of Mpro inhibition by naphthoquinone-based derivatives, we evaluated whether compounds **382** and **415** were time-dependent inhibitors, a hallmark of covalent-acting molecules. First, enzyme inhibition after 15 min preincubation with the compounds was compared to activity without preincubation [130]. The IC_50_ values observed in these two assay conditions were similar, with slightly lower IC_50_ values upon preincubation (0.42 µM ± 0.02 upon incubation vs 0.80 µM ± 0.06 without incubation for **382** and 5.0 µM ± 0.2 upon incubation vs 16 µM ± 1 without incubation for **415**) (**Figure 6A** and **B**), while for the positive control GC373 the IC_50_ was ten-fold lower upon preincubation (0.003 µM ± 0.001). A dilution experiment was also performed, to check whether the compounds were irreversible. We incubated the inhibitors and Mpro at high concentrations and then diluted the incubation mixture, resulting in inhibitor concentrations 10-fold lower than their apparent IC_50_. In this assay, an irreversible inhibitor will maintain approximately 10% of enzymatic activity, while a rapidly reversible inhibitor will dissociate from the enzyme to restore approximately 90% of enzymatic activity following the dilution event [130, 131]. When this was performed with Mpro and GC373, a covalent Mpro inhibitor, the enzyme remained inhibited upon dilution. The same behavior was observed for compound **415** suggesting that this inhibitor is an irreversible covalent inhibitor (see **Figure 7** for the proposed binding mechanism). However, when the same test was carried out with compound **382** enzyme activity returned after dilution (**Figure 6C**). This suggested that the inhibition by **382** is reversible.

**Figure 6.**
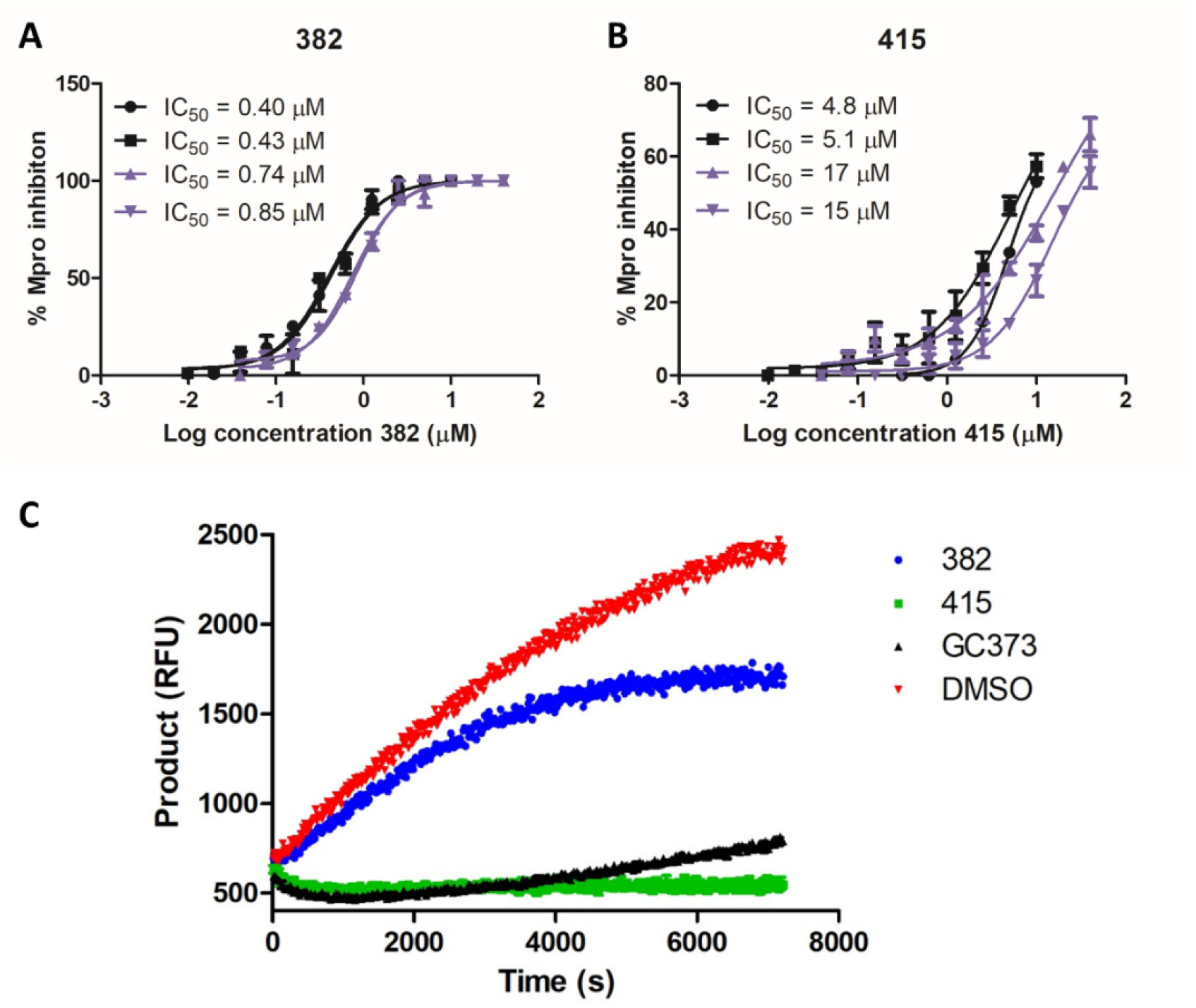
Evaluation of time-dependance and reversibility of Mpro inhibition by compounds **382** and **415**. IC_50_ curves for SARS-CoV-2 Mpro inhibitors **382** (A) and **415** (B). For each compound, two IC_50_ curves are shown, corresponding to two independent experiments (data shown as spheres or squares for each experiment), in which the compounds were pre-incubated with Mpro prior to substrate addition (black) and without preincubation with the compounds (purple). Reversibility assay (C). After preincubation of Mpro with compounds, at higher concentrations, the sample was diluted, and product formation was monitored for 120 minutes. Compound **382** reduced the enzymatic reaction rate by 26% compared to vehicle control (red), while the compound **415** reduced product formation by 100%, and this activity was not restored over a 2h period post dilution, as observed for the covalent inhibitor GC373 (black).

**Figure 7.**
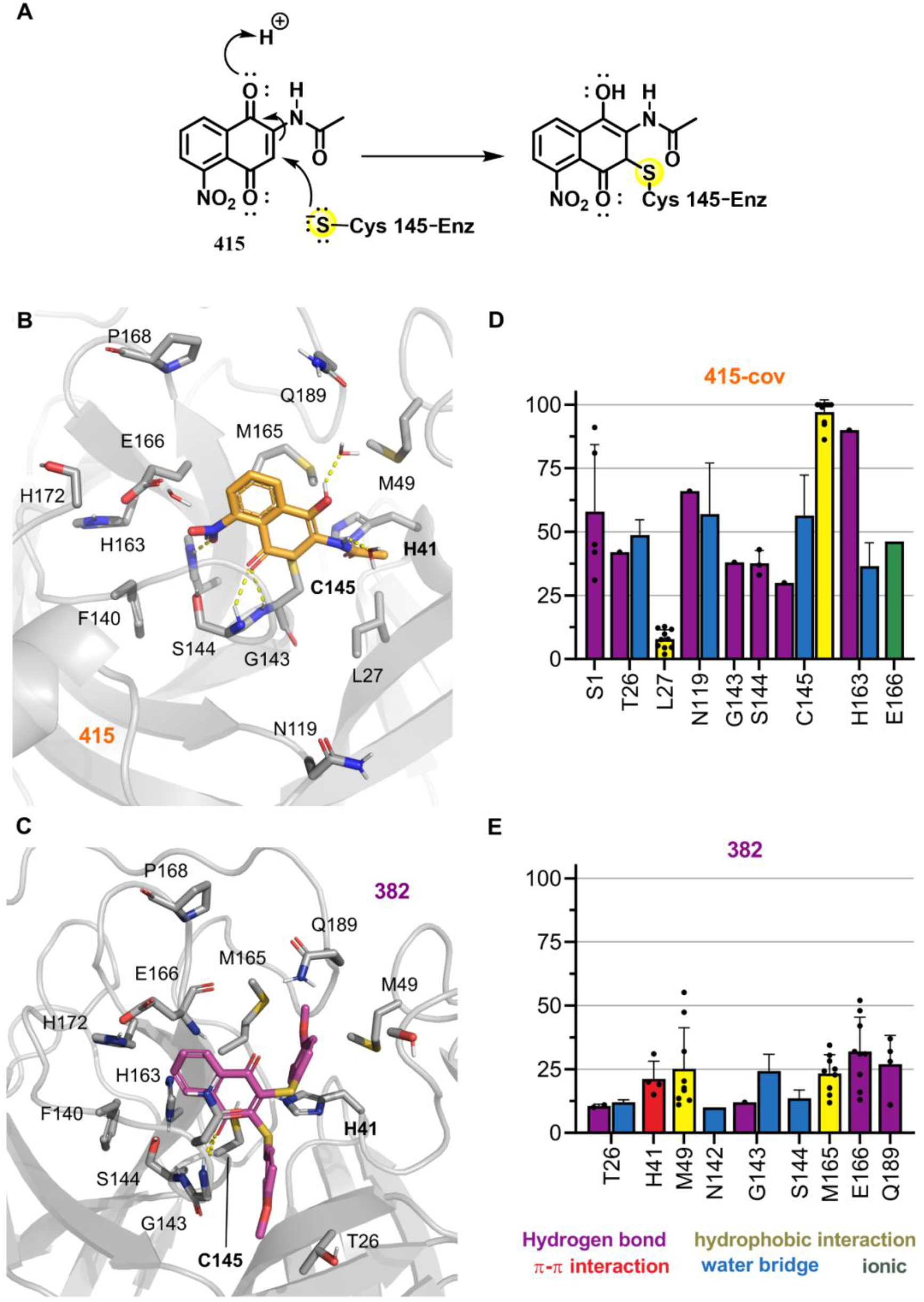
Predicted binding modes of compounds **415** and **382** to SARS-CoV-2 Mpro. The proposed mechanism of **415** covalent binding to C_145_ (A). Proposed binding modes from a representative frame in the MD simulation of compounds (B) **415** (orange) covalently bound to Mpro C_145_, and (C) **382** (pink) bound to Mpro, and the frequency of protein-ligand interactions for all simulations with ligands (D) **415** and (E) **382**. Mpro residues are colored according to the types of atoms in the interacting amino acid residues (protein carbon, light gray; nitrogen, blue; oxygen, red; sulfur, yellow), hydrogen bond interactions are represented as yellow dashed lines. Mean interaction frequency is represented, with standard error of the mean (N=5) interval depicted as error bars, each point displays the individual value for a particular simulation replica and each chain.

To gain insights into the proposed binding mode of our Mpro inhibitors and guide future optimization efforts, we conducted docking and MD studies with compounds **382** and **415,** representatives from two inhibitor scaffolds discovered. Our simulations considered the **415** ligand covalently bound, given the proposed reaction mechanism (**Figure 7A**), to both monomers in the Mpro dimer, and **382** freely. For the free simulations, however, the loss of interactions with E_166_ resulted in ligands being expelled from one of the binding sites within a few nanoseconds (∼200 ns) of simulation (Supporting Information **Table S2**). Our analysis is focused on the other binding site, that retained the ligand with stable interactions along the analyzed trajectory.

For both ligands, the most representative binding modes observed in the MD simulations (**Figure 7A** and **7B**) retain key interactions proposed based on docking with Glide (Supporting Information **Figure S45**). However, compound **415** showed a more conserved binding mode throughout the trajectory, being well represented by a single pose, in which the nitro group interacts with the S1 pocket and the 1,4 naphthoquinone interacts with the catalytic H_41_ and S2 subsite residues (**Figure 7B**). On the other hand, the higher variability in the orientation of compound **382** led to four clusters with frequency between 17.5 and 31.7% (Supporting Information **Figure S48**). Overall, the 1,4-naphthoquinone ring of ligand **415** occupies the S1 pocket, but fluctuations in the ring orientation reflect on varied positions for the phenyl substituents. In the most populated cluster (**Figure 7C**), the methoxyphenyl substituents occupy the S1ʹ and S2 subsites.

Compounds **382**, and **415** display stable polar contacts (hydrogen bond and water bridges) with G_143_ and S_144_ in the S1 pocket and π-cation or π-π interactions with the sidechain of H_41_. These interactions were more frequent in the covalent simulations. The ligands also display stable polar interactions with the main-chain nitrogen from E_166_ and electrostatic contacts with its side-chain (**Figure 7D** and **7E**), a residue that adopts a stable conformation due to an interaction between its sidechain and the S_1_ from the other protomer (S_1_*). Hydrophobic interactions to M_49_ and M_165_ from the S2 pocket, are seldomly observed for these inhibitors and frequent interactions with the side-chain of C_145_ was seen for the covalent inhibitor.

### 2.6. Validation of novel PLpro inhibitors

Although our virtual screening studies were focused solely on Mpro, we were also interested in testing the virtual screening hits against the second SARS-CoV-2 cysteine protease, PLpro, to determine if any of the molecules were dual inhibitors of the viral enzymes. PLpro cleaves three sites on the viral polypeptide but also acts as a de-ubiquitinase. Therefore, we identified a fluorogenic substrate for human de-ubiquitinases (Z-RLRGG-7-amino-4-methylcoumarin) as a substrate for SAR2-CoV-2 PLpro. Recombinant PLpro was incubated with 6 µM to 500 µM of this substrate and the K_M_ value was calculated to be 376.6 ± 32.3 µM. PLpro enzyme was pre-incubated with the same set of 23 compounds at 10 µM and then assayed with the fluorogenic substrate. Compound **668** was again eliminated due to insolubility in the assay buffer. Surprisingly, a total of 12 compounds inhibited PLpro by > 50% and the top three (compounds **159**, **189**, and **191**) inhibited at > 90%. These top compounds are *ortho*-quinone-based 1,2,3-triazoles derivatives, sharing a common scaffold. The IC_50_ values were calculated to be 1.7 µM, 2.2 µM, and 3.1 µM for compounds **159**, **189**, and **191**, respectively (**Figure 8**). Among the compounds that caused lower PLpro inhibition, five are N-substituted analogs of these hits, compounds **193-197**, and had IC_50_ values between 20 and 46 µM (**Table 1**, **Figure 7**). These eight PLpro inhibitors share a tricyclic 1,2-naphthoquinone ring that seems important for enzyme inhibition, as its replacement by a *para*-tolyl sulfone abolished activity against PLpro (compare compounds **189** vs **319**; **191** vs **321**; **193** vs **314**; and **197** vs **318**, **Table 1**).

**Figure 8.**
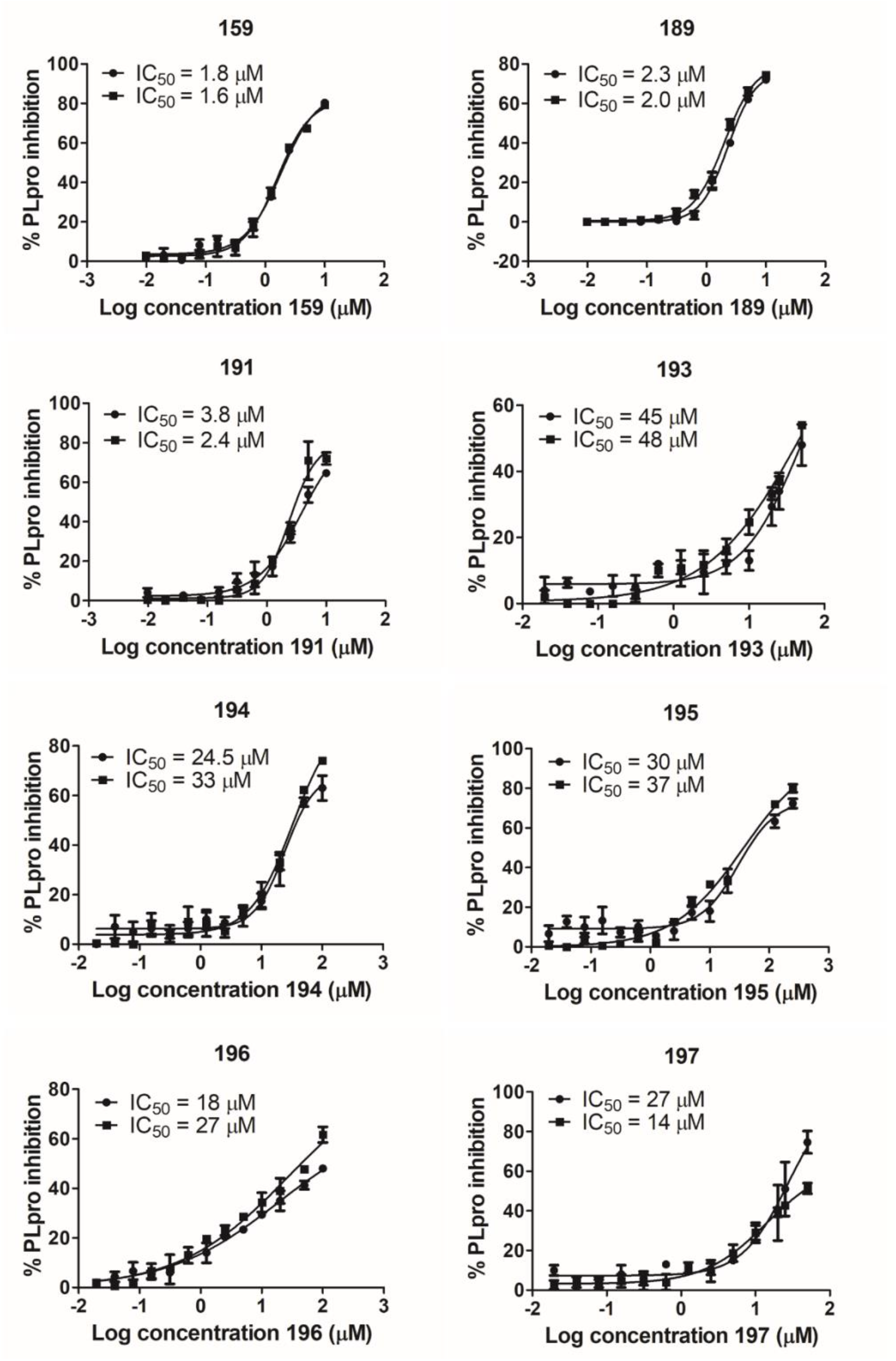
IC_50_ curves for SARS-CoV-2 PLpro inhibitors. For each compound, two IC_50_ curves are shown, corresponding to two independent experiments (data shown as spheres or squares for each experiment). Each curve was determined based on at least 11 compound concentrations in triplicate.

Since the PLpro inhibitors have a shared scaffold, we selected compounds **189** and **195** (N-substituted) for computational studies. The SARS-CoV-2 PLpro has similar folding to the homologous enzymes from other coronaviruses [132], with domains showing a “thumb-palm-fingers” pattern and an N-terminal ubiquitin-like (Ubl) domain (first 60 residues) (**Figure 9A**) [40]. As a cysteine protease, PLpro contains a canonical catalytic triad, Cys-His-Asp (C_111_, H_272_, and D_286_) [10] located in a solvent-exposed cleft at the interface of the palm and thumb domains [40]. Analysis of common protein-ligand interactions from the crystallographic structures showed little or no interaction with the catalytic triad, in agreement with very narrow S1 and S2 pockets, which have high specificity for glycine (**Figure 9B** and Supporting Information **Figure S49**). Only covalently bound peptidic inhibitors, containing glycines at P1 and P2, occupy these pockets [37, 40]. Instead, the non-covalent ligands bind to a groove corresponding to the S3 and S4 subsites, approximately 8 Å from the catalytic cysteine [37]. This groove is created due to the blocking loop 2 (BL2 loop), a flexible substrate-binding loop (Gly_266_-Gly_271_) found adjacent to the active site (**Figure 9A** and **B**). The BL2 loop is found in an open conformation in unbound PLpro, while it closes upon substrate or inhibitor binding [37].

**Figure 9.**
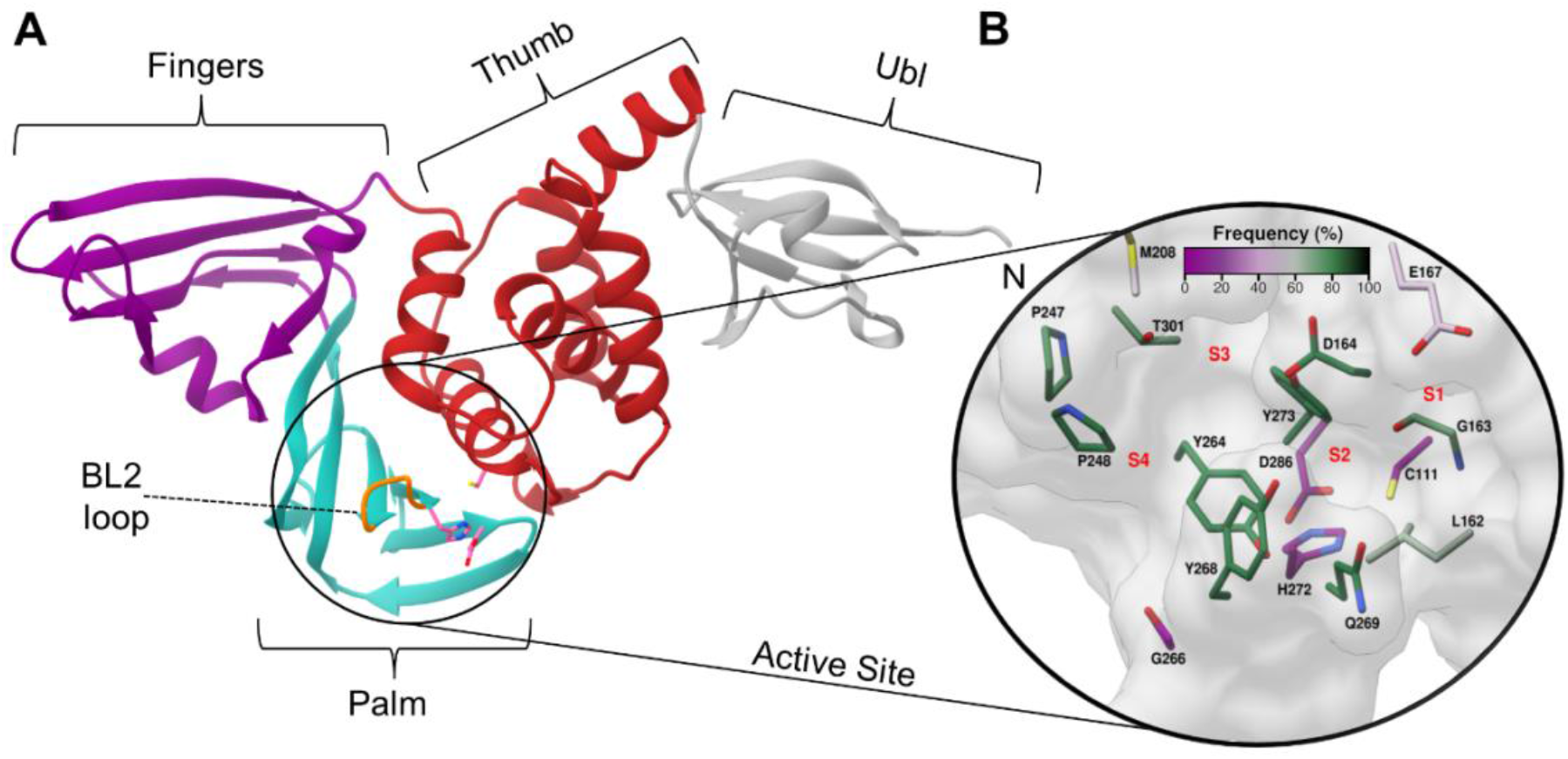
PLpro three-dimensional structure (PDB code: 7LBR [40]) and surface view of the active site. The four domains, fingers (purple), palm (green), thumb (red), and Ubl (gray) are showed in the cartoon representation (A). The BL2 loop (orange, Gly_266_-Gly_271_) is indicated by a line. The substrate binding cleft is highlighted with the catalytic triad C_111_, H_272_, and D_286_ displayed as sticks (pink) (B). In the close-up view of the active site, the residues are colored by the frequency they are involved in interactions with 21 crystallography ligands, according to an analysis using the program LUNA.

As observed for the crystallography ligands, compounds **189** and **195** showed docking predicted binding modes occupying the S3 and S4 subsites (**Figures 9**, **10** and Supporting Information **S49**). To verify the stability of these proposed binding modes, compounds **189** and **195** underwent MD simulations. The XR8-89 ligand (PDB code 7LBR [40]) was used as a positive control.

**Figure 10.**
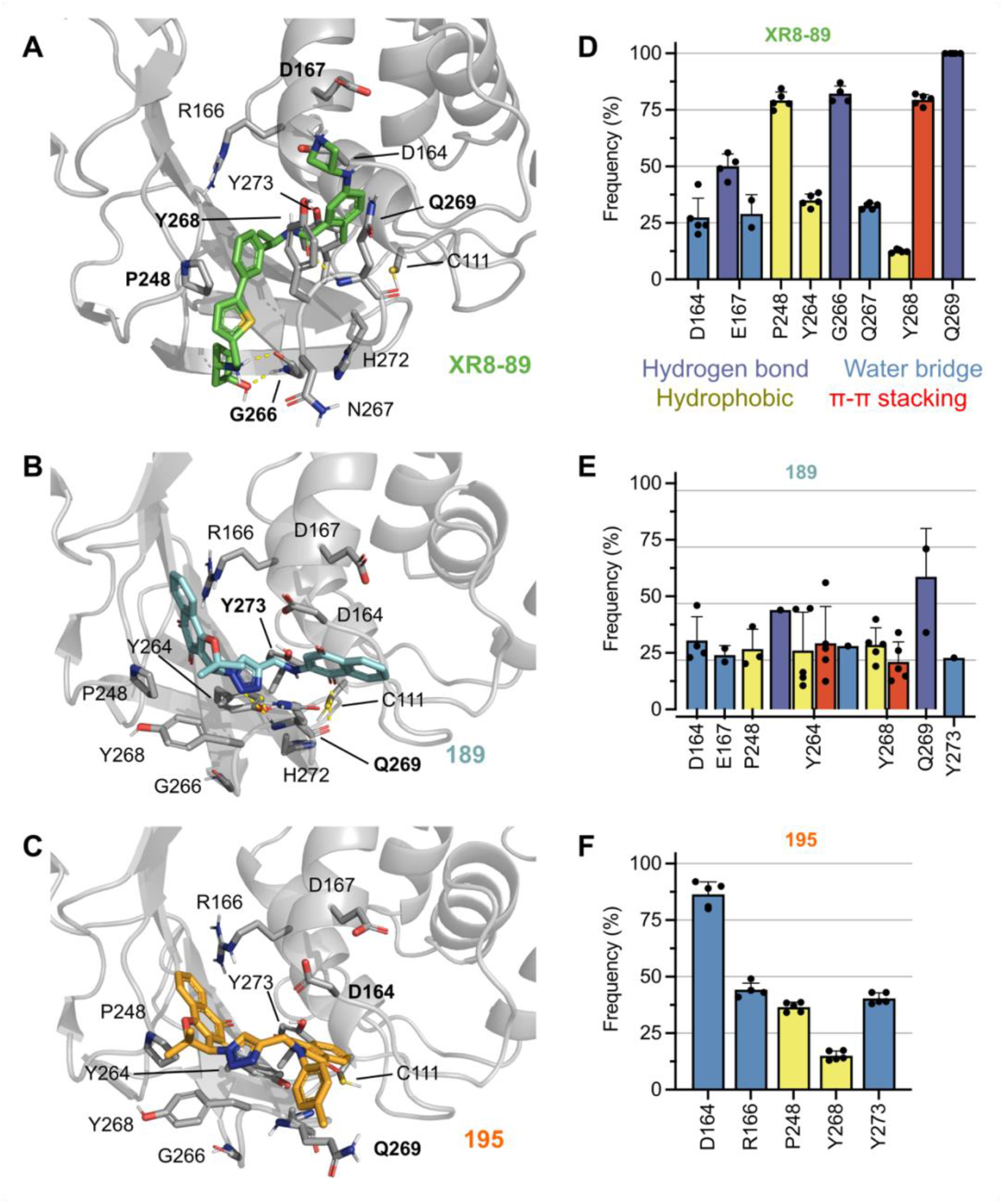
Proposed binding modes and protein-ligand interactions profile for PLpro inhibitors. Representative frames from the MD simulation describing the potential binding mode (A-C) for compounds **XR8-89** (green, from PDB code 7LBR [40]), **189** (cyan) and **195** (orange) bound to PLpro. PLpro residues are colored according to the types of atoms in the interacting amino acid residues (protein carbon, light gray; nitrogen, blue; oxygen, red; sulfur, yellow), hydrogen bond interactions are represented as yellow dashed lines. Frequency of protein-ligand interactions for all simulations with ligands **XR8-89** (D), **189** (E) and **195** (F). Mean interaction frequency is represented, with standard error of the mean (N=5) interval depicted as error bars, each point displays the individual value for a particular simulation replica.

In simulations with XR8-89, the BL2 loop remained in the closed conformation, and the ligand binding mode remained stable in all simulations, with its core structure being stabilized by hydrogen bond interactions between the carbonyl group and the backbone of Q_269_ (100% of the analyzed simulation time), as well as π-stacking interactions with Y_268_ (79% of the analyzed simulation time) (**Figures 10** and Supporting Information **S50**). We also observed a water bridge between the nitrogen on the amide and D_164_ (subsite S3, present on average 22% of the analyzed trajectory).

For **189**, four of the five simulated replicas showed stable interactions, with the initial pose changing dramatically from the initial coordinates after 500 ns in one of the replicas. In terms of binding mode, the triazole and central amine groups of **189** stablished hydrogen bond interactions with the Q_269_ (35% of the analyzed trajectory) and π-based interactions with Y_264_. The carbonyl groups from the naphthalene-1,4-dione moiety displayed water-mediated interactions with D_164_. The 1,2-naphthoquinone ring, shown to be essential for protease inhibition in our biochemical assays, binds to the S4 pocket, establishing hydrophobic interactions to P_248_ (**Figure 10B**).

In contrast to the observed for XR8-89 and **189**, the tolyl substituent on the amine of **195** prevented stable simulations on BL2 closed conformation. Thus, we performed 1 μs simulation initially, which displayed at first few interactions with the sidechain of Q_269_ (less than 20% of simulation time) and later stable interactions with D_164_ (over 66% of simulation time), while leading to the opening of BL2 and accommodating of the ligand. The last frame of this simulation was used to generate a further five new replicas (5 x 500 ns), to analyze the stability of this new binding mode, which was shown to be stabilized by water bridges with the D_164_ (>75%), R_166_ (>40%) and Y_273_ (∼ 40%) and hydrophobic contacts with P_248_ (>40%) (**Figure 10**).

### 2.7. Evaluation of hit compounds in a SARS-CoV-2 viral infection assay

We evaluated two Mpro hit compounds **382** and **415** and two PLpro hit compounds **189** and **191** in a SARS-CoV-2 viral infection assay of monkey-derived Vero E6 cells. The clinically approved RNA-polymerase inhibitor, remdesivir was used as a positive control. Remdesivir displayed antiviral efficacy in Vero E6 cells with EC_50_ of 2.45 µM and no host cell toxicity at concentrations up to 20 µM. Under the same culture conditions the hit naphthoquinone compounds were tested in three concentrations (24 µM, 6 µM, and 1.4 µM) and showed no significant antiviral activity dissociated from host cytotoxicity (Supporting Information **Figure S51**). We decided to test some compounds in serial dilution starting at 1 µM and in infected human-derived HeLa cells expressing ACE2 in addition to infected Vero cells. For HeLa-ACE2 cells, remdesivir was more potent with EC_50_ of 40 nM, however, cell cytotoxicity was also noted at concentrations above 2.4 µM. At lower concentrations the naphthoquinone compounds had no significant antiviral activity up to 1 µM (Supporting Information **Figure S52**). Therefore, it is important to further study the mechanism of action to understand the cytotoxicity and decouple from the direct antiviral activity.

## 3. Discussion

In this study, we used computational and biochemical approaches to evaluate a library of quinones to find inhibitors against SARS-CoV-2 proteases. The wealth of structural information from Mpro and PLpro allowed us to generate patterns of common protein-ligand interactions, which were helpful in two stages of our computational analysis. First, the selection of computational hits was guided by protein-ligand interactions frequently observed in Mpro crystallographic complexes. Thus, we prioritized compounds that interacted with conserved water molecules, S1 and S2 residues, filling one or more of the subsites with minimum solvent exposure. This strategy was successful as, among 24 compounds selected for inhibitory assays, three molecules with two different scaffolds were confirmed as Mpro inhibitors with low micromolar or submicromolar potency (compounds **382**, IC_50_ of 0.41 µM; **379**, IC_50_ of 0.63 µM; and **415**, IC_50_ of 5.0 µM), and another four compounds inhibited the enzyme by more than 50% at 10 μM. Additionally, for Mpro and PLpro inhibitors that were evaluated experimentally, we conducted MD simulations of the protein-ligands complexes. Together with the observed stability of binding poses during the simulations, the fact that our inhibitors establish interaction patterns commonly observed in the crystal structures encourages the application of our results in structure-based optimization projects.

During the validation of Mpro and PLpro inhibitors, several precautions were taken to avoid artifactual inhibition. We were especially careful considering previous reports that indicate quinones as potential Pan Assay Interference Compounds (PAINS) [133]. To avoid common causes of artifacts [128–130], we conducted the assays in the presence of detergent, avoiding compound aggregation, and verified that compounds were not highly fluorescent. In addition, a comparison of the inhibition of both target enzymes by each of the compounds indicates that all inhibitors showed specificity to one of our targets, reducing the likelihood they would be promiscuous inhibitors. Furthermore, to assess if Mpro inhibitors were time-dependent and/or irreversible inhibitors, we determined IC_50_ values of compounds **382** and **415** upon or without preincubation with the enzyme, and evaluated recovery of enzyme activity in a reversibility assays. Our results indicated that compound **382** is a reversible Mpro inhibitor, while **415** binds irreversibly to the target. This information was taken into account in our MD simulations, in which compound **415** was covalently bound to the enzyme, while compound **382** was simulated in a noncovalent complex.

An interesting pattern emerged in our MD simulations with Mpro complexes. For both compounds **382** and **415**, the simulations suggested stability of the complexes via multiple intermolecular interactions, with H_41_, G_143_, and E_166_. All three residues have reported key roles in the active site. As part of the catalytic dyad, H_41_ serves as a base for nucleophilic attack performed by C_145_ in peptide-bond cleavage [134], while G_143_, an oxyanion hole residue, helps stabilize the tetrahedral intermediate of the peptide-bond cleavage [135]. Moreover, E_166_ is essential for dimerization and its interactions with the other protomer N-finger also aid the correct orientation of H_163_ and H_172_ to form the S1 pocket [135, 136]. However, long-lasting interactions were observed only from one of the protomers’ binding sites, while the ligand bound to the other protomer was expelled within a few nanoseconds. The instability in one of the protomers was observed as a reproducible pattern in most replicates of our MD simulations. The complete deletion of the N-finger (residues S_1_* – R_4_*) in SARS-CoV Mpro, reduces the extent of the dimerization and completely abolishes the enzymatic activity (<1%) [137]. Simulations of Mpro from SARS-CoV-2 with peptidomimetic inhibitors or substrate [138], suggest that a similar mechanism exists, where the N-finger conformation upon dimerization exerts a direct influence on the oxyanion-loop motions and the stabilization of the catalytic conformation.

Our initial focus was on Mpro inhibition, however, we also tested the 23 soluble compounds selected against PLpro, to possibly find dual inhibitors for both SARS-CoV-2 viral enzymes. Despite the numerous efforts to develop inhibitors of the SARS-CoV-2 proteases, reports of dual Mpro/PLpro SARS-CoV-2 inhibitors are still scarce [139]. In the current study, the only dual inhibitor found was **191**, with modest Mpro inhibition (IC_50_ = 66 µM) and more potent PLpro inhibition (IC_50_ = 3.1 µM). Developing an effective dual inhibitor would require further optimization, but compound **191** is a candidate for such efforts. In addition, eight *ortho*-quinone-based 1,2,3, triazoles had IC_50_ < 50 µM against PLpro, including three inhibitors with IC_50_ in the single-digit micromolar range (compounds **159**, IC_50_ of 1.7 µM; **189**, IC_50_ of 2.2 µM; and **191**, IC_50_ of 3.1 µM). Considering the evidence supporting each SARS-CoV-2 protease as a therapeutic target [37,140,141], these compounds are interesting even in the absence of dual inhibition. MD simulations of **189** and **195** bound to PLpro were not as stable as the positive control XR8-89 (PDB code 7LBR [40]). The two scaffolds interacted, with low or moderate frequency, with residues in the S3 and S4 subsites, however, lacking long-lasting interactions with key residues, such as Y_268_ and Q_269_. These two residues form an unusual β-turn in the flexible β-hairpin BL2 loop that controls the access to the active site in the binding of host and viral proteins [40]. Thus, compound **189** and, particularly, **195** might not fully stabilize the closed conformation of BL2 loop as well as the potent XR8-89.

## 4. Conclusion

Here, we employed computational and biochemical assays to evaluate a quinones library, leading to the identification of 11 promising naphthoquinoidal inhibitors against the two SARS-CoV-2 viral proteases, Mpro and PLpro, with potency in the mid micromolar to nanomolar range. For all inhibitors experimentally characterized, we propose likely binding modes with good complementarity to the protease active sites, that closely resemble protein-ligand interaction patterns observed in crystallographic complexes and which were stable in MD simulations. Hence, the inhibitors presented here are novel scaffolds for further optimization to develop a treatment against SARS-CoV-2 infection.

## 5. Experimental Section

### 5.1. Compounds general experimental details

All chemicals were obtained from commercial sources and used without further purification. Melting points were obtained on a Thomas Hoover apparatus and are uncorrected. Column chromatography was performed on silica gel (Silica Flash G60 UltraPure 60-200 µm, 60 Å). Infrared spectra were recorded on a Shimadzu FTIR Spectrometer IR Prestige-21. ^1^H and ^13^C NMR were recorded at room temperature using a Bruker AVANCE DRX 200 and DRX 400 MHz, in the solvents indicated, with tetramethylsilane (TMS) as internal reference. Chemical shifts (δ) are given in parts per million (ppm) and coupling constants (J) in Hertz (Hz). The mass spectrometer was operated in the positive ion mode. A standard atmospheric pressure photoionization (APPI) source was used to generate the ions. The sample was injected using a constant flow (3 µL/min). The solvent was an acetonitrile/methanol mixture. The APPI-Q-TOF MS instrument was calibrated in the mass range of 50-3000 m/z using an internal calibration standard (low concentration tuning mix solution) supplied by Agilent Technologies. Data were processed employing Bruker Data Analysis software version 4.0. Compounds were named following IUPAC rules as applied by ChemBioDraw Ultra (version 12.0).

### 5.2. Synthesis of candidate inhibitors

*Ortho*-quinone-based 1,2,3-triazoles compounds **159**, **189**, **191-197**, were prepared according to previously reported reports and their data are consistent with the literature [80,82,91,142]. *Para*-quinones-based 1,2,3-triazoles compounds **314**, **318**-**321** were prepared as described in the literature [90]. *Para*-quinones and derivatives **379**, **380**, **382**, **414**, **415**, **465**, **470**, **477**, were synthesized following the previously published studies in the literature [76, 99]. Hydrazo derivatives **666**, **668** and **673** were prepared according to previously published reports and their data are consistent with the literature [68]. NMR spectra for all compounds have been previously published when they were originally described.

### 5.3. Comparison of available SARS-CoV-2 Mpro structures

All 72 crystallographic structures were downloaded from the PDB [143] (structures available in April/2020). Structural superposition was performed with program R [144]’s package, Bio3D [145], using the protein’s Cα. RMSD and PCA were also done with Bio3D package. As water molecules might play important roles in Mpro catalysis and ligand stabilization, we used the ProBiS H2O plugin [123]. This PyMol [146] plugin enables the identification of conserved water sites in proteins using experimental determined protein structures. The highest resolution structure, PDB code 5R82 [117], was used as reference to establish the water molecules position.

### 5.4. Analysis of protein-ligand interactions

The program LUNA (https://github.com/keiserlab/LUNA) was used to perform large-scale analysis of non-covalent interactions between the protein-ligands complexes of Mpro. With this program, it was possible identify frequently interacting residues between the ligands and Mpro active site. We submitted a list containing the of PDB ids of the 72 structures, discriminating chain A and the binding site ligands to be analyzed. After processing, we investigated the table (in .csv format) of the interacting frequencies by residues and ligands with program R.

### 5.5. Ligand and protein preparation

Three-dimensional ligand structures were generated with LigPrep (version 46013), using Epik to predict their protonation in pH 7.0 ± 2.0, and generating tautomers and diastereoisomers. The OPLS3e force-field was employed for structure generation. The SARS-CoV-2 Mpro protein structure was prepared from the PDB 5R82 [117], using the Protein Wizard Preparation tool, with standard options. Two Mpro receptor files were prepared for docking: one with all water removed and another containing waters 1189 and 1284 from the original PDB. The SARS-CoV-2 PLpro structure was prepared from the PDB 7LBR [40], using the same protocol as Mpro.

### 5.6. Molecular docking

Molecular docking was carried out with Glide SP (version 9.1) and Autodock Vina. For docking to Mpro with Glide [124], grids were centered at the central point of the active site residues G_143_, C_145_, M_49_ and H_41_ (coordinates: 10.7313390385, - 4.49000171154, 22.4985591538). Two docking grids were generated: one without waters and one containing the two conserved waters described in the protein preparation. Each compound was docked using both grids. In both cases, the dimensions of the inner box had 10 Å in each direction and the outer box had 30 Å in each direction. Whenever mentioned, covalent docking as performed using CovDock [147] using the C_145_ as anchor, nucleophilic addition to double bond as reaction type and generating up to 10 poses for each ligand. Poses were selected according to the docking score and relevant interactions.

For docking to Mpro with Autodock Vina [125], a grid box of size 22.5×24.5×22.5 Å was centralized in the geometrical center among the residues T_26_, M_49_, N_142_, and M_165_. All the experiments were done in triplicate starting from a random seed. Energy range, exhaustiveness, and the number of maximum modes parameters were set to 3 kcal/mol, 8, 9, respectively. Similar to docking using Glide, two experiments were done with and without conserved waters. For selected ligands, induced-fit docking was performed (with and without the conserved waters) by flexing the residues N_142_, E_189_, M_49_, and M_165_.

For docking to PLpro with Glide, using the Induced-Fit mode [148], a cubic grid box of size 12 Å was centralized in the geometric center of the co-crystallized ligand (PDB code 7LBR [40]).

### 5.7. Molecular Dynamics simulations

Prepared SARS-CoV-2 Mpro and PLpro structures were simulated with the selected ligands. MD simulations were carried out by using the Desmond engine [149] with the OPLS3e force-field [150] according to a previously described protocol [151]. In short, the system encompassed the protein-ligand/cofactor complex, a predefined water model (TIP3P [152]) as a solvent and counterions (Na^+^ or Cl^-^ adjusted to neutralize the overall system charge). The system was treated in a cubic box with a periodic boundary condition (PBC) specifying the shape and the size of the box as 13 Å distance from the box edges to any atom of the protein. Short-range coulombic interactions were calculated using 1 fs time steps and 9.0 Å cut-off value, whereas long-range coulombic interactions were estimated using the Smooth Particle Mesh Ewald (PME) method [153]. Each Mpro and PLpro systems were subjected to at least 5 μs simulations (five replicas of 1 μs each), with exception of PLpro – compound **195**, which had one preliminary 1 μs simulation, from which a stable conformation was selected for further shorter simulations. Atomic interactions and distances were determined using the Simulation Event Analysis pipeline as implemented in Maestro 2020.2 (Schrödinger LCC).

Representative frames of the simulations were retrieved from clustering, which was performed with hierarchical clustering analyses. Trajectories were clustered using the script trj_cluster.py (implemented in Maestro 2021.2, Schrödinger LCC) using 2 Å as cut-off, which was chosen upon evaluating the RMSD of ligand’s heavy atoms. Trajectories where the ligand was expelled of the pocket were not considered for clustering or interaction analyses.

RMSD values of the protein backbone were used to monitor simulation equilibration and protein folding changes (Supporting Information Figure S50). All the trajectory and interaction data are available on the Zenodo repository (code: 10.5281/zenodo.5147951). MD trajectories were visualized, and figures produced by PyMol v.2.4 (Schrödinger LCC, New York, NY, USA).

### 5.8. Synthesis of Mpro substrate

A quenched fluorogenic peptide substrate with the sequence (D-Arg)-(D-Arg)-Lys(MCA)-Ala-Thr-Leu-Gln-Ala-Ile-Ala-Ser-Lys(DNP)-COOH (ATLQAIAS) was synthesized on a Biotage Syroll peptide synthesizer at room temperature through fluorenylmethyloxycarbonyl (Fmoc) solid-phase synthesis. The synthesis scale was 12.5 μmole with preloaded lysine(2-dinitrophenyl) Wang resin, where the DNP quencher was linked to the epsilon nitrogen of the lysine. For each coupling reaction, 4.9 equivalents of HCTU (O-(1H-6-chlorobenzotriazole-1-yl)-1,1,3,3-tetramethyluronium hexafluoro-phosphate), 5 equivalents of Fmoc-amino acid-OH, and 20 equivalents of N-methylmorpholine (NMM) in 500 μL N,N-dimethylformamide (DMF) were used. The coupling reaction was carried out with shaking for 8 minutes. Each amino acid position was double coupled, and subsequent Fmoc deprotection was done with 500 μL of 40% 4-methylpiperidine in dimethyl formamide (DMF) for 10 minutes. Deprotection was followed by a wash with 500 μL of DMF for 3-minutes and the wash was repeated 6 times. The lysine amino acid, lysine (7-methoxycoumarin-4-acetic acid (MCA), was coupled where MCA was linked to the epsilon nitrogen of the lysine. The two final amino acid position couplings used d-Arginine to increase peptide solubility. The cleavage of the peptide from the Wang resin was carried out with a 500 μL of solution composed of 95% trifluoroacetic acid, 2.5% water, and 2.5% triisopropylsilane at room temperature for 1 hour with shaking. The crude peptide product was precipitated in 30 mL of a 1:1 mixture of cold diethyl ether and hexane. Product was then solubilized in a 1:1:1 mixture of DMSO, water and acetonitrile. The solubilized crude material was purified by high-performance liquid chromatography (HPLC) using an Agilent Pursuit 5 C18 column (5 mm bead size, 150 x 21.2 mm) on an Agilent PrepStar 218 series preparative HPLC. Mobile phase A was water + 0.1% TFA, and mobile phase B was acetonitrile + 0.1% TFA. The peptide product fractions were collected, combined, and had solvent removed under reduced atmosphere. The peptide substrate was solubilized in DMSO to a final concentration of 50 mM. Purity was confirmed by liquid chromatography-mass spectrometry and the stock was stored at −20°C.

### 5.9. Assays against Mpro

Recombinant SARS-CoV-2 Mpro was expressed and purified as described previously in Mellot et al. [48]. Mpro activity was measured using the fluorogenic substrate, ATLQAIAS, on a Biotek® Synergy HTX plate reader. All assays were performed in black flat-bottom 384-well plates, in 30 µL of 50 mM Tris-HCl pH 7.5, 150 mM NaCl, 1 mM EDTA, 0.01% Tween-20 using 50 nM Mpro and 10 µM of FRET substrate. Initial screening was performed at 10 µM. Prior to addition of the substrate, enzyme was incubated with the compounds for 15 minutes. Following the substrate addition proteolysis was measured at 320/420 nm (excitation/emission) at 25 °C. Percent inhibition was calculated relative to control reactions containing a maximum of 0.5% DMSO. Two independent experiments were performed in triplicate wells. Half-maximal inhibitory concentration (IC_50_) was determined by nonlinear regression analysis of the velocity vs. inhibitor concentration plot using GraphPad Prism 6 (GraphPad Prism, version 6.00, La Jolla, California, USA). At least seven inhibitor concentrations were used to build each curve. DMSO was used as negative control. The hit compounds **382** and **415** were also tested without incubation to investigate the time-dependency behavior.

### 5.10. Reversibility assay

Mpro at 100-fold its final assay concentration was incubated with the hits at 10-fold its respective IC_50_ value for 30 min in a volume of 2 μL. This mixture was diluted 100-fold with an assay buffer containing 10 μM ATLQAIAS substrate to a final volume of 30 μL, resulting in a standard concentration of Mpro and 0.1 times the IC_50_ value of hits [130, 131]. Fluorescence intensities were monitored continuously during substrate hydrolysis on Synergy 2 (BioTek^®^) plate reader for 120 minutes.

### 5.11. Assays against PLpro

Recombinant SARS-CoV-2 PLpro was purchased from Acro Biosystems, PAE-C5184. Proteolytic activity was measured using Z-Arg-Leu-Arg-Gly-Gly-AMC substrate (Bachem, 369 I1690) as described previously in Ashhurst et al. [154] The release of fluorescent 7-amido-4-metyhlcoumarin was measured at 360 nm/460 nm wavelengths for excitation/emission, on a Biotek® Synergy HTX. All assays were performed in 384-well black plate at 25 °C, in a final volume of 30 µL of 50 mM HEPES pH 6.5, 150 mM NaCl, 0.1 mM DTT, 0.01% Tween-20, 50 nM enzyme and 50 µM of substrate. Enzymatic activity was calculated by comparison to initial rates of reaction of a DMSO control. Initial screening was performed at 10 µM of each compound in triplicate wells. Compounds that inhibited by 75% or more of the PLpro activity in the initial screen had their IC_50_ determined. At least two independent experiments were performed, each involving at least eleven compound concentrations in triplicates. IC_50_ curves were obtained by non-linear regressions analysis of the velocity vs. inhibitor concentration using GraphPad Prism 6 (GraphPad Prism, version 6.00, La Jolla, California, USA). Reported IC_50_ values refer to the mean values and the standard error of the mean.

### 5.12. Antiviral activity

Antiviral assays were performed according to the protocol previously described in Mellot et al. [48]. Compounds **189**, **191**, **382**, and **415** were evaluated in a SARS-CoV-2 viral infection assay of monkey-derived Vero E6 cells and human-derived HeLa cells that overexpress ACE2. Remdesivir was employed as a positive control. Each compound was evaluated in ten concentrations, in two-fold dilutions, from 20 µM to 39 nM in the case of remdesivir and from 1.0 µM to 1.9 nM for all other compounds, in triplicates.

## 6. Author Contributions

L.H.S. analyzed PDB structures and interaction patterns; L.H.S., R.E.O.R., and R.S.F performed and analyzed virtual screening results; T.K. and A.P. performed and analyzed molecular dynamics simulations; R.G.A. and E.N.S.J. designed the chemical library for virtual screening; C.B., A.L.L. and C.S.C. designed and synthesized the Mpro substrate. D.S., E.B.S. and A.J.O. performed Mpro and PLpro assays; J.C.O., L.V.B. and T.K. performed docking studies; P.F., D.S., L.M.P., J.H.M. and A.J.O. expressed and purified Mpro; M.A.G., B.W., and J.L.S.N. performed antiviral assays. All authors were involved in experiment design and analyses; L.H.S., T.K., R.G.A., E.B.S., E.N.S.J., and R.S.F wrote the manuscript, with revisions and contributions from all authors. E.N.S.J. and R.S.F conceived the overall design of the study.

## 8. Acknowledgments

The authors would like to thank CNPq, CAPES (Finance Code 001) and FAPEMIG for the financial support and scholarships. E. N. da Silva Júnior acknowledges funding from CNPq (PQ 309774/2020-9), Fapemig (Rede de Pesquisa e Inovação para Bioengenharia de Nanossistemas-RED-00282-16 and PPM-00635-18), Return Fellowship of the Alexander von Humboldt Foundation (AvH) and the Royal Society of Chemistry for the research fund grant (R19-9781). R.S.F. received funding from CAPES (grant CAPES-EPIDEMIAS-0688/2020), FAPEMIG (Rede Mineira de Imunobiologicos grant # REDE-00140-16) and holds a CNPq Researcher Scholarship (Bolsa de Produtividade em Pesquisa, 306606/2017-8). J.C.O. and L.V.B. received scholarships from CAPES (processes 88887.508402/2020-00 and 88887.518393/2020-00, grant CAPES-EPIDEMIAS - Programa Estratégico Emergencial de Prevenção e Combate a Surtos, Endemias, Epidemias e Pandemias). C.B., A.L.L. and C.S.C. were supported by National Institutes of Health grant P50AI150476. The SARS-CoV-2 Mpro plasmid was provided by Rolf Hilgenfeld, University of Lübeck, Germany. Authors would also like to thank the CSC-Finland for the generous computational resources provided.

## 9. Appendix A. Supplementary data

Supplementary Information that contains general information of the assembly of the chemical library, structural information of all virtual screened compounds, detailed information of the computational procedures is available for this manuscript, and SARS-CoV-2 viral infection assay.

## Abbreviations

SARS: severe acute respiratory syndrome
MERS: Middle East Respiratory Syndrome
Mpro: SARS-CoV-2 Main protease
PLpro: SARS-CoV-2 Papain-like protease
PCA: Principal Component Analysis
Hbond: Hydrogen bond
IC_50_: Half-maximal inhibitory concentration
MD: Molecular Dynamics
BL2: Blocking Loop 2
RMSD: Root-Mean-Square Deviation

## References

[1] A.E. Gorbalenya, S.C. Baker, R.S. Baric, R.J. de Groot, C. Drosten, A.A. Gulyaeva, B.L. Haagmans, C. Lauber, A.M. Leontovich, B.W. Neuman, D. Penzar, S. Perlman, L.L.M. Poon, D. V. Samborskiy, I.A. Sidorov, I. Sola, J. Ziebuhr, The species Severe acute respiratory syndrome-related coronavirus: classifying 2019-nCoV and naming it SARS-CoV-2, Nat. Microbiol. 5 (2020) 536–544. https://doi.org/10.1038/s41564-020-0695-z.

[2] P. Zhou, X. Lou Yang, X.G. Wang, B. Hu, L. Zhang, W. Zhang, H.R. Si, Y. Zhu, B. Li, C.L. Huang, H.D. Chen, J. Chen, Y. Luo, H. Guo, R. Di Jiang, M.Q. Liu, Y. Chen, X.R. Shen, X. Wang, X.S. Zheng, K. Zhao, Q.J. Chen, F. Deng, L.L. Liu, B. Yan, F.X. Zhan, Y.Y. Wang, G.F. Xiao, Z.L. Shi, A pneumonia outbreak associated with a new coronavirus of probable bat origin, Nature. 579 (2020) 270–273. https://doi.org/10.1038/s41586-020-2012-7.

[3] L.T. Phan, T. V. Nguyen, Q.C. Luong, T. V. Nguyen, H.T. Nguyen, H.Q. Le, T.T. Nguyen, T.M. Cao, Q.D. Pham, Importation and Human-to-Human Transmission of a Novel Coronavirus in Vietnam, N. Engl. J. Med. 382 (2020) 872–874. https://doi.org/10.1056/nejmc2001272.

[4] M.A. Shereen, S. Khan, A. Kazmi, N. Bashir, R. Siddique, COVID-19 infection: Origin, transmission, and characteristics of human coronaviruses, J. Adv. Res. 24 (2020) 91–98. https://doi.org/10.1016/j.jare.2020.03.005.

[5] J. Liu, X. Zheng, Q. Tong, W. Li, B. Wang, K. Sutter, M. Trilling, M. Lu, U. Dittmer, D. Yang, Overlapping and discrete aspects of the pathology and pathogenesis of the emerging human pathogenic coronaviruses SARS-CoV, MERS-CoV, and 2019-nCoV, J. Med. Virol. 92 (2020) 491–494. https://doi.org/10.1002/jmv.25709.

[6] N. Zhu, D. Zhang, W. Wang, X. Li, B. Yang, J. Song, X. Zhao, B. Huang, W. Shi, R. Lu, P. Niu, F. Zhan, X. Ma, D. Wang, W. Xu, G. Wu, G.F. Gao, W. Tan, A Novel Coronavirus from Patients with Pneumonia in China, 2019, N. Engl. J. Med. 382 (2020) 727–733. https://doi.org/10.1056/nejmoa2001017.

[7] N. Drayman, J.K. DeMarco, K.A. Jones, S.-A. Azizi, H.M. Froggatt, K. Tan, N.I. Maltseva, S. Chen, V. Nicolaescu, S. Dvorkin, K. Furlong, R.S. Kathayat, M.R. Firpo, V. Mastrodomenico, E.A. Bruce, M.M. Schmidt, R. Jedrzejczak, M.Á. Muñoz-Alía, B. Schuster, V. Nair, K. Han, A. O’Brien, A. Tomatsidou, B. Meyer, M. Vignuzzi, D. Missiakas, J.W. Botten, C.B. Brooke, H. Lee, S.C. Baker, B.C. Mounce, N.S. Heaton, W.E. Severson, K.E. Palmer, B.C. Dickinson, A. Joachimiak, G. Randall, S. Tay, Masitinib is a broad coronavirus 3CL inhibitor that blocks replication of SARS-CoV-2, Science (80-.). (2021) eabg5827. https://doi.org/10.1126/science.abg5827.

[8] W. Rut, Z. Lv, M. Zmudzinski, S. Patchett, D. Nayak, S.J. Snipas, F. El Oualid, T.T. Huang, M. Bekes, M. Drag, S.K. Olsen, Activity profiling and crystal structures of inhibitor-bound SARS-CoV-2 papain-like protease: A framework for anti–COVID-19 drug design, Sci. Adv. 6 (2020) eabd4596. https://doi.org/10.1126/sciadv.abd4596.

[9] S. Ullrich, C. Nitsche, The SARS-CoV-2 main protease as drug target, Bioorganic Med. Chem. Lett. 30 (2020) 127377. https://doi.org/10.1016/j.bmcl.2020.127377.

[10] Y.M. Báez-Santos, S.E. St. John, A.D. Mesecar, The SARS-coronavirus papain-like protease: Structure, function and inhibition by designed antiviral compounds, Antiviral Res. 115 (2015) 21–38. https://doi.org/10.1016/j.antiviral.2014.12.015.

[11] C. Wu, Y. Liu, Y. Yang, P. Zhang, W. Zhong, Y. Wang, Q. Wang, Y. Xu, M. Li, X. Li, M. Zheng, L. Chen, H. Li, Analysis of therapeutic targets for SARS-CoV-2 and discovery of potential drugs by computational methods, Acta Pharm. Sin. B. 10 (2020) 766–788. https://doi.org/10.1016/j.apsb.2020.02.008.

[12] Y. Kim, H. Liu, A.C. Galasiti Kankanamalage, S. Weerasekara, D.H. Hua, W.C. Groutas, K.O. Chang, N.C. Pedersen, Reversal of the Progression of Fatal Coronavirus Infection in Cats by a Broad-Spectrum Coronavirus Protease Inhibitor, PLoS Pathog. 12 (2016) e1005531. https://doi.org/10.1371/journal.ppat.1005531.

[13] L. Zhang, D. Lin, Y. Kusov, Y. Nian, Q. Ma, J. Wang, A. Von Brunn, P. Leyssen, K. Lanko, J. Neyts, A. De Wilde, E.J. Snijder, H. Liu, R. Hilgenfeld, α-Ketoamides as Broad-Spectrum Inhibitors of Coronavirus and Enterovirus Replication: Structure-Based Design, Synthesis, and Activity Assessment, J. Med. Chem. 63 (2020) 4562–4578. https://doi.org/10.1021/acs.jmedchem.9b01828.

[14] H. Yang, W. Xie, X. Xue, K. Yang, J. Ma, W. Liang, Q. Zhao, Z. Zhou, D. Pei, J. Ziebuhr, R. Hilgenfeld, Y.Y. Kwok, L. Wong, G. Gao, S. Chen, Z. Chen, D. Ma, M. Bartlam, Z. Rao, Design of wide-spectrum inhibitors targeting coronavirus main proteases, PLoS Biol. 3 (2005) e324. https://doi.org/10.1371/journal.pbio.0030324.

[15] K. Anand, G.J. Palm, J.R. Mesters, S.G. Siddell, J. Ziebuhr, R. Hilgenfeld, Structure of coronavirus main proteinase reveals combination of a chymotrypsin fold with an extra α-helical domain, EMBO J. 21 (2002) 3213–3224. https://doi.org/10.1093/emboj/cdf327.

[16] H. Yang, M. Yang, Y. Ding, Y. Liu, Z. Lou, Z. Zhou, L. Sun, L. Mo, S. Ye, H. Pang, G.F. Gao, K. Anand, M. Bartlam, R. Hilgenfeld, Z. Rao, The crystal structures of severe acute respiratory syndrome virus main protease and its complex with an inhibitor, Proc. Natl. Acad. Sci. U. S. A. 100 (2003) 13190– 13195. https://doi.org/10.1073/pnas.1835675100.

[17] X. Xue, H. Yu, H. Yang, F. Xue, Z. Wu, W. Shen, J. Li, Z. Zhou, Y. Ding, Q. Zhao, X.C. Zhang, M. Liao, M. Bartlam, Z. Rao, Structures of Two Coronavirus Main Proteases: Implications for Substrate Binding and Antiviral Drug Design, J. Virol. 82 (2008) 2515–2527. https://doi.org/10.1128/jvi.02114-07.

[18] T. Pillaiyar, M. Manickam, V. Namasivayam, Y. Hayashi, S.H. Jung, An overview of severe acute respiratory syndrome-coronavirus (SARS-CoV) 3CL protease inhibitors: Peptidomimetics and small molecule chemotherapy, J. Med. Chem. 59 (2016) 6595–6628. https://doi.org/10.1021/acs.jmedchem.5b01461.

[19] W. Dai, B. Zhang, X.M. Jiang, H. Su, J. Li, Y. Zhao, X. Xie, Z. Jin, J. Peng, F. Liu, C. Li, Y. Li, F. Bai, H. Wang, X. Cheng, X. Cen, S. Hu, X. Yang, J. Wang, X. Liu, G. Xiao, H. Jiang, Z. Rao, L.K. Zhang, Y. Xu, H. Yang, H. Liu, Structure-based design of antiviral drug candidates targeting the SARS-CoV-2 main protease, Science (80-.). 368 (2020) 1331–1335. https://doi.org/10.1126/science.abb4489.

[20] M.D. Sacco, C. Ma, P. Lagarias, A. Gao, J.A. Townsend, X. Meng, P. Dube, X. Zhang, Y. Hu, N. Kitamura, B. Hurst, B. Tarbet, M.T. Marty, A. Kolocouris, Y. Xiang, Y. Chen, J. Wang, Structure and inhibition of the SARS-CoV-2 main protease reveal strategy for developing dual inhibitors against Mpro and cathepsin L, Sci. Adv. 6 (2020) eabe0751. https://doi.org/10.1126/sciadv.abe0751.

[21] H.C. Hung, Y.Y. Ke, S.Y. Huang, P.N. Huang, Y.A. Kung, T.Y. Chang, K.J. Yen, T.T. Peng, S.E. Chang, C.T. Huang, Y.R. Tsai, S.H. Wu, S.J. Lee, J.H. Lin, B.S. Liu, W.C. Sung, S.R. Shih, C.T. Chen, J.T.A. Hsu, Discovery of M protease inhibitors encoded by SARS-CoV-2, Antimicrob. Agents Chemother. 64 (2020). https://doi.org/10.1128/AAC.00872-20.

[22] L. Zhang, D. Lin, X. Sun, U. Curth, C. Drosten, L. Sauerhering, S. Becker, K. Rox, R. Hilgenfeld, Crystal structure of SARS-CoV-2 main protease provides a basis for design of improved a-ketoamide inhibitors, Science (80-.). 368 (2020) 409–412. https://doi.org/10.1126/science.abb3405.

[23] C. Ma, M.D. Sacco, B. Hurst, J.A. Townsend, Y. Hu, T. Szeto, X. Zhang, B. Tarbet, M.T. Marty, Y. Chen, J. Wang, Boceprevir, GC-376, and calpain inhibitors II, XII inhibit SARS-CoV-2 viral replication by targeting the viral main protease, Cell Res. 30 (2020) 678–692. https://doi.org/10.1038/s41422-020-0356-z.

[24] Z. Jin, X. Du, Y. Xu, Y. Deng, M. Liu, Y. Zhao, B. Zhang, X. Li, L. Zhang, C. Peng, Structure of M pro from SARS-CoV-2 and discovery of its inhibitors, Nature. 582 (2020) 289–293.

[25] C.-H. Zhang, E.A. Stone, M. Deshmukh, J.A. Ippolito, M.M. Ghahremanpour, J. Tirado-Rives, K.A. Spasov, S. Zhang, Y. Takeo, S.N. Kudalkar, Z. Liang, F. Isaacs, B. Lindenbach, S.J. Miller, K.S. Anderson, W.L. Jorgensen, Potent Noncovalent Inhibitors of the Main Protease of SARS-CoV-2 from Molecular Sculpting of the Drug Perampanel Guided by Free Energy Perturbation Calculations, ACS Cent. Sci. 7 (2021) 467–475. https://doi.org/10.1021/acscentsci.1c00039.

[26] H. Su, S. Yao, W. Zhao, Y. Zhang, J. Liu, Q. Shao, Q. Wang, M. Li, H. Xie, W. Shang, C. Ke, L. Feng, X. Jiang, J. Shen, G. Xiao, H. Jiang, L. Zhang, Y. Ye, Y. Xu, Identification of pyrogallol as a warhead in design of covalent inhibitors for the SARS-CoV-2 3CL protease, Nat. Commun. 12 (2021) 3623. https://doi.org/10.1038/s41467-021-23751-3.

[27] R. Oerlemans, A.J. Ruiz-Moreno, Y. Cong, N. Dinesh Kumar, M.A. Velasco-Velazquez, C.G. Neochoritis, J. Smith, F. Reggiori, M.R. Groves, A. Dömling, Repurposing the HCV NS3-4A protease drug boceprevir as COVID-19 therapeutics, RSC Med. Chem. 12 (2021) 370–379. https://doi.org/10.1039/d0md00367k.

[28] D.W. Kneller, G. Phillips, K.L. Weiss, Q. Zhang, L. Coates, A. Kovalevsky, Direct Observation of Protonation State Modulation in SARS-CoV-2 Main Protease upon Inhibitor Binding with Neutron Crystallography, J. Med. Chem. 64 (2021) 4991–5000. https://doi.org/10.1021/acs.jmedchem.1c00058.

[29] S. Günther, P.Y.A. Reinke, Y. Fernández-García, J. Lieske, T.J. Lane, H.M. Ginn, F.H.M. Koua, C. Ehrt, W. Ewert, D. Oberthuer, X-ray screening identifies active site and allosteric inhibitors of SARS-CoV-2 main protease, Science (80-.). (2021).

[30] D.R. Owen, C.M.N. Allerton, A.S. Anderson, L. Aschenbrenner, M. Avery, S. Berritt, B. Boras, R.D. Cardin, A. Carlo, K.J. Coffman, A. Dantonio, L. Di, H. Eng, R. Ferre, K.S. Gajiwala, S.A. Gibson, S.E. Greasley, B.L. Hurst, E.P. Kadar, A.S. Kalgutkar, J.C. Lee, J. Lee, W. Liu, S.W. Mason, S. Noell, J.J. Novak, R.S. Obach, K. Ogilvie, N.C. Patel, M. Pettersson, D.K. Rai, M.R. Reese, M.F. Sammons, J.G. Sathish, R.S.P. Singh, C.M. Steppan, A.E. Stewart, J.B. Tuttle, L. Updyke, P.R. Verhoest, L. Wei, Q. Yang, Y. Zhu, An oral SARS-CoV-2 M pro inhibitor clinical candidate for the treatment of COVID-19, Science (80-.). 0 (2021) eabl4784. https://doi.org/10.1126/science.abl4784.

[31] R. Robbins, Pfizer Says Its Antiviral Pill is Highly Effective in Treating Covid, New York Times. (2021). https://www.nytimes.com/2021/11/05/health/pfizer-covid-pill.html.

[32] Y.M. Báez-Santos, S.J. Barraza, M.W. Wilson, M.P. Agius, A.M. Mielech, N.M. Davis, S.C. Baker, S.D. Larsen, A.D. Mesecar, X-ray structural and biological evaluation of a series of potent and highly selective inhibitors of human coronavirus papain-like proteases, J. Med. Chem. 57 (2014) 2393–2412. https://doi.org/10.1021/jm401712t.

[33] K. Ratia, S. Pegan, J. Takayama, K. Sleeman, M. Coughlin, S. Baliji, R. Chaudhuri, W. Fu, B.S. Prabhakar, M.E. Johnson, S.C. Baker, A.K. Ghosh, A.D. Mesecar, A noncovalent class of papain-like protease/deubiquitinase inhibitors blocks SARS virus replication, Proc. Natl. Acad. Sci. U. S. A. 105 (2008) 16119–16124. https://doi.org/10.1073/pnas.0805240105.

[34] M.H. Lin, D.C. Moses, C.H. Hsieh, S.C. Cheng, Y.H. Chen, C.Y. Sun, C.Y. Chou, Disulfiram can inhibit MERS and SARS coronavirus papain-like proteases via different modes, Antiviral Res. 150 (2018) 155–163. https://doi.org/10.1016/j.antiviral.2017.12.015.

[35] J. Lei, Y. Kusov, R. Hilgenfeld, Nsp3 of coronaviruses: Structures and functions of a large multi-domain protein, Antiviral Res. 149 (2018) 58–74. https://doi.org/10.1016/j.antiviral.2017.11.001.

[36] L.A. Armstrong, S.M. Lange, V.D. Cesare, S.P. Matthews, R.S. Nirujogi, I. Cole, A. Hope, F. Cunningham, R. Toth, R. Mukherjee, D. Bojkova, F. Gruber, D. Gray, P.G. Wyatt, J. Cinatl, I. Dikic, P. Davies, Y. Kulathu, Biochemical characterization of protease activity of Nsp3 from SARS-CoV-2 and its inhibition by nanobodies, PLoS One. 16 (2021) e0253364. https://doi.org/10.1371/journal.pone.0253364.

[37] J. Osipiuk, S.A. Azizi, S. Dvorkin, M. Endres, R. Jedrzejczak, K.A. Jones, S. Kang, R.S. Kathayat, Y. Kim, V.G. Lisnyak, S.L. Maki, V. Nicolaescu, C.A. Taylor, C. Tesar, Y.A. Zhang, Z. Zhou, G. Randall, K. Michalska, S.A. Snyder, B.C. Dickinson, A. Joachimiak, Structure of papain-like protease from SARS-CoV-2 and its complexes with non-covalent inhibitors, Nat. Commun. 12 (2021) 1–9. https://doi.org/10.1038/s41467-021-21060-3.

[38] Z. Fu, B. Huang, J. Tang, S. Liu, M. Liu, Y. Ye, Z. Liu, Y. Xiong, W. Zhu, D. Cao, J. Li, X. Niu, H. Zhou, Y.J. Zhao, G. Zhang, H. Huang, The complex structure of GRL0617 and SARS-CoV-2 PLpro reveals a hot spot for antiviral drug discovery, Nat. Commun. 12 (2021) 1–12. https://doi.org/10.1038/s41467-020-20718-8.

[39] H. Shan, J. Liu, J. Shen, J. Dai, G. Xu, K. Lu, C. Han, Y. Wang, X. Xu, Y. Tong, H. Xiang, Z. Ai, G. Zhuang, J. Hu, Z. Zhang, Y. Li, L. Pan, L. Tan, Development of potent and selective inhibitors targeting the papain-like protease of SARS-CoV-2, Cell Chem. Biol. 28 (2021) 855–865.e9. https://doi.org/10.1016/j.chembiol.2021.04.020.

[40] Z. Shen, K. Ratia, L. Cooper, D. Kong, H. Lee, Y. Kwon, Y. Li, S. Alqarni, F. Huang, O. Dubrovskyi, L. Rong, G.R.J. Thatcher, R. Xiong, Design of SARS-CoV-2 PLpro Inhibitors for COVID-19 Antiviral Therapy Leveraging Binding Cooperativity, J. Med. Chem. (2021). https://doi.org/10.1021/acs.jmedchem.1c01307.

[41] X. Gao, B. Qin, P. Chen, K. Zhu, P. Hou, J.A. Wojdyla, M. Wang, S. Cui, Crystal structure of SARS-CoV-2 papain-like protease, Acta Pharm. Sin. B. 11 (2021) 237–245.

[42] T. Klemm, G. Ebert, D. Calleja, C. Allison, L. Richardson, J. Bernardini, B. Lu, N. Kuchel, C. Grohmann, Y. Shibata, Z.Y. Gan, J. Cooney, M. Doerflinger, A. Au, T. Blackmore, P. Geurink, H. Ovaa, J. Newman, A. Riboldi-Tunnicliffe, P. Czabotar, J. Mitchell, R. Feltham, B. Lechtenberg, K. Lowes, G. Dewson, M. Pellegrini, G. Lessene, D. Komander, Mechanism and inhibition of SARS-CoV-2 PLpro, EMBO J. 39 (2020) e106275. https://doi.org/10.1101/2020.06.18.160614.

[43] S. Pushpakom, F. Iorio, P.A. Eyers, K.J. Escott, S. Hopper, A. Wells, A. Doig, T. Guilliams, J. Latimer, C. McNamee, Drug repurposing: progress, challenges and recommendations, Nat. Rev. Drug Discov. 18 (2019) 41–58.

[44] G. Li, E. De Clercq, Therapeutic options for the 2019 novel coronavirus (2019-nCoV), Nat. Rev. Drug Discov. 19 (2020) 149–150. https://doi.org/10.1038/d41573-020-00016-0.

[45] E.H.G. da Cruz, C.M.B. Hussene, G.G. Dias, E.B.T. Diogo, I.M.M. de Melo, B.L. Rodrigues, M.G. da Silva, W.O. Valença, C.A. Camara, R.N. de Oliveira, Y.G. de Paiva, M.O.F. Goulart, B.C. Cavalcanti, C. Pessoa, E.N. da Silva Júnior, 1,2,3-Triazole-, arylamino- and thio-substituted 1,4-naphthoquinones: Potent antitumor activity, electrochemical aspects, and bioisosteric replacement of C-ring-modified lapachones, Bioorganic Med. Chem. 22 (2014) 1608–1619. https://doi.org/10.1016/j.bmc.2014.01.033.

[46] R.L. de Carvalho, G.A.M. Jardim, A.C.C. Santos, M.H. Araujo, W.X.C. Oliveira, A.C.S. Bombaça, R.F.S. Menna-Barreto, E. Gopi, E. Gravel, E. Doris, E.N. da Silva Júnior, Combination of Aryl Diselenides/Hydrogen Peroxide and Carbon-Nanotube/Rhodium Nanohybrids for Naphthol Oxidation: An Efficient Route towards Trypanocidal Quinones, Chemistry. 24 (2018) 15227–15235. https://doi.org/10.1002/chem.201802773.

[47] G.A.M. Jardim, T.L. Silva, M.O.F. Goulart, C.A. de Simone, J.M.C. Barbosa, K. Salomão, S.L. de Castro, J.F. Bower, E.N. da Silva Júnior, Rhodium-catalyzed C-H bond activation for the synthesis of quinonoid compounds: Significant Anti-Trypanosoma cruzi activities and electrochemical studies of functionalized quinones, Eur. J. Med. Chem. 136 (2017) 406–419. https://doi.org/10.1016/j.ejmech.2017.05.011.

[48] D.M. Mellott, C. Te Tseng, A. Drelich, P. Fajtová, B.C. Chenna, D.H. Kostomiris, J. Hsu, J. Zhu, Z.W. Taylor, K.I. Kocurek, V. Tat, A. Katzfuss, L. Li, M.A. Giardini, D. Skinner, K. Hirata, M.C. Yoon, S. Beck, A.F. Carlin, A.E. Clark, L. Beretta, D. Maneval, V. Hook, F. Frueh, B.L. Hurst, H. Wang, F.M. Raushel, A.J. O’Donoghue, J.L. De Siqueira-Neto, T.D. Meek, J.H. McKerrow, A Clinical-Stage Cysteine Protease Inhibitor blocks SARS-CoV-2 Infection of Human and Monkey Cells, ACS Chem. Biol. 16 (2021) 642–650. https://doi.org/10.1021/acschembio.0c00875.

[49] R. Lu, X. Zhao, J. Li, P. Niu, B. Yang, H. Wu, W. Wang, H. Song, B. Huang, N. Zhu, Y. Bi, X. Ma, F. Zhan, L. Wang, T. Hu, H. Zhou, Z. Hu, W. Zhou, L. Zhao, J. Chen, Y. Meng, J. Wang, Y. Lin, J. Yuan, Z. Xie, J. Ma, W.J. Liu, D. Wang, W. Xu, E.C. Holmes, G.F. Gao, G. Wu, W. Chen, W. Shi, W. Tan, Genomic characterisation and epidemiology of 2019 novel coronavirus: implications for virus origins and receptor binding, Lancet. 395 (2020) 565–574. https://doi.org/10.1016/S0140-6736(20)30251-8.

[50] Gilead Sciences Inc, Gilead’s Investigational Antiviral Remdesivir Receives U.S. Food and Drug Administration Emergency Use Authorization for the Treatment of COVID-19, Https://Www.Gilead.Com/. (2020). https://www.gilead.com/news-and-press/press-room/press-releases/2020/5/gileads-investigational-antiviral-remdesivir-receives-us-food-and-drug-administration-emergency-use-authorization-for-the-treatment-of-covid19 (accessed April 21, 2021).

[51] Federation Drug American (FDA), Fact Sheet for Health Care Providers Emergency Use Authorization of Bamlanivimab and Etesevimab, Http://Www.Fda.Gov/. (2020) 1–36. https://www.cdc.gov/growthcharts/clinical_charts.htm %0A https://www.cdc.gov/growthcharts/clinical_charts.htm,%0A https://www.fda.gov/media/137566/download (accessed April 21, 2021).

[52] J. Pardo, A.M. Shukla, G. Chamarthi, A. Gupte, The journey of remdesivir: From Ebola to COVID-19, Drugs Context. 9 (2020). https://doi.org/10.7573/DIC.2020-4-14.

[53] J.H. Beigel, K.M. Tomashek, L.E. Dodd, A.K. Mehta, B.S. Zingman, A.C. Kalil, E. Hohmann, H.Y. Chu, A. Luetkemeyer, S. Kline, D. Lopez de Castilla, R.W. Finberg, K. Dierberg, V. Tapson, L. Hsieh, T.F. Patterson, R. Paredes, D.A. Sweeney, W.R. Short, G. Touloumi, D.C. Lye, N. Ohmagari, M. Oh, G.M. Ruiz-Palacios, T. Benfield, G. Fätkenheuer, M.G. Kortepeter, R.L. Atmar, C.B. Creech, J. Lundgren, A.G. Babiker, S. Pett, J.D. Neaton, T.H. Burgess, T. Bonnett, M. Green, M. Makowski, A. Osinusi, S. Nayak, H.C. Lane, Remdesivir for the Treatment of Covid-19 — Final Report, N. Engl. J. Med. 383 (2020) 1813–1826. https://doi.org/10.1056/nejmoa2007764.

[54] E.K. McCreary, J.M. Pogue, Coronavirus disease 2019 treatment: A review of early and emerging options, in: Open Forum Infect. Dis., Oxford University Press US, 2020: p. ofaa105. https://doi.org/10.1093/ofid/ofaa105.

[55] M.S. Hossan, A. Fatima, M. Rahmatullah, T.J. Khoo, V. Nissapatorn, A. V. Galochkina, A. V. Slita, A.A. Shtro, Y. Nikolaeva, V. V. Zarubaev, C. Wiart, Antiviral activity of Embelia ribes Burm. f. against influenza virus in vitro, Arch. Virol. 163 (2018) 2121–2131. https://doi.org/10.1007/s00705-018-3842-6.

[56] M.K. Parvez, M. Tabish Rehman, P. Alam, M.S. Al-Dosari, S.I. Alqasoumi, M.F. Alajmi, Plant-derived antiviral drugs as novel hepatitis B virus inhibitors: Cell culture and molecular docking study, Saudi Pharm. J. 27 (2019) 389–400. https://doi.org/10.1016/j.jsps.2018.12.008.

[57] F. Caruso, M. Rossi, J.Z. Pedersen, S. Incerpi, Computational studies reveal mechanism by which quinone derivatives can inhibit SARS-CoV-2. Study of embelin and two therapeutic compounds of interest, methyl prednisolone and dexamethasone, J. Infect. Public Health. 13 (2020) 1868–1877. https://doi.org/10.1016/j.jiph.2020.09.015.

[58] Y.B. Ryu, S.J. Park, Y.M. Kim, J.Y. Lee, W.D. Seo, J.S. Chang, K.H. Park, M.C. Rho, W.S. Lee, SARS-CoV 3CLpro inhibitory effects of quinone-methide triterpenes from Tripterygium regelii, Bioorganic Med. Chem. Lett. 20 (2010) 1873–1876. https://doi.org/10.1016/j.bmcl.2010.01.152.

[59] E.N. da Silva Júnior, M.C.B.V. de Souza, A. V. Pinto, M. do C.F.R. Pinto, M.O.F. Goulart, F.W.A. Barros, C. Pessoa, L. V. Costa-Lotufo, R.C. Montenegro, M.O. de Moraes, V.F. Ferreira, Synthesis and potent antitumor activity of new arylamino derivatives of nor-β-lapachone and nor-α-lapachone, Bioorganic Med. Chem. 15 (2007) 7035–7041. https://doi.org/10.1016/j.bmc.2007.07.043.

[60] E.N. da Silva Júnior, M.C.B.V. de Souza, M.C. Fernandes, R.F.S. Menna-Barreto, M. do C.F.R. Pinto, F. de Assis Lopes, C.A. de Simone, C.K.Z. Andrade, A. V. Pinto, V.F. Ferreira, S.L. de Castro, Synthesis and anti-Trypanosoma cruzi activity of derivatives from nor-lapachones and lapachones, Bioorganic Med. Chem. 16 (2008) 5030–5038. https://doi.org/10.1016/j.bmc.2008.03.032.

[61] E.N. da Silva Júnior, T.T. Guimarães, R.F.S. Menna-Barreto, M. do C.F.R. Pinto, C.A. de Simone, C. Pessoa, B.C. Cavalcanti, J.R. Sabino, C.K.Z. Andrade, M.O.F. Goulart, S.L. de Castro, A. V. Pinto, The evaluation of quinonoid compounds against Trypanosoma cruzi: Synthesis of imidazolic anthraquinones, nor-β-lapachone derivatives and β-lapachone-based 1,2,3-triazoles, Bioorganic Med. Chem. 18 (2010) 3224–3230. https://doi.org/10.1016/j.bmc.2010.03.029.

[62] E.N. da Silva Júnior, C.F. de Deus, B.C. Cavalcanti, C. Pessoa, L. V. Costa-Lotufo, R.C. Montenegro, M.O. de Moraes, M.D.C.F.R. Pinto, C.A. de Simone, V.F. Ferreira, M.O.F. Goulart, C.K.Z. Andrade, A. V. Pinto, 3-Arylamino and 3-alkoxy-nor-β-lapachone Derivatives: Synthesis and Cytotoxicity against Cancer Cell Lines, J. Med. Chem. 53 (2010) 504–508. https://doi.org/10.1021/jm900865m.

[63] A.A. de Souza, M.A.B.F. de Moura, F.C. de Abreu, M.O.F. Goulart, E.N. da Silva Jr., A. V. Pinto, V.F. Ferreira, R. Moscoso, L.J. Núñez-Vergara, J.A. Squella, Electrochemical study, on mercury, of a Meta-nitroarylamine derivative of nor-β-lapachone, an antitumor and trypanocidal compound, Quim. Nova. 33 (2010) 2075–2079. https://doi.org/10.1590/s0100-40422010001000013.

[64] E.H.G. da Cruz, M.A. Silvers, G.A.M. Jardim, J.M. Resende, B.C. Cavalcanti, I.S. Bomfim, C. Pessoa, C.A. de Simone, G. V. Botteselle, A.L. Braga, D.K. Nair, I.N.N. Namboothiri, D.A. Boothman, E.N. da Silva Júnior, Synthesis and antitumor activity of selenium-containing quinone-based triazoles possessing two redox centres, and their mechanistic insights, Eur. J. Med. Chem. 122 (2016) 1– 16. https://doi.org/10.1016/j.ejmech.2016.06.019.

[65] A.A. Vieira, I.R. Brandão, W.O. Valença, C.A. de Simone, B.C. Cavalcanti, C. Pessoa, T.R. Carneiro, A.L. Braga, E.N. da Silva, Hybrid compounds with two redox centres: Modular synthesis of chalcogen-containing lapachones and studies on their antitumor activity, Eur. J. Med. Chem. 101 (2015) 254–265. https://doi.org/10.1016/j.ejmech.2015.06.044.

[66] A. Kharma, C. Jacob, Í.A.O. Bozzi, G.A.M. Jardim, A.L. Braga, K. Salomão, C.C. Gatto, M.F.S. Silva, C. Pessoa, M. Stangier, L. Ackermann, E.N. da Silva Júnior, Electrochemical Selenation/Cyclization of Quinones: A Rapid, Green and Efficient Access to Functionalized Trypanocidal and Antitumor Compounds, European J. Org. Chem. 2020 (2020) 4474–4486. https://doi.org/10.1002/ejoc.202000216.

[67] R.G. Almeida, W.O. Valença, L.G. Rosa, C.A. de Simone, S.L. de Castro, J.M.C. Barbosa, D.P. Pinheiro, C.R.K. Paier, G.G.C. de Carvalho, C. Pessoa, M.O.F. Goulart, A. Kharma, E.N. da Silva Júnior, Synthesis of quinone imine and sulphur-containing compounds with antitumor and trypanocidal activities: Redox and biological implications, RSC Med. Chem. 11 (2020) 1145–1160. https://doi.org/10.1039/d0md00072h.

[68] G.A.M. Jardim, T.T. Guimarães, M.D.C.F.R. Pinto, B.C. Cavalcanti, K.M. de Farias, C. Pessoa, C.C. Gatto, D.K. Nair, I.N.N. Namboothiri, E.N. da Silva Júnior, Naphthoquinone-based chalcone hybrids and derivatives: Synthesis and potent activity against cancer cell lines, Med. Chem. Commun. 6 (2015) 120– 150. https://doi.org/10.1039/c4md00371c.

[69] B.C. Cavalcanti, I.O. Cabral, F.A.R. Rodrigues, F.W.A. Barros, D.D. Rocha, H.I.F. Magalhães, D.J. Moura, J. Saffi, J.A.P. Henriques, T.S.C. Carvalho, M.O. Moraes, C. Pessoa, I.M.M. de Melo, E.N. da Silva Júnior, Potent antileukemic action of naphthoquinoidal compounds: Evidence for an intrinsic death mechanism based on oxidative stress and inhibition of DNA repair, J. Braz. Chem. Soc. 24 (2013) 145–163. https://doi.org/10.1590/S0103-50532013000100019.

[70] S.L. de Castro, F.S. Emery, E.N. da Silva Júnior, Synthesis of quinoidal molecules: Strategies towards bioactive compounds with an emphasis on lapachones, Eur. J. Med. Chem. 69 (2013) 678–700. https://doi.org/10.1016/j.ejmech.2013.07.057.

[71] G.A.M. Jardim, W.J. Reis, M.F. Ribeiro, F.M. Ottoni, R.J. Alves, T.L. Silva, M.O.F. Goulart, A.L. Braga, R.F.S. Menna-Barreto, K. Salomão, S.L. de Castro, E.N. da Silva Júnior, On the investigation of hybrid quinones: Synthesis, electrochemical studies and evaluation of trypanocidal activity, RSC Adv. 5 (2015) 78047–78060. https://doi.org/10.1039/c5ra16213k.

[72] G.G. Dias, T. Rogge, R. Kuniyil, C. Jacob, R.F.S. Menna-Barreto, E.N. da Silva Júnior, L. Ackermann, Ruthenium-catalyzed C-H oxygenation of quinones by weak O-coordination for potent trypanocidal agents, Chem. Commun. 54 (2018) 12840–12843. https://doi.org/10.1039/c8cc07572g.

[73] T. V. Baiju, R.G. Almeida, S.T. Sivanandan, C.A. de Simone, L.M. Brito, B.C. Cavalcanti, C. Pessoa, I.N.N. Namboothiri, E.N. da Silva Júnior, Quinonoid compounds via reactions of lawsone and 2-aminonaphthoquinone with α-bromonitroalkenes and nitroallylic acetates: Structural diversity by C-ring modification and cytotoxic evaluation against cancer cells, Eur. J. Med. Chem. 151 (2018) 686–704. https://doi.org/10.1016/j.ejmech.2018.03.079.

[74] H. Suginome, A. Konishi, H. Sakurai, H. Minakawa, T. Takeda, H. Senboku, M. Tokuda, K. Kobayashi, Photoinduced molecular transformations. Part 156. New photoadditions of 2-hydroxy-1,4-naphthoquinones with naphthols and their derivatiyes, Tetrahedron. 51 (1995) 1377–1386. https://doi.org/10.1016/0040-4020(94)01026-V.

[75] J.M. Wood, N.S. Satam, R.G. Almeida, V.S. Cristani, D.P. de Lima, L. Dantas-Pereira, K. Salomão, R.F.S. Menna-Barreto, I.N.N. Namboothiri, J.F. Bower, E.N. da Silva Júnior, Strategies towards potent trypanocidal drugs: Application of Rh-catalyzed [2 + 2 + 2] cycloadditions, sulfonyl phthalide annulation and nitroalkene reactions for the synthesis of substituted quinones and their evaluation against Trypanosoma cruzi, Bioorganic Med. Chem. 28 (2020) 115565. https://doi.org/10.1016/j.bmc.2020.115565.

[76] E.N. da Silva Júnior, R.L. de Carvalho, R.G. Almeida, L.G. Rosa, F. Fantuzzi, T. Rogge, P.M.S. Costa, C. Pessoa, C. Jacob, L. Ackermann, Ruthenium(II)-Catalyzed Double Annulation of Quinones: Step-Economical Access to Valuable Bioactive Compounds, Chem. - A Eur. J. 26 (2020) 10981–10986. https://doi.org/10.1002/chem.202001434.

[77] V.K. Tandon, D.B. Yadav, R. V. Singh, M. Vaish, A.K. Chaturvedi, P.K. Shukla, Synthesis and biological evaluation of novel 1,4-naphthoquinone derivatives as antibacterial and antiviral agents, Bioorganic Med. Chem. Lett. 15 (2005) 3463– 3466. https://doi.org/10.1016/j.bmcl.2005.04.075.

[78] G. Krishnamoorthy, S.P. Webb, T. Nguyen, P.K. Chowdhury, M. Halder, N.J. Wills, S. Carpenter, G.A. Kraus, M.S. Gordon, J.W. Petrich, Synthesis of hydroxy and methoxy perylene quinones, their spectroscopic and computational characterization, and their antiviral activity, Photochem. Photobiol. 81 (2005) 924. https://doi.org/10.1562/2004-11-23-ra-378r1.1.

[79] L.R. Silva, A.S. Guimarães, J. do Nascimento, I.J. do Santos Nascimento, E.B. da Silva, J.H. McKerrow, S.H. Cardoso, E.F. da Silva-Júnior, Computer-aided design of 1, 4-naphthoquinone-based inhibitors targeting cruzain and rhodesain cysteine proteases, Bioorg. Med. Chem. 41 (2021) 116213.

[80] E.N. da Silva Jr., R.F.S. Menna-Barreto, M. do C.F.R. Pinto, R.S.F. Silva, D. V. Teixeira, M.C.B.V. de Souza, C.A. De Simone, S.L. De Castro, V.F. Ferreira, A. V. Pinto, Naphthoquinoidal [1,2,3]-triazole, a new structural moiety active against Trypanosoma cruzi, Eur. J. Med. Chem. 43 (2008) 1774–1780. https://doi.org/10.1016/j.ejmech.2007.10.015.

[81] E.N. da Silva Júnior, M.A.B.F. de Moura, A. V. Pinto, M. do C.F.R. Pinto, M.C.B.V. de Souza, A.J. Araújo, C. Pessoa, L. V. Costa-Lotufo, R.C. Montenegro, M.O. de Moraes, V.F. Ferreira, M.O.F. Goulart, Cytotoxic, trypanocidal activities and physicochemical parameters of nor-β-lapachone-based 1,2,3-triazoles, J. Braz. Chem. Soc. 20 (2009) 635–643. https://doi.org/10.1590/s0103-50532009000400007.

[82] E.N. da Silva Júnior, I.M.M. de Melo, E.B.T. Diogo, V.A. Costa, J.D. de Souza Filho, W.O. Valença, C.A. Camara, R.N. de Oliveira, A.S. de Araujo, F.S. Emery, M.R. dos Santos, C.A. de Simone, R.F.S. Menna-Barreto, S.L. de Castro, On the search for potential anti-Trypanosoma cruzi drugs: Synthesis and biological evaluation of 2-hydroxy-3-methylamino and 1,2,3-triazolic naphthoquinoidal compounds obtained by click chemistry reactions, Eur. J. Med. Chem. 52 (2012) 304–312. https://doi.org/10.1016/j.ejmech.2012.03.039.

[83] M.F.C. Cardoso, P.C. Rodrigues, M.E.I.M. Oliveira, I.L. Gama, I.M.C.B. da Silva, I.O. Santos, D.R. Rocha, R.T. Pinho, V.F. Ferreira, M.C.B.V. de Souza, F.D.C. da Silva, F.P. Silva-Jr, Synthesis and evaluation of the cytotoxic activity of 1,2-furanonaphthoquinones tethered to 1,2,3-1H-triazoles in myeloid and lymphoid leukemia cell lines, Eur. J. Med. Chem. 84 (2014) 708–717. https://doi.org/10.1016/j.ejmech.2014.07.079.

[84] G.A.M. Jardim, E.H.G. Cruz, W.O. Valença, J.M. Resende, B.L. Rodrigues, D.F. Ramos, R.N. Oliveira, P.E.A. Silva, E.N. da Silva Júnior, On the search for potential antimycobacterial drugs: Synthesis of naphthoquinoidal, phenazinic and 1,2,3-triazolic compounds and evaluation against Mycobacterium tuberculosis, J. Braz. Chem. Soc. 26 (2015) 1013–1027. https://doi.org/10.5935/0103-5053.20150067.

[85] F.S. dos Santos, G.G. Dias, R.P. de Freitas, L.S. Santos, G.F. de Lima, H.A. Duarte, C.A. de Simone, L.M.S.L. Rezende, M.J.X. Vianna, J.R. Correa, B.A.D. Neto, E.N. da Silva Júnior, Redox Center Modification of Lapachones towards the Synthesis of Nitrogen Heterocycles as Selective Fluorescent Mitochondrial Imaging Probes, European J. Org. Chem. 2017 (2017) 3763–3773. https://doi.org/10.1002/ejoc.201700227.

[86] S.B.B.B. Bahia, W.J. Reis, G.A.M. Jardim, F.T. Souto, C.A. de Simone, C.C. Gatto, R.F.S. Menna-Barreto, S.L. de Castro, B.C. Cavalcanti, C. Pessoa, M.H. Araujo, E.N. da Silva Júnior, Molecular hybridization as a powerful tool towards multitarget quinoidal systems: Synthesis, trypanocidal and antitumor activities of naphthoquinone-based 5-iodo-1,4-disubstituted-, 1,4- and 1,5-disubstituted-1,2,3-triazoles, Med. Chem. Commun. 7 (2016) 1555–1563. https://doi.org/10.1039/c6md00216a.

[87] G.A.M. Jardim, D.J.B. Lima, W.O. Valença, D.J.B. Lima, B.C. Cavalcanti, C. Pessoa, J. Rafique, A.L. Braga, C. Jacob, E.N. Da Silva Júnior, E.H.G. Da Cruz, Synthesis of Selenium-Quinone Hybrid Compounds with Potential Antitumor Activity via Rh-Catalyzed C-H Bond Activation and Click Reactions, Molecules. 23 (2018) 83. https://doi.org/10.3390/molecules23010083.

[88] T.B. Gontijo, R.P. de Freitas, G.F. de Lima, L.C.D. de Rezende, L.F. Pedrosa, T.L. Silva, M.O.F. Goulart, B.C. Cavalcanti, C. Pessoa, M.P. Bruno, J.R. Corrêa, F.S. Emery, E.N. da Silva Júnior, Novel fluorescent lapachone-based BODIPY: Synthesis, computational and electrochemical aspects, and subcellular localisation of a potent antitumour hybrid quinone, Chem. Commun. 52 (2016) 13281–13284. https://doi.org/10.1039/c6cc07054j.

[89] T.B. Gontijo, R.P. de Freitas, F.S. Emery, L.F. Pedrosa, J.B. Vieira Neto, B.C. Cavalcanti, C. Pessoa, A. King, F. de Moliner, M. Vendrell, E.N. da Silva Júnior, On the synthesis of quinone-based BODIPY hybrids: New insights on antitumor activity and mechanism of action in cancer cells, Bioorganic Med. Chem. Lett. 27 (2017) 4446–4456. https://doi.org/10.1016/j.bmcl.2017.08.007.

[90] W.O. Valença, T. V. Baiju, F.G. Brito, M.H. Araujo, C. Pessoa, B.C. Cavalcanti, C.A. de Simone, C. Jacob, I.N.N. Namboothiri, E.N. da Silva Júnior, Synthesis of Quinone-Based N-Sulfonyl-1,2,3-triazoles: Chemical Reactivity of Rh(II) Azavinyl Carbenes and Antitumor Activity, ChemistrySelect. 2 (2017) 4301– 4308. https://doi.org/10.1002/slct.201700885.

[91] E.B.T. Diogo, G.G. Dias, B.L. Rodrigues, T.T. Guimarães, W.O. Valença, C.A. Camara, R.N. de Oliveira, M.G. da Silva, V.F. Ferreira, Y.G. de Paiva, M.O.F. Goulart, R.F.S. Menna-Barreto, S.L. de Castro, E.N. da Silva Júnior, Synthesis and anti-Trypanosoma cruzi activity of naphthoquinone-containing triazoles: Electrochemical studies on the effects of the quinoidal moiety, Bioorganic Med. Chem. 21 (2013) 6337–6348. https://doi.org/10.1016/j.bmc.2013.08.055.

[92] V.N. Melo, W.M. Dantas, C.A. Camara, R.N. De Oliveira, Synthesis of 2,3-unsaturated alkynyl O-glucosides from tri-O-acetyl-d-glucal by using montmorillonite K-10/iron(III) chloride hexahydrate with subsequent copper(I)-catalyzed 1,3-dipolar cycloaddition, Synth. 47 (2015) 3529–3541. https://doi.org/10.1055/s-0034-1378829.

[93] R.N. De Oliveira, A.L. De Xavier, B.M. Guimaraes, V.N.E. Melo, W.O. Valença, W.S. Nascimento Do, P.L.F. Da Costa, C.A. Camara, Combining clays and ultrasound irradiation for an o-acetylation reaction of N-glucopyranosyl and other molecules, J. Chil. Chem. Soc. 59 (2014) 2610–2614. https://doi.org/10.4067/S0717-97072014000300018.

[94] Z. Cheng, W.O. Valença, G.G. Dias, J. Scott, N.D. Barth, F. de Moliner, G.B.P. Souza, R.J. Mellanby, M. Vendrell, E.N. da Silva Júnior, Natural product-inspired profluorophores for imaging NQO1 activity in tumour tissues, Bioorganic Med. Chem. 27 (2019) 3938–3946. https://doi.org/10.1016/j.bmc.2019.07.017.

[95] V. V. Rostovtsev, L.G. Green, V. V. Fokin, K.B. Sharpless, A stepwise huisgen cycloaddition process: Copper(I)-catalyzed regioselective “ligation” of azides and terminal alkynes, Angew. Chemie - Int. Ed. 41 (2002) 2596–2599. https://doi.org/10.1002/1521-3773(20020715)41:14<2596::AID-ANIE2596>3.0.CO;2-4.

[96] F. de Moliner, A. King, G.G. Dias, G.F. de Lima, C.A. de Simone, E.N. da Silva Júnior, M. Vendrell, Quinone-derived π-extended phenazines as new fluorogenic probes for live-cell imaging of lipid droplets, Front. Chem. 6 (2018) 339. https://doi.org/10.3389/fchem.2018.00339.

[97] R.S.F. Silva, M.B. De Amorim, M.D.C.F.R. Pinto, F.S. Emery, M.O.F. Goulart, A. V. Pinto, Chemoselective oxidation of benzophenazines by m-CPBA: N-oxidation vs. oxidative cleavage, J. Braz. Chem. Soc. 18 (2007) 759–764. https://doi.org/10.1590/S0103-50532007000400014.

[98] G.A.M. Jardim, W.X.C. Oliveira, R.P. de Freitas, R.F.S. Menna-Barreto, T.L. Silva, M.O.F. Goulart, E.N. da Silva Júnior, Direct sequential C-H iodination/organoyl-thiolation for the benzenoid A-ring modification of quinonoid deactivated systems: A new protocol for potent trypanocidal quinones, Org. Biomol. Chem. 16 (2018) 1686–1691. https://doi.org/10.1039/c8ob00196k.

[99] R.G. Almeida, R.L. De Carvalho, M.P. Nunes, R.S. Gomes, L.F. Pedrosa, C.A. De Simone, E. Gopi, V. Geertsen, E. Gravel, E. Doris, E.N. da Silva Júnior, Carbon nanotube-ruthenium hybrid towards mild oxidation of sulfides to sulfones: Efficient synthesis of diverse sulfonyl compounds, Catal. Sci. Technol. 9 (2019) 2742–2748. https://doi.org/10.1039/c9cy00384c.

[100] G.A.M. Jardim, Í.A.O. Bozzi, W.X.C. Oliveira, C. Mesquita-Rodrigues, R.F.S. Menna-Barreto, R.A. Kumar, E. Gravel, E. Doris, A.L. Braga, E.N. da Silva Júnior, Copper complexes and carbon nanotube-copper ferrite-catalyzed benzenoid A-ring selenation of quinones: An efficient method for the synthesis of trypanocidal agents, New J. Chem. 43 (2019) 13751–13763. https://doi.org/10.1039/c9nj02026h.

[101] G.A.M. Jardim, J.F. Bower, E.N. da Silva Júnior, Rh-Catalyzed Reactions of 1,4-Benzoquinones with Electrophiles: C-H Iodination, Bromination, and Phenylselenation, Org. Lett. 18 (2016) 4454–4457. https://doi.org/10.1021/acs.orglett.6b01586.

[102] G.G. Dias, T.A. d. Nascimento, A.K.A. de Almeida, A.C.S. Bombaça, R.F.S. Menna-Barreto, C. Jacob, S. Warratz, E.N. da Silva Júnior, L. Ackermann, Ruthenium(II)-Catalyzed C–H Alkenylation of Quinones: Diversity-Oriented Strategy for Trypanocidal Compounds, European J. Org. Chem. 2019 (2019) 2344–2353. https://doi.org/10.1002/ejoc.201900004.

[103] S.N. Sunassee, C.G.L. Veale, N. Shunmoogam-Gounden, O. Osoniyi, D.T. Hendricks, M.R. Caira, J.A. de La Mare, A.L. Edkins, A. V. Pinto, E.N. da Silva Júnior, M.T. Davies-Coleman, Cytotoxicity of lapachol, β-lapachone and related synthetic 1,4-naphthoquinones against oesophageal cancer cells, Eur. J. Med. Chem. 62 (2013) 98–110. https://doi.org/10.1016/j.ejmech.2012.12.048.

[104] T. Kumar, N. Satam, I.N.N. Namboothiri, Hauser–Kraus Annulation of Phthalides with Nitroalkenes for the Synthesis of Fused and Spiro Heterocycles, European J. Org. Chem. 2016 (2016) 3316–3321. https://doi.org/10.1002/ejoc.201600390.

[105] A. Suresh, T. V. Baiju, T. Kumar, I.N.N. Namboothiri, Synthesis of Spiro- and Fused Heterocycles via (4+4) Annulation of Sulfonylphthalide with o-Hydroxystyrenyl Derivatives, J. Org. Chem. 84 (2019) 3158–3168. https://doi.org/10.1021/acs.joc.8b03039.

[106] J.M. Wood, E.N. da Silva Júnior, J.F. Bower, Rh-Catalyzed [2 + 2 + 2] Cycloadditions with Benzoquinones: De Novo Access to Naphthoquinones for Lignan and Type II Polyketide Synthesis, Org. Lett. 22 (2020) 265–269. https://doi.org/10.1021/acs.orglett.9b04266.

[107] G.A.M. Jardim, E.N. da Silva Júnior, J.F. Bower, Overcoming naphthoquinone deactivation: Rhodium-catalyzed C-5 selective C-H iodination as a gateway to functionalized derivatives, Chem. Sci. 7 (2016) 3780–3784. https://doi.org/10.1039/c6sc00302h.

[108] W.J. Reis, Í.A.O. Bozzi, M.F. Ribeiro, P.C.B. Halicki, L.A. Ferreira, P.E. Almeida da Silva, D.F. Ramos, C.A. de Simone, E.N. da Silva Júnior, Design of hybrid molecules as antimycobacterial compounds: Synthesis of isoniazid-naphthoquinone derivatives and their activity against susceptible and resistant strains of Mycobacterium tuberculosis, Bioorganic Med. Chem. 27 (2019) 4143– 4150. https://doi.org/10.1016/j.bmc.2019.07.045.

[109] K.C.G. Moura, P.F. Carneiro, M.D.C.F.R. Pinto, J.A. da Silva, V.R.S. Malta, C.A. de Simone, G.G. Dias, G.A.M. Jardim, J. Cantos, T.S. Coelho, P.E.A. da Silva, E.N. da Silva Jr., 1,3-Azoles from ortho-naphthoquinones: Synthesis of aryl substituted imidazoles and oxazoles and their potent activity against Mycobacterium tuberculosis, Bioorganic Med. Chem. 20 (2012) 6482–6488. https://doi.org/10.1016/j.bmc.2012.08.041.

[110] G.G. Dias, P.V.B. Pinho, H.A. Duarte, J.M. Resende, A.B.B. Rosa, J.R. Correa, B.A.D. Neto, E.N. da Silva Júnior, Fluorescent oxazoles from quinones for bioimaging applications, RSC Adv. 6 (2016) 76053–76063. https://doi.org/10.1039/c6ra14701a.

[111] G.G. Dias, B.L. Rodrigues, J.M. Resende, H.D.R. Calado, C.A. de Simone, V.H.C. Silva, B.A.D. Neto, M.O.F. Goulart, F.R. Ferreira, A.S. Meira, C. Pessoa, J.R. Correa, E.N. da Silva Júnior, Selective endocytic trafficking in live cells with fluorescent naphthoxazoles and their boron complexes, Chem. Commun. 51 (2015) 9141–9144. https://doi.org/10.1039/c5cc02383a.

[112] V. Grum-Tokars, K. Ratia, A. Begaye, S.C. Baker, A.D. Mesecar, Evaluating the 3C-like protease activity of SARS-Coronavirus: Recommendations for standardized assays for drug discovery, Virus Res. 133 (2008) 63–73. https://doi.org/10.1016/j.virusres.2007.02.015.

[113] H. xia Su, S. Yao, W. feng Zhao, M. jun Li, J. Liu, W. juan Shang, H. Xie, C. qiang Ke, H. chen Hu, M. na Gao, K. qian Yu, H. Liu, J. shan Shen, W. Tang, L. ke Zhang, G. fu Xiao, L. Ni, D. wen Wang, J. ping Zuo, H. liang Jiang, F. Bai, Y. Wu, Y. Ye, Y. chun Xu, Anti-SARS-CoV-2 activities in vitro of Shuanghuanglian preparations and bioactive ingredients, Acta Pharmacol. Sin. 41 (2020) 1167–1177. https://doi.org/10.1038/s41401-020-0483-6.

[114] A.D. Mesecar, A taxonomically-driven approach to development of potent, broad-spectrum inhibitors of coronavirus main protease including SARS-CoV-2 (COVID-19), Be Publ. (2020).

[115] M. Bzówka, K. Mitusińska, A. Raczyńska, A. Samol, J.A. Tuszyński, A. Góra, Structural and Evolutionary Analysis Indicate That the SARS-CoV-2 Mpro Is a Challenging Target for Small-Molecule Inhibitor Design, Int. J. Mol. Sci. 21 (2020) 3099. https://doi.org/10.3390/ijms21093099.

[116] D.W. Kneller, G. Phillips, H.M. O’Neill, R. Jedrzejczak, L. Stols, P. Langan, A. Joachimiak, L. Coates, A. Kovalevsky, Structural plasticity of SARS-CoV-2 3CL Mpro active site cavity revealed by room temperature X-ray crystallography, Nat. Commun. 11 (2020) 1–6. https://doi.org/10.1038/s41467-020-16954-7.

[117] A. Douangamath, D. Fearon, P. Gehrtz, T. Krojer, P. Lukacik, C.D. Owen, E. Resnick, C. Strain-Damerell, A. Aimon, P. Ábrányi-Balogh, J. Brandão-Neto, A. Carbery, G. Davison, A. Dias, T.D. Downes, L. Dunnett, M. Fairhead, J.D. Firth, S.P. Jones, A. Keeley, G.M. Keserü, H.F. Klein, M.P. Martin, M.E.M. Noble, P. O’Brien, A. Powell, R.N. Reddi, R. Skyner, M. Snee, M.J. Waring, C. Wild, N. London, F. von Delft, M.A. Walsh, Crystallographic and electrophilic fragment screening of the SARS-CoV-2 main protease, Nat. Commun. 11 (2020) 1–11. https://doi.org/10.1038/s41467-020-18709-w.

[118] D. Kuhn, N. Weskamp, S. Schmitt, E. Hüllermeier, G. Klebe, From the Similarity Analysis of Protein Cavities to the Functional Classification of Protein Families Using Cavbase, J. Mol. Biol. 359 (2006) 1023–1044. https://doi.org/10.1016/j.jmb.2006.04.024.

[119] L.S. Franco, R.C. Maia, E.J. Barreiro, Identification of LASSBio-1945 as an inhibitor of SARS-CoV-2 main protease (MPRO) through in silico screening supported by molecular docking and a fragment-based pharmacophore model, RSC Med. Chem. 12 (2021) 110–119. https://doi.org/10.1039/D0MD00282H.

[120] J. Gossen, S. Albani, A. Hanke, B.P. Joseph, C. Bergh, M. Kuzikov, E. Costanzi, C. Manelfi, P. Storici, P. Gribbon, A.R. Beccari, C. Talarico, F. Spyrakis, E. Lindahl, A. Zaliani, P. Carloni, R.C. Wade, F. Musiani, D.B. Kokh, G. Rossetti, A Blueprint for High Affinity SARS-CoV-2 Mpro Inhibitors from Activity-Based Compound Library Screening Guided by Analysis of Protein Dynamics, ACS Pharmacol. Transl. Sci. 4 (2021) 1079–1095. https://doi.org/10.1021/acsptsci.0c00215.

[121] B.L. Ho, S.C. Cheng, L. Shi, T.Y. Wang, K.I. Ho, C.Y. Chou, Critical assessment of the important residues involved in the dimerization and catalysis of MERS Coronavirus Main Protease, PLoS One. 10 (2015) e0144865. https://doi.org/10.1371/journal.pone.0144865.

[122] J. Ziebuhr, Molecular biology of severe acute respiratory syndrome coronavirus, Curr. Opin. Microbiol. 7 (2004) 412–419. https://doi.org/10.1016/j.mib.2004.06.007.

[123] M. Jukič, J. Konc, S. Gobec, D. Janežič, Identification of Conserved Water Sites in Protein Structures for Drug Design, J. Chem. Inf. Model. 57 (2017) 3094– 3103. https://doi.org/10.1021/acs.jcim.7b00443.

[124] T.A. Halgren, R.B. Murphy, R.A. Friesner, H.S. Beard, L.L. Frye, W.T. Pollard, J.L. Banks, Glide: A New Approach for Rapid, Accurate Docking and Scoring. 2. Enrichment Factors in Database Screening, J. Med. Chem. 47 (2004) 1750–1759. https://doi.org/10.1021/jm030644s.

[125] O. Trott, A.J. Olson, AutoDock Vina: Improving the speed and accuracy of docking with a new scoring function, efficient optimization, and multithreading, J. Comput. Chem. 31 (2009) NA-NA. https://doi.org/10.1002/jcc.21334.

[126] M. Miczi, M. Golda, B. Kunkli, T. Nagy, J. Tőzsér, J.A. Mótyán, Identification of host cellular protein substrates of sars-cov-2 main protease, Int. J. Mol. Sci. 21 (2020) 1–19. https://doi.org/10.3390/ijms21249523.

[127] S.L. McGovern, B.T. Helfand, B. Feng, B.K. Shoichet, A specific mechanism of nonspecific inhibition, J. Med. Chem. 46 (2003) 4265–4272. https://doi.org/10.1021/jm030266r.

[128] B.Y. Feng, B.K. Shoichet, A detergent-based assay for the detection of promiscuous inhibitors, Nat. Protoc. 1 (2006) 550–553. https://doi.org/10.1038/nprot.2006.77.

[129] A. Jadhav, R.S. Ferreira, C. Klumpp, B.T. Mott, C.P. Austin, J. Inglese, C.J. Thomas, D.J. Maloney, B.K. Shoichet, A. Simeonov, Quantitative analyses of aggregation, autofluorescence, and reactivity artifacts in a screen for inhibitors of a thiol protease, J. Med. Chem. 53 (2010) 37–51. https://doi.org/10.1021/jm901070c.

[130] S.L. McGovern, E. Caselli, N. Grigorieff, B.K. Shoichet, A common mechanism underlying promiscuous inhibitors from virtual and high-throughput screening, J. Med. Chem. 45 (2002) 1712–1722. https://doi.org/10.1021/jm010533y.

[131] P.D. Boudreau, B.W. Miller, L.I. McCall, J. Almaliti, R. Reher, K. Hirata, T. Le, J.L. Siqueira-Neto, V. Hook, W.H. Gerwick, Design of Gallinamide A Analogs as Potent Inhibitors of the Cysteine Proteases Human Cathepsin L and Trypanosoma cruzi Cruzain, J. Med. Chem. 62 (2019) 9026–9044. https://doi.org/10.1021/acs.jmedchem.9b00294.

[132] R. Hilgenfeld, From SARS to MERS: crystallographic studies on coronaviral proteases enable antiviral drug design, FEBS J. 281 (2014) 4085–4096. https://doi.org/10.1111/febs.12936.

[133] J.B. Baell, G.A. Holloway, New substructure filters for removal of pan assay interference compounds (PAINS) from screening libraries and for their exclusion in bioassays, J. Med. Chem. 53 (2010) 2719–2740. https://doi.org/10.1021/jm901137j.

[134] J. Lee, L.J. Worrall, M. Vuckovic, F.I. Rosell, F. Gentile, A.T. Ton, N.A. Caveney, F. Ban, A. Cherkasov, M. Paetzel, N.C.J. Strynadka, Crystallographic structure of wild-type SARS-CoV-2 main protease acyl-enzyme intermediate with physiological C-terminal autoprocessing site, Nat. Commun. 11 (2020) 1–9. https://doi.org/10.1038/s41467-020-19662-4.

[135] R. Hilgenfeld, K. Anand, J.R. Mesters, Z. Rao, X. Shen, H. Jiang, J. Tan, K.H.G. Verschueren, Structure and dynamics of SARS coronavirus main proteinase (M pro), in: Adv. Exp. Med. Biol., Springer, 2006: pp. 585–591. https://doi.org/10.1007/978-0-387-33012-9_106.

[136] B. Goyal, D. Goyal, Targeting the Dimerization of the Main Protease of Coronaviruses: A Potential Broad-Spectrum Therapeutic Strategy, ACS Comb. Sci. 22 (2020) 297–305. https://doi.org/10.1021/acscombsci.0c00058.

[137] S. Chen, L. Chen, J. Tan, J. Chen, L. Du, T. Sun, J. Shen, K. Chen, H. Jiang, X. Shen, Severe acute respiratory syndrome coronavirus 3C-like proteinase N terminus is indispensable for proteolytic activity but not for enzyme dimerization: Biochemical and thermodynamic investigation in conjunction with molecular dynamics simulations, J. Biol. Chem. 280 (2005) 164–173. https://doi.org/10.1074/jbc.M408211200.

[138] D. Suárez, N. Díaz, SARS-CoV-2 main protease: a molecular dynamics study, J. Chem. Inf. Model. (2020).

[139] S. Hattori, N. Higashi-Kuwata, H. Hayashi, S.R. Allu, J. Raghavaiah, H. Bulut, C. Das, B.J. Anson, E.K. Lendy, Y. Takamatsu, N. Takamune, N. Kishimoto, K. Murayama, K. Hasegawa, M. Li, D.A. Davis, E.N. Kodama, R. Yarchoan, A. Wlodawer, S. Misumi, A.D. Mesecar, A.K. Ghosh, H. Mitsuya, A small molecule compound with an indole moiety inhibits the main protease of SARS-CoV-2 and blocks virus replication, Nat. Commun. 12 (2021) 1–12. https://doi.org/10.1038/s41467-021-20900-6.

[140] W.M. Singh, J.B. Baruah, Synthesis of mixed aryl 2,3-diarylsulphanyl-1,4-naphthoquinones, Synth. Commun. 39 (2009) 1433–1442. https://doi.org/10.1080/00397910802528951.

[141] J. Qiao, Y.S. Li, R. Zeng, F.L. Liu, R.H. Luo, C. Huang, Y.F. Wang, J. Zhang, B. Quan, C. Shen, X. Mao, X. Liu, W. Sun, W. Yang, X. Ni, K. Wang, L. Xu, Z.L. Duan, Q.C. Zou, H.L. Zhang, W. Qu, Y.H.P. Long, M.H. Li, R.C. Yang, X. Liu, J. You, Y. Zhou, R. Yao, W.P. Li, J.M. Liu, P. Chen, Y. Liu, G.F. Lin, X. Yang, J. Zou, L. Li, Y. Hu, G.W. Lu, W.M. Li, Y.Q. Wei, Y.T. Zheng, J. Lei, S. Yang, SARS-CoV-2 Mpro inhibitors with antiviral activity in a transgenic mouse model, Science (80-.). 371 (2021) 1374–1378. https://doi.org/10.1126/science.abf1611originally.

[142] D.J.B. Lima, R.G. Almeida, G.A.M. Jardim, B.P.A. Barbosa, A.C.C. Santos, W.O. Valença, M.R. Scheide, C.C. Gatto, G.G.C. de Carvalho, P.M.S. Costa, C. Pessoa, C.L.M. Pereira, C. Jacob, A.L. Braga, E.N. da Silva Júnior, It takes two to tango: synthesis of cytotoxic quinones containing two redox active centers with potential antitumor activity, RSC Med. Chem. (2021). https://doi.org/10.1039/d1md00168j.

[143] H.M. Berman, J. Westbrook, Z. Feng, G. Gilliland, T.N. Bhat, H. Weissig, I.N. Shindyalov, P.E. Bourne, The Protein Data Bank, Nucleic Acids Res. 28 (2000) 235–242. https://doi.org/10.1093/nar/28.1.235.

[144] Core R Team, A Language and Environment for Statistical Computing, R Found. Stat. Comput. 2 (2019) https://www.R--project.org. http://www.r-project.org.

[145] B.J. Grant, A.P.C. Rodrigues, K.M. ElSawy, J.A. McCammon, L.S.D. Caves, Bio3d: an R package for the comparative analysis of protein structures, Bioinformatics. 22 (2006) 2695–2696.

[146] W.L. Delano, The PyMOL Molecular Graphics System, (2002). http://www.pymol.org.

[147] K. Zhu, K.W. Borrelli, J.R. Greenwood, T. Day, R. Abel, R.S. Farid, E. Harder, Docking covalent inhibitors: A parameter free approach to pose prediction and scoring, J. Chem. Inf. Model. 54 (2014) 1932–1940. https://doi.org/10.1021/ci500118s.

[148] W. Sherman, T. Day, M.P. Jacobson, R.A. Friesner, R. Farid, Novel procedure for modeling ligand/receptor induced fit effects, J. Med. Chem. 49 (2006) 534– 553. https://doi.org/10.1021/jm050540c.

[149] K.J. Bowers, D.E. Chow, H. Xu, R.O. Dror, M.P. Eastwood, B.A. Gregersen, J.L. Klepeis, I. Kolossvary, M.A. Moraes, F.D. Sacerdoti, J.K. Salmon, Y. Shan, D.E. Shaw, Scalable Algorithms for Molecular Dynamics Simulations on Commodity Clusters, in: SC ’06 Proc. 2006 ACM/IEEE Conf. Supercomput., 2007: pp. 43–43. https://doi.org/10.1109/sc.2006.54.

[150] E. Harder, W. Damm, J. Maple, C. Wu, M. Reboul, J.Y. Xiang, L. Wang, D. Lupyan, M.K. Dahlgren, J.L. Knight, J.W. Kaus, D.S. Cerutti, G. Krilov, W.L. Jorgensen, R. Abel, R.A. Friesner, OPLS3: A Force Field Providing Broad Coverage of Drug-like Small Molecules and Proteins, J. Chem. Theory Comput. 12 (2016) 281–296. https://doi.org/10.1021/acs.jctc.5b00864.

[151] G.M. Ferreira, T. Kronenberger, A.K. Tonduru, R.D.C. Hirata, M.H. Hirata, A. Poso, SARS-COV-2 Mpro conformational changes induced by covalently bound ligands, J. Biomol. Struct. Dyn. (2021) 1–11. https://doi.org/10.1080/07391102.2021.1970626.

[152] W.L. Jorgensen, J. Chandrasekhar, J.D. Madura, R.W. Impey, M.L. Klein, Comparison of simple potential functions for simulating liquid water, J. Chem. Phys. 79 (1983) 926–935. https://doi.org/10.1063/1.445869.

[153] T. Darden, D. York, L. Pedersen, Particle mesh Ewald: An N·log(N) method for Ewald sums in large systems, J. Chem. Phys. 98 (1993) 10089–10092. https://doi.org/10.1063/1.464397.

[154] A.S. Ashhurst, A.H. Tang, P. Fajtová, M. Yoon, A. Aggarwal, A. Stoye, M. Larance, L. Beretta, A. Drelich, D. Skinner, L. Li, T.D. Meek, J.H. McKerrow, V. Hook, C.-T.K. Tseng, S. Turville, W.H. Gerwick, A.J. O’Donoghue, R.J. Payne, Potent in vitro anti-SARS-CoV-2 activity by gallinamide A and analogues via inhibition of cathepsin L., BioRxiv Prepr. Serv. Biol. (2020). https://doi.org/10.1101/2020.12.23.424111.

