## Supplementary Material for "Structure-based identification of naphthoquinones and derivatives as novel inhibitors of main protease Mpro and papain-like protease PLpro of SARS-CoV-2"

### 1. Assembly of a chemical library for virtual screening

#### 1.1 General Information

The compounds studied in this work were selected according to our previous experience in the synthesis of quinoidal structures and their derivatives. In this sense, the structures evaluated by computational methods have been previously synthesized and described in the literature. We divided the substances into eight groups of molecules according to their structural features. Group 1 are *ortho*-quinones with different substitution patterns. We studied compounds containing arylamino (**1-20**),[1,2,3,4,5,6] alcohol (**21**)[2] and alkoxy groups (**22-25**),[3,4] selenium (**26-36** and **42-57**)[7] and sulfur (**37-41** and **58-61**),[8,9,10] the basic chalcone framework (**62-67**),[11] among simple *ortho*-quinones (**69-75**).[3,11,12,13,14] Group 2 are *para*-quinones with different substitution patterns. We studied compounds of α-lapachones (**77** and **78**)[2,15] arylamino (**79-86**),[2,6,8] furanonaphthoquinone (**87-95** and **137-154**),[16,17,18] pyrrolonaphthoquinones (**96-136**),[17,19] other derivatives of *para*-quinones (**155-158**).[6,17] Group 3 are *ortho*-quinones-based 1,2,3-triazoles with different substitution patterns. We studied compounds with aromatic and aliphatic substituents (**159-174**, **189-197, 202-207, 217-223, 226-231, 235-241** and **258-260**)[3,8,12,20,21,22,23,24,25,26] selenium (**175-184, 208-215, 224, 225, 232** and **242-257**)[6,12,27] and sulfur (**185** and **216**)[6,27] carbohydrates (**186-188, 233** and **234**),[8] *para*-quinones (**189-191**),[28] bodipy (**198-201**).[29,30] Group 4 are *para*-quinones-based 1,2,3-triazoles with different substitution patterns. We studied triazoles derived from lapachol (**261-291, 322-324** and **333-345**),[6,27,29,30,31,32] arylamino (**292-321**)[27,33,34] and *N*-sulfonyl (**325-332**).[10] Group 5 are phenazinic with different substitution patterns. We studied triazoles derived of phenazinic (**346-355, 357** and **367-371**),[25,27,35,36,37] π-extended phenazines (**358-360** and **362-366**),[37] quinone (**356** and **372**)[35] and simple phenazines (**361** and **373**).[14,35,36,38] Group 6 are *para*-quinones and derived with diferrent substituition patterns. We studied sulfur (**379-384, 458-485, 558-561** and **593-604**)[39,40,41,42] and selenium compounds (**486-507** and **551-557**)[,43] iodo (**385-405** and **608-612**),[44] arylamino (**406-428** and **587-592**),[19,34] benzoquinones (**605-607** and **613-616**),[45] imine (**533-540**)[10] and other derivatives (**429-547**, **508-532**, **541-550**, **562-586** and **617-659**).[6,8,15,17,29, 46,47,48,49,50,51,52,53,54] Group 7 are hydrazo derivatives. We studied compounds of *para*-quinones with diferrent substituition patterns (**660-665**)[11,55] and *ortho*-quinones (**666-676**).[11,55] Group 8 are imidazole and oxazole with diferrent substituition patterns. We studied compounds imidazoles (**667-669**),[56] and oxazoles (**680-688**).[25,35,57,58]

#### 1.2 Group 1: *Ortho*-quinones

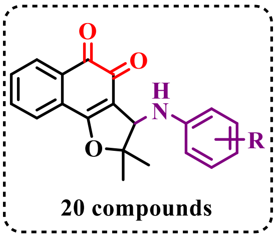

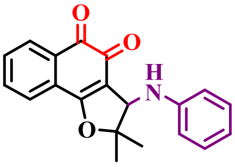

**(1)**

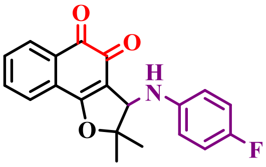

**(2)**

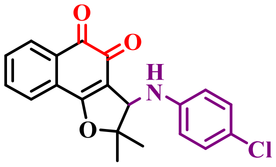

**(3)**

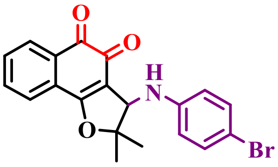

**(4)**

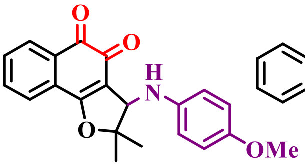

**(5)**

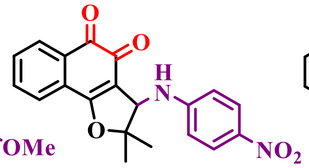

**(6)**

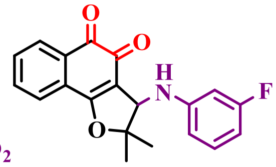

**(7)**

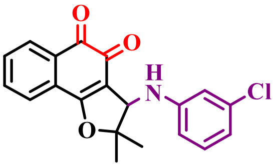

**(8)**

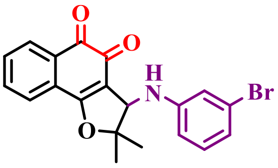

**(9)**

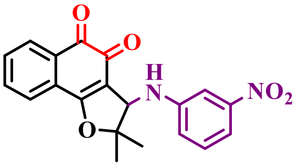

**(10)**

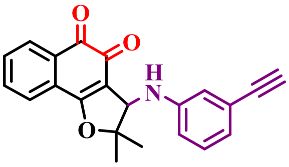

**(11)**

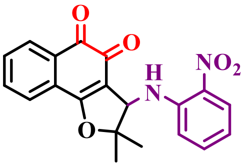

**(12)**

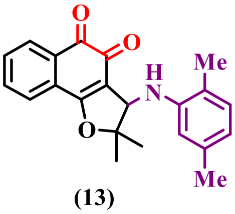

**(13)**

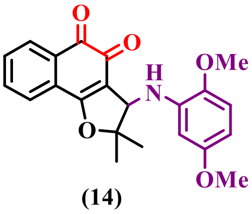

**(14)**

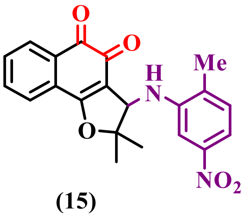

**(15)**

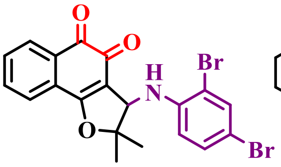

**(16)**

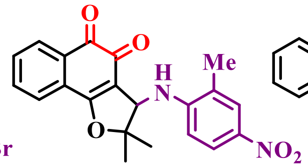

**(17)**

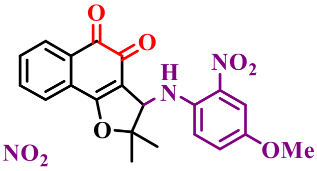

**(18)**

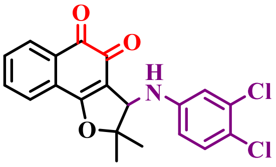

**(19)**

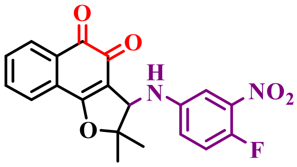

**(20)**

**Figure S1.** Group 1: *Ortho*-quinones (part I).

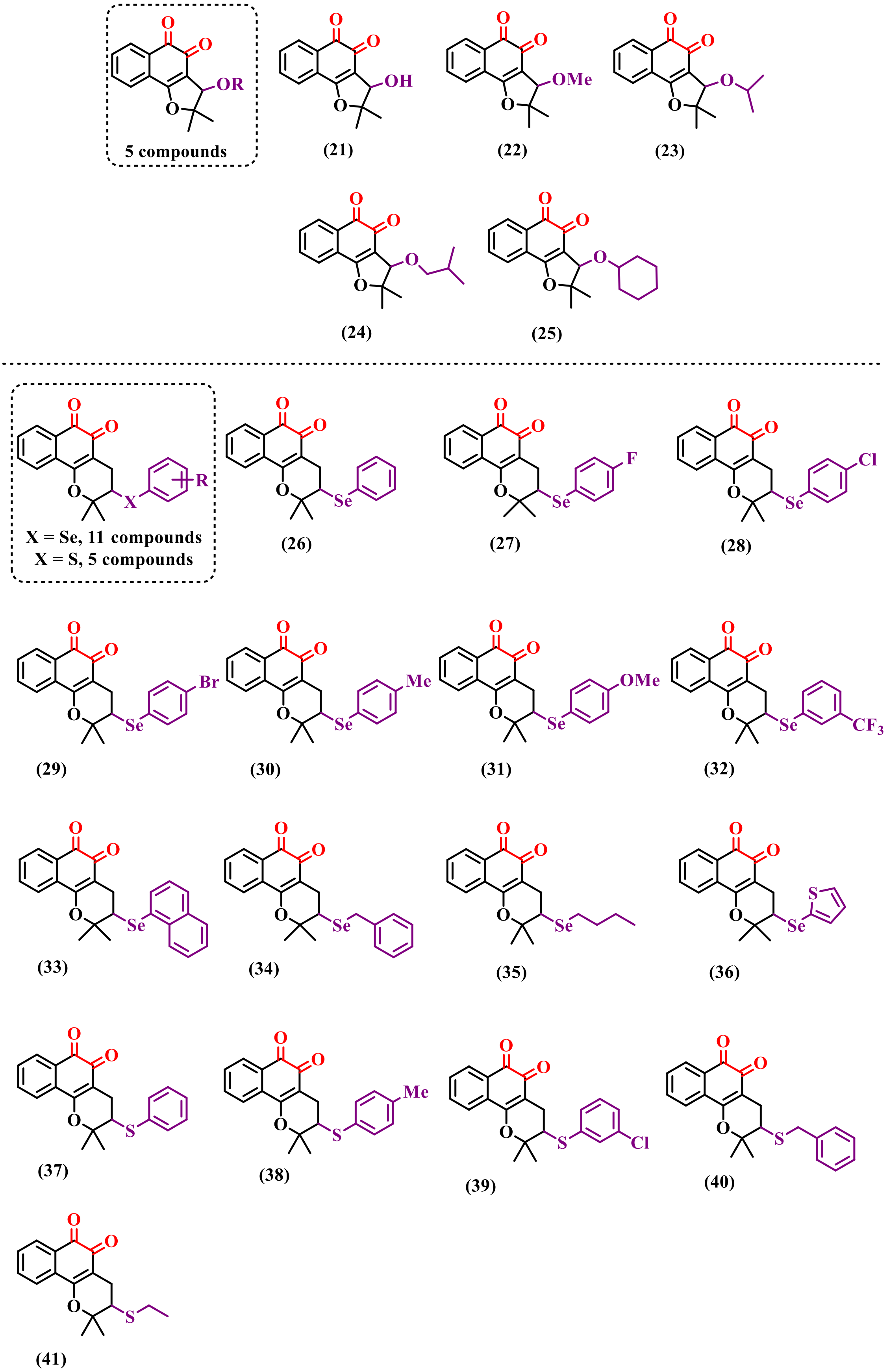

**Figure S2.** Group 1: *Ortho*-quinones (part II).

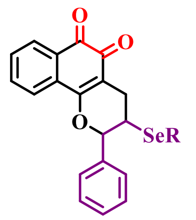

9 compounds

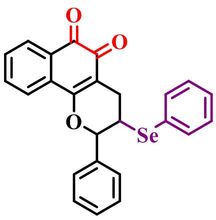

(42)

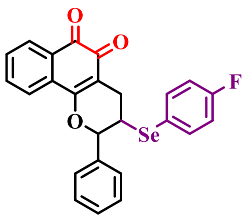

(43)

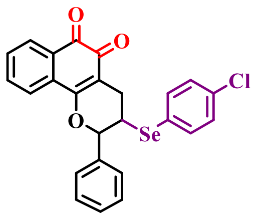

(44)

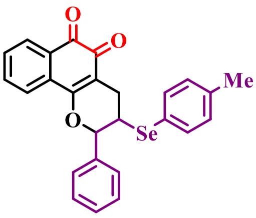

(45)

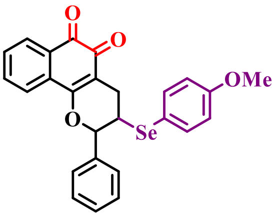

(46)

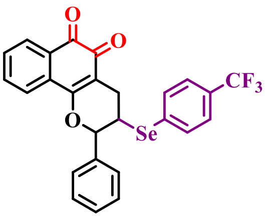

(47)

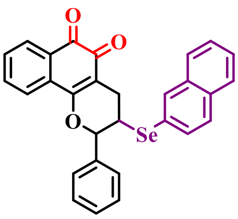

(48)

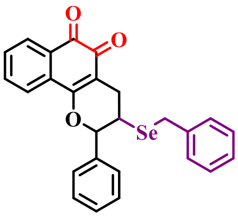

(49)

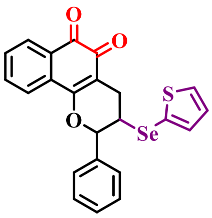

(50)

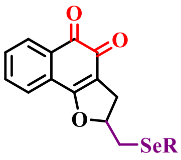

7 compounds

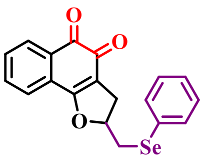

(51)

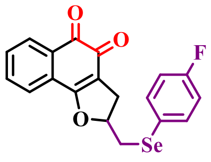

(52)

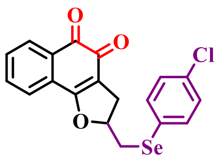

(53)

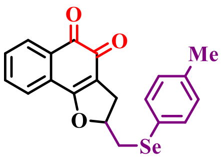

(54)

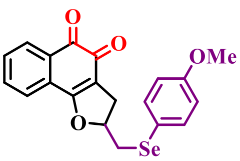

(55)

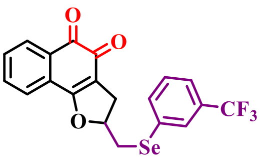

(56)

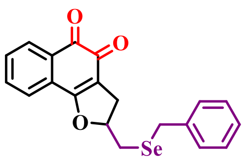

(57)

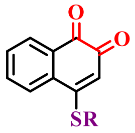

4 compounds

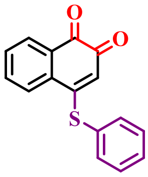

(58)

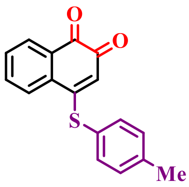

(59)

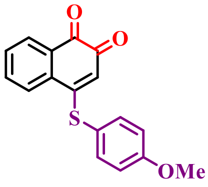

(60)

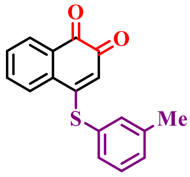

(61)

**Figure S3.** Group 1: Ortho-quinones (part III).

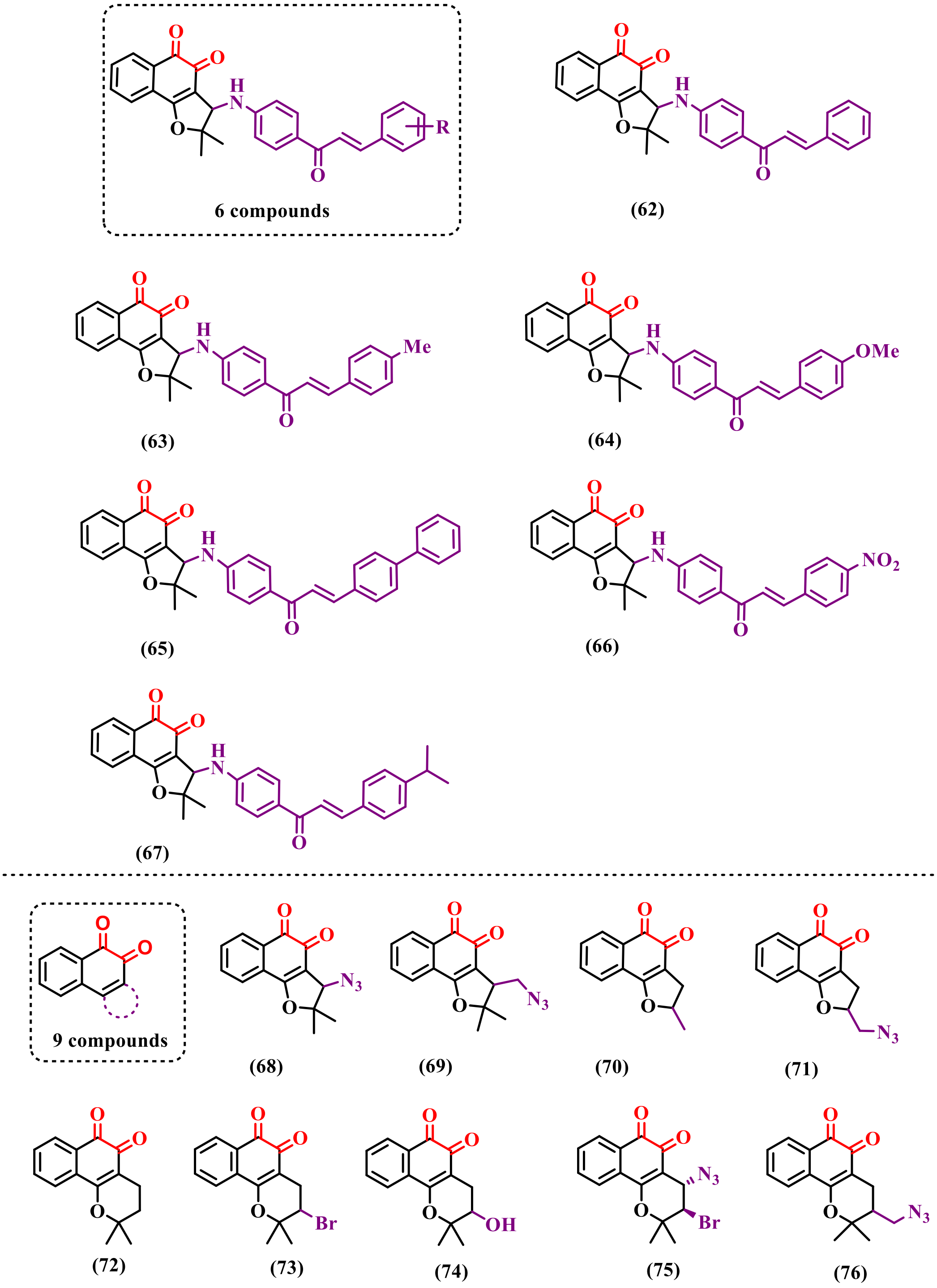

**Figure S4.** Group 1: *Ortho*-quinones (part IV).

#### 1.3 Group 2: *Para*-quinones

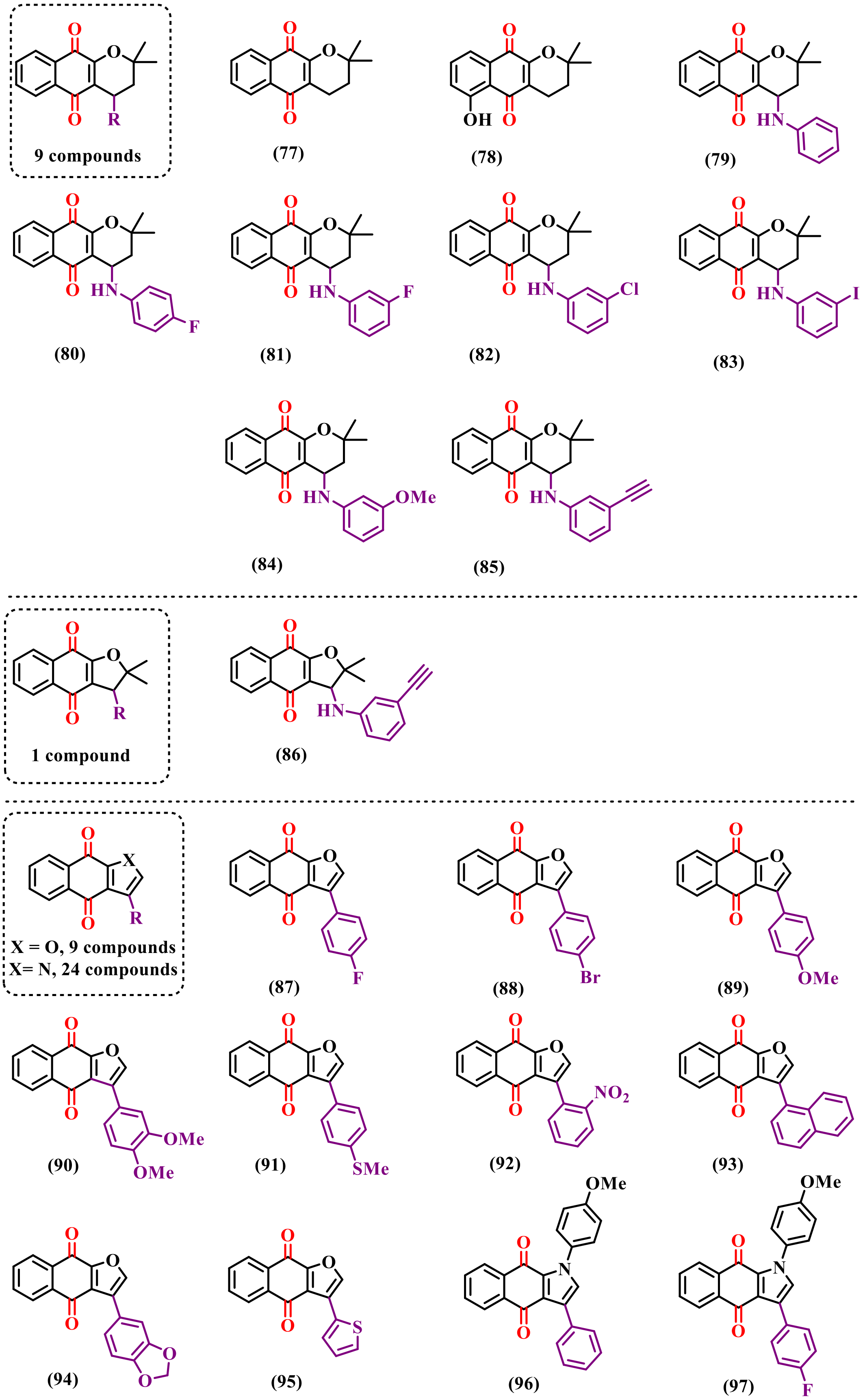

**Figure S5.** Group 2: *Para*-quinones (part I).

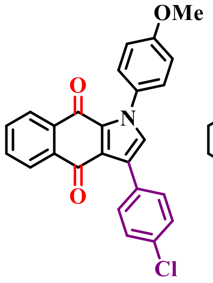

**(98)**

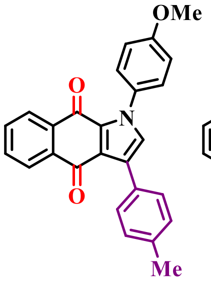

**(99)**

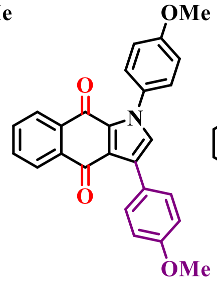

**(100)**

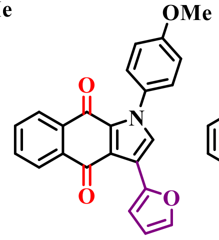

**(101)**

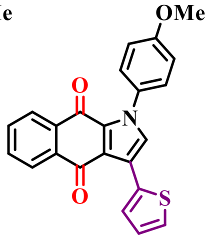

**(102)**

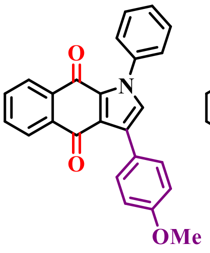

**(103)**

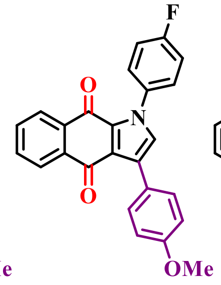

**(104)**

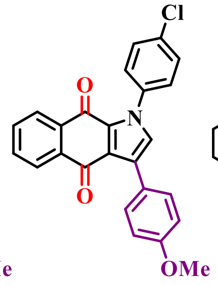

**(105)**

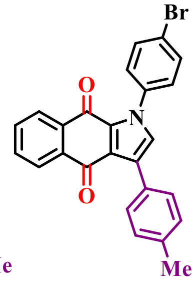

**(106)**

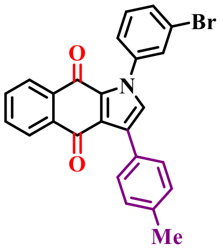

**(107)**

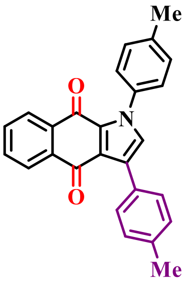

**(108)**

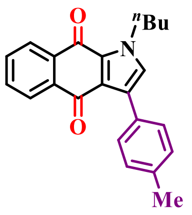

**(109)**

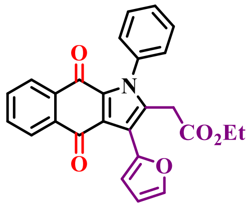

**(110)**

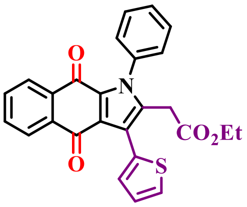

**(111)**

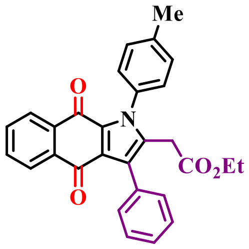

**(112)**

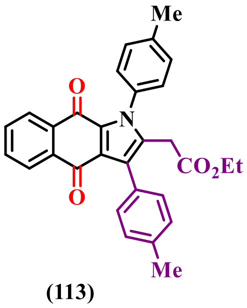

**(113)**

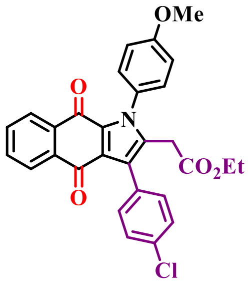

**(114)**

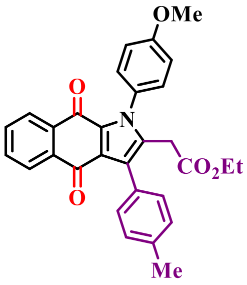

**(115)**

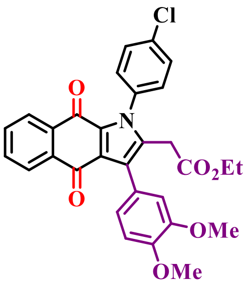

**(116)**

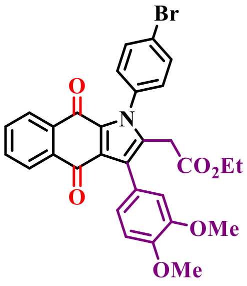

**(117)**

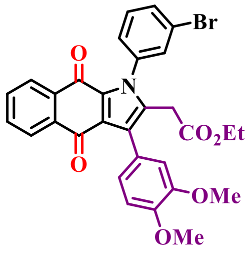

**(118)**

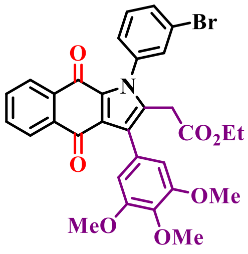

**(119)**

**Figure S6.** Group 2: *Para*-quinones (part II).

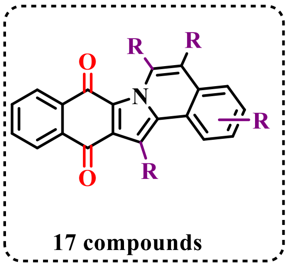

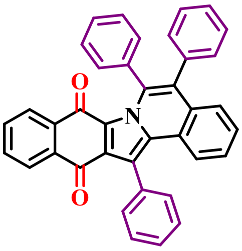

(120)

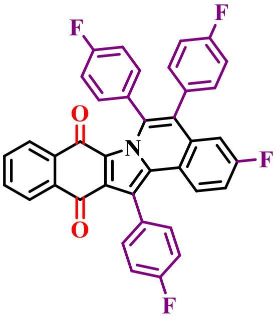

(121)

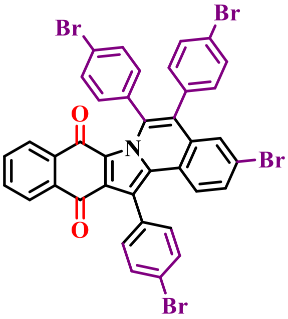

(122)

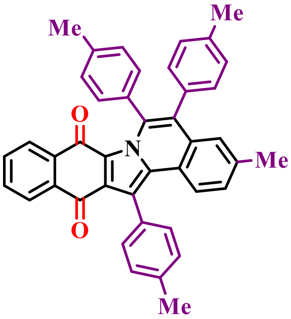

(123)

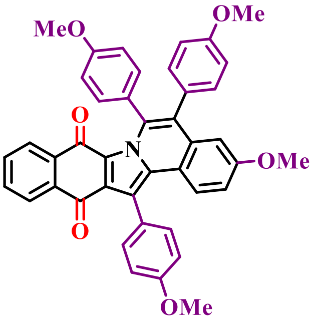

(124)

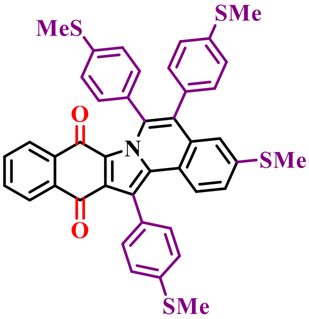

(125)

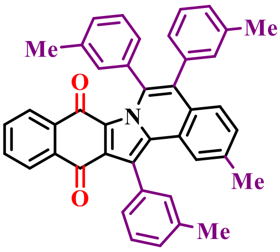

(126)

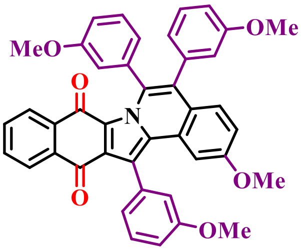

(127)

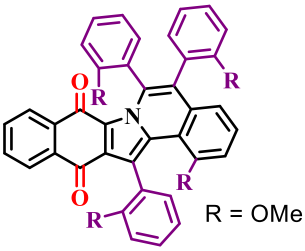

(128)

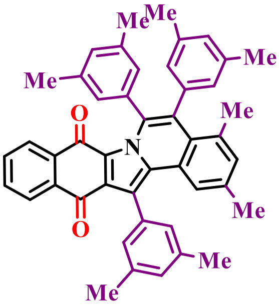

(129)

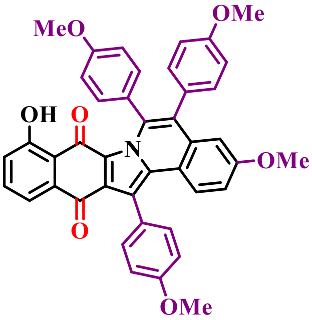

(130)

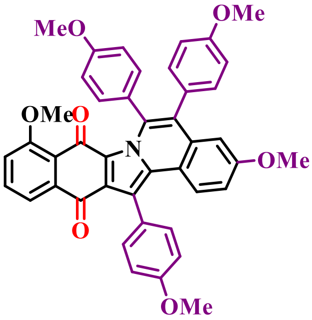

(131)

**Figure S7.** Group 2: Para-quinones (part III).

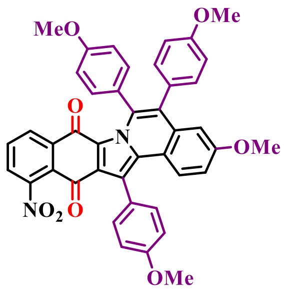

(132)

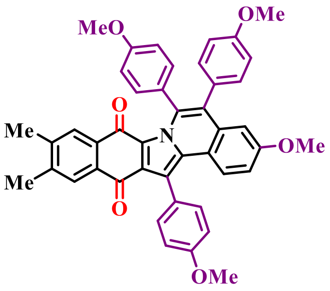

(133)

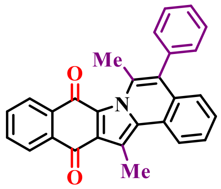

(134)

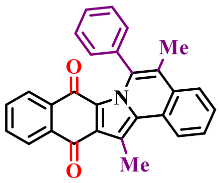

(135)

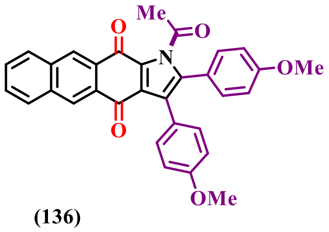

(136)

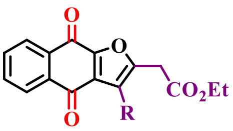

10 compounds

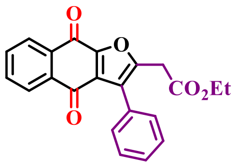

(137)

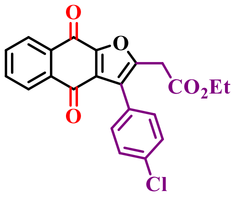

(138)

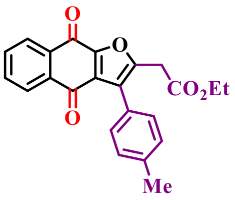

(139)

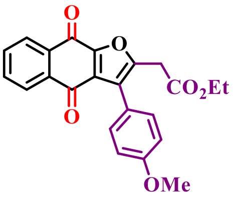

(140)

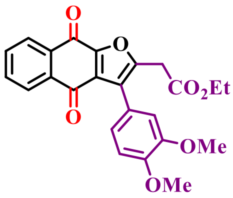

(141)

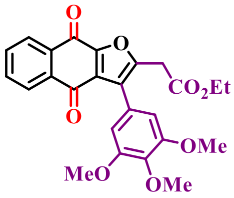

(142)

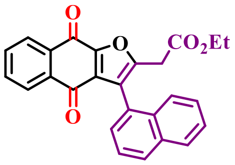

(143)

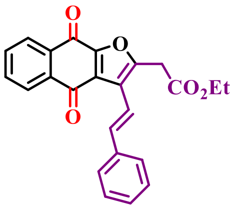

(144)

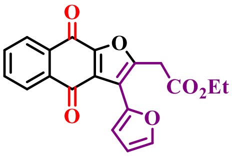

(145)

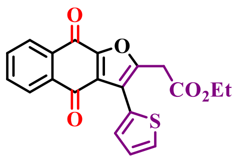

(146)

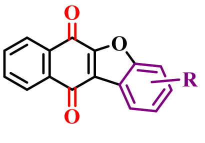

8 compounds

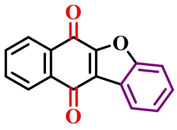

(147)

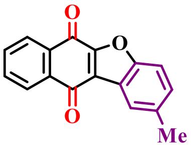

(148)

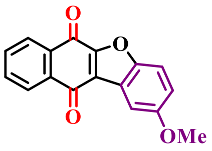

(149)

**Figure S8.** Group 2: Para-quinones (part IV).

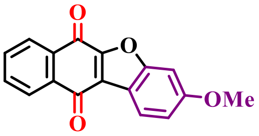

**(150)**

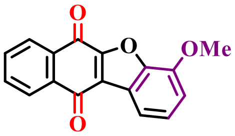

**(151)**

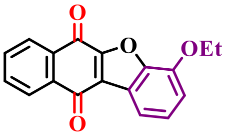

**(152)**

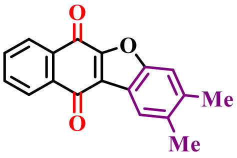

**(153)**

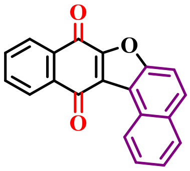

**(154)**

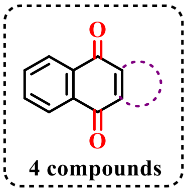

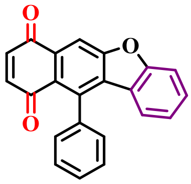

**(155)**

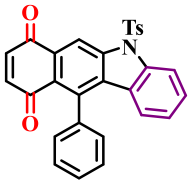

**(156)**

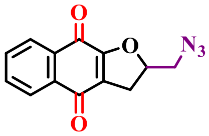

**(157)**

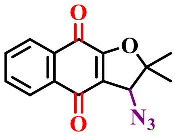

**(158)**

**Figure S9.** Group 2: *Para*-quinones (part V).

#### 1.4 Group 3: *Ortho*-quinones-based 1,2,3-triazoles

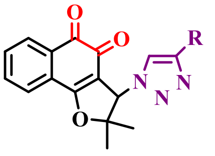

44 compounds

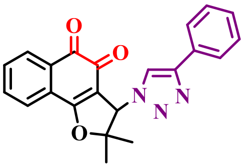

(159)

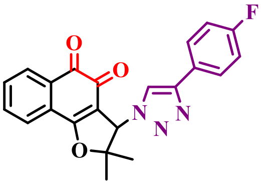

(160)

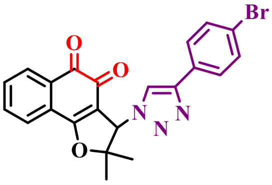

(161)

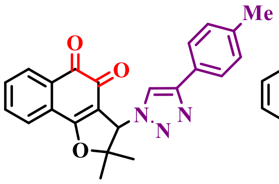

(162)

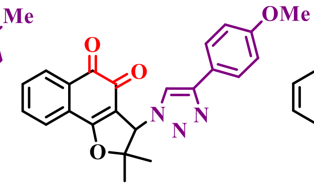

(163)

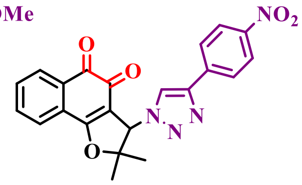

(164)

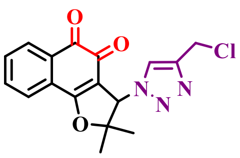

(165)

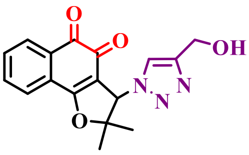

(166)

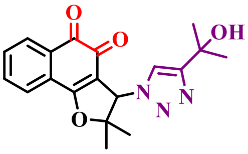

(167)

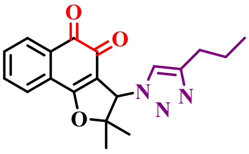

(168)

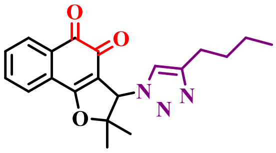

(169)

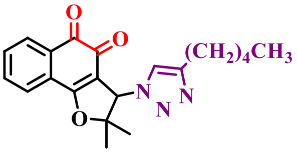

(170)

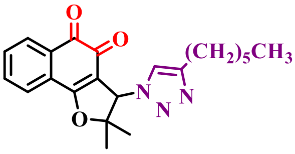

(171)

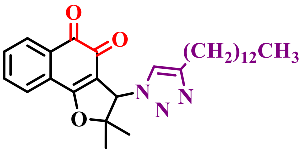

(172)

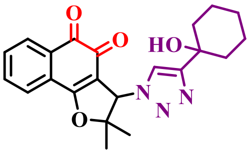

(173)

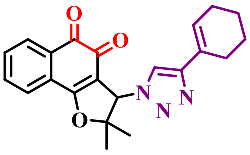

(174)

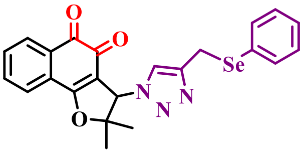

(175)

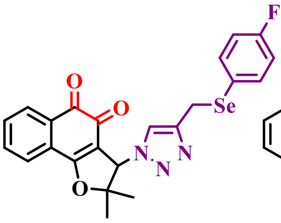

(176)

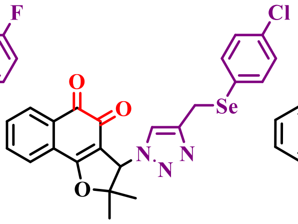

(177)

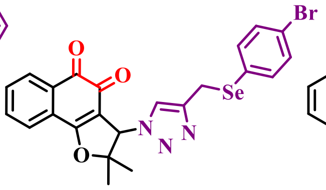

(178)

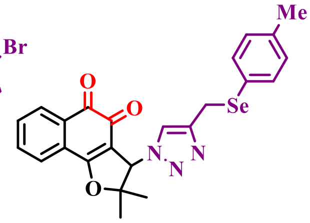

(179)

**Figure S10.** Group 3: *Ortho*-quinones-based 1,2,3-triazoles (part I).

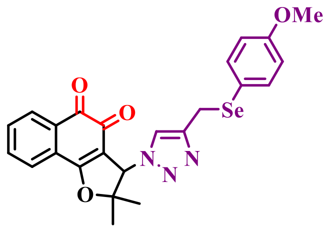

**(180)**

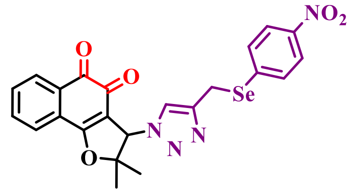

**(181)**

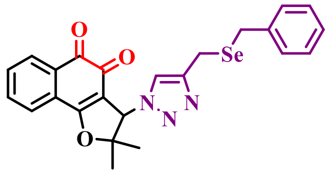

**(182)**

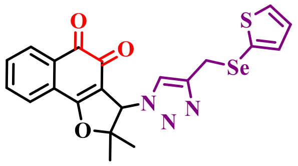

**(183)**

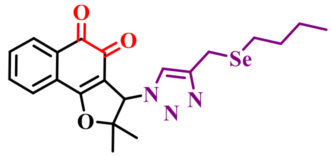

**(184)**

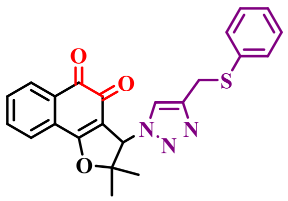

**(185)**

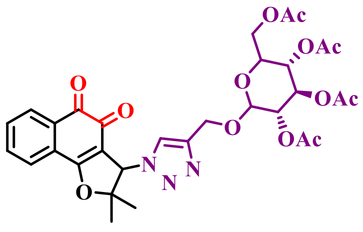

**(186)**

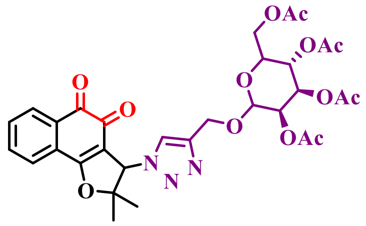

**(187)**

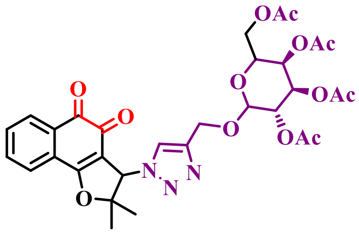

**(188)**

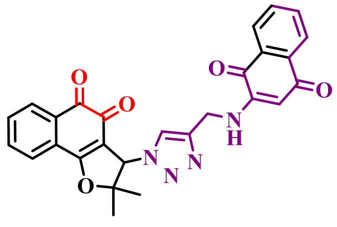

**(189)**

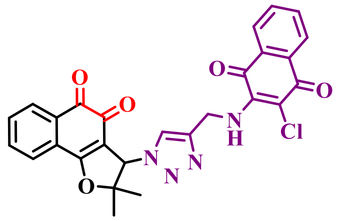

**(190)**

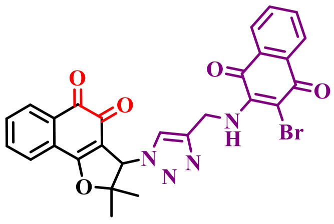

**(191)**

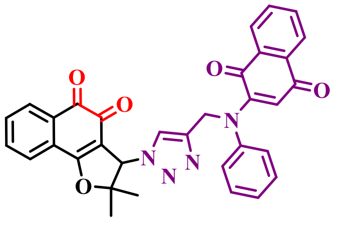

**(192)**

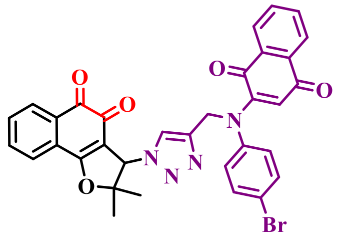

**(193)**

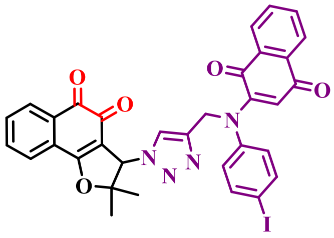

**(194)**

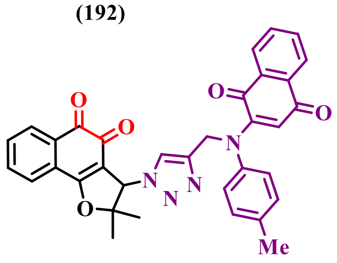

**(195)**

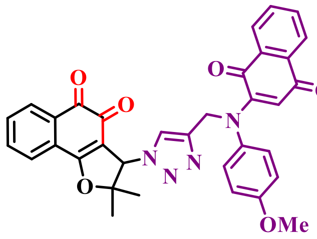

**(196)**

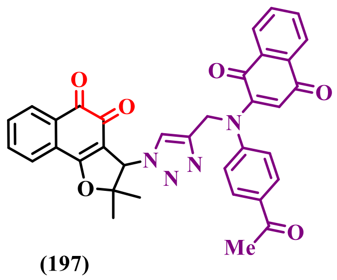

**(197)**

**Figure S11.** Group 3: *Ortho*-quinones-based 1,2,3-triazoles (part II).

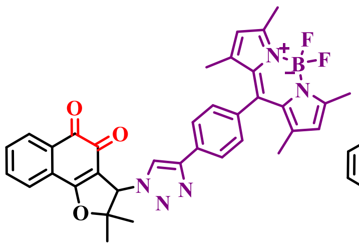

**(198)**

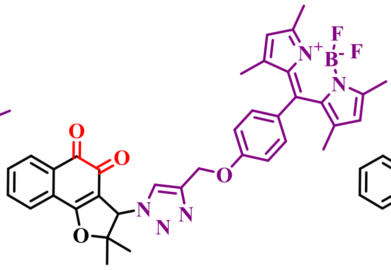

**(199)**

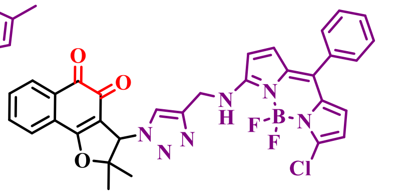

**(200)**

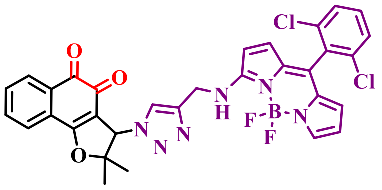

**(201)**

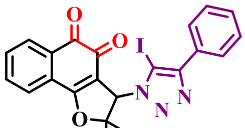

**(202)**

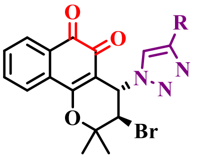

**21 compounds**

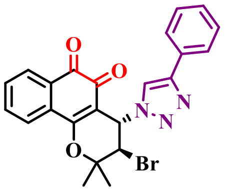

**(203)**

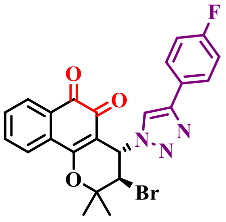

**(204)**

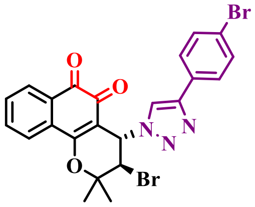

**(205)**

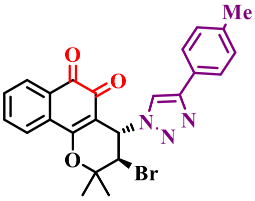

**(206)**

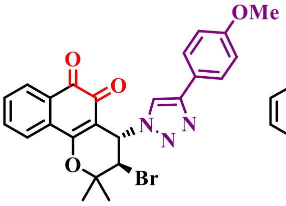

**(207)**

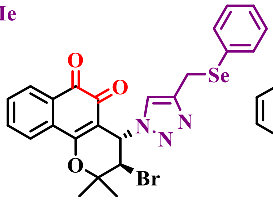

**(208)**

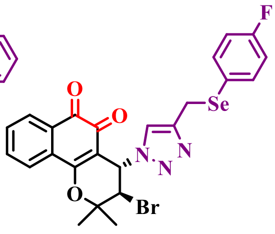

**(209)**

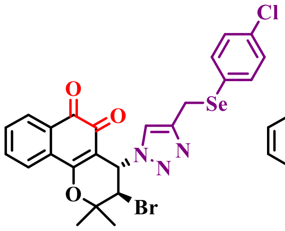

**(210)**

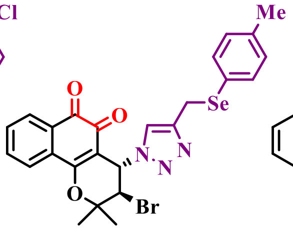

**(211)**

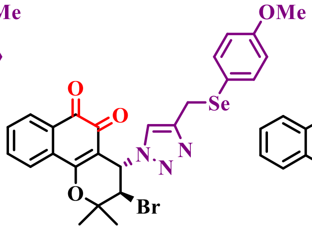

**(212)**

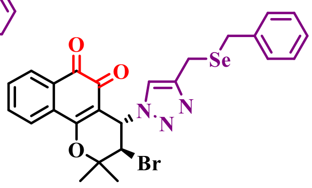

**(213)**

**Figure S12.** Group 3: *Ortho*-quinones-based 1,2,3-triazoles (part III).

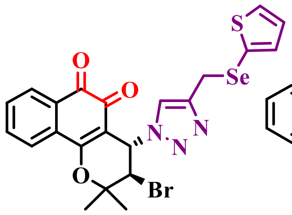

**(214)**

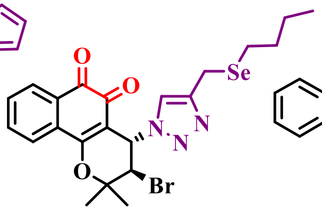

**(215)**

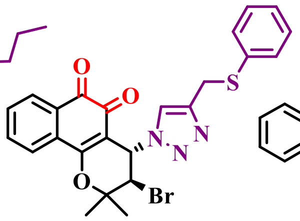

**(216)**

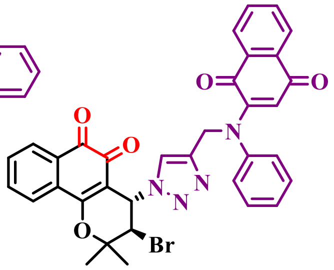

**(217)**

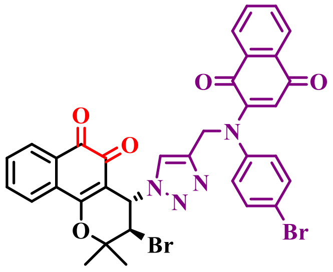

**(218)**

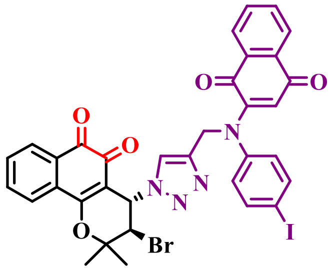

**(219)**

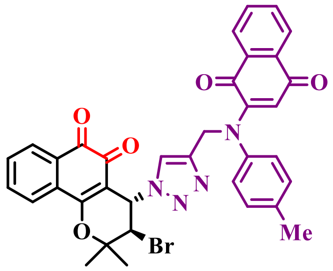

**(220)**

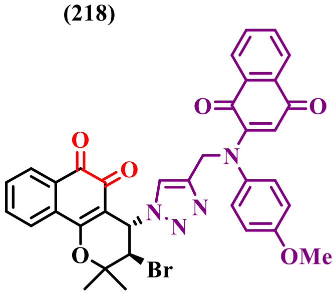

**(221)**

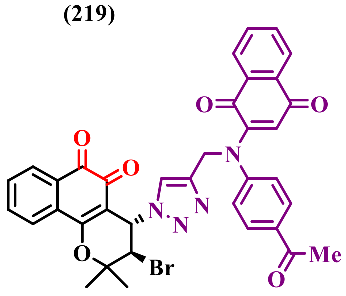

**(222)**

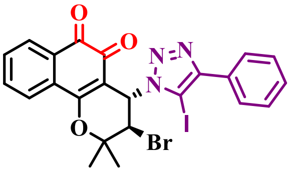

**(223)**

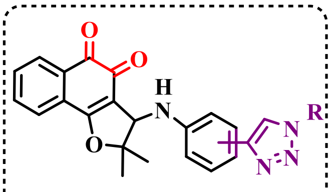

**2 compounds**

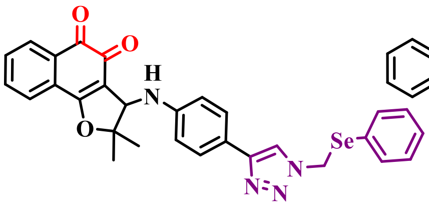

**(224)**

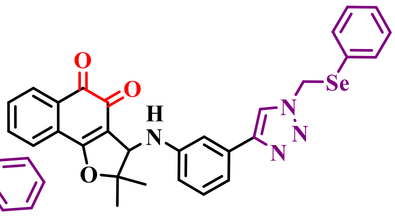

**(225)**

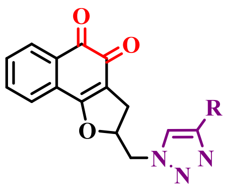

**16 compounds**

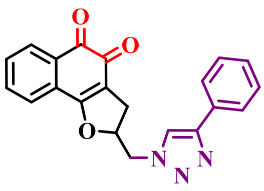

**(226)**

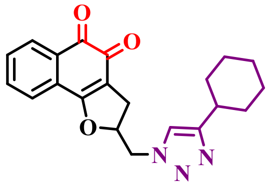

**(227)**

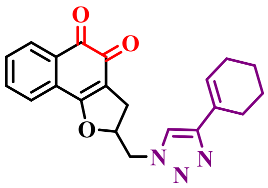

**(228)**

**Figure S13.** Group 3: *Ortho*-quinones-based 1,2,3-triazoles (part IV).

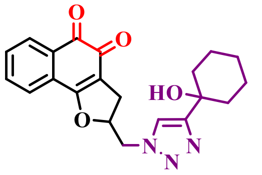

**(229)**

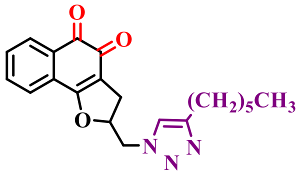

**(230)**

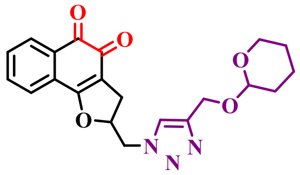

**(231)**

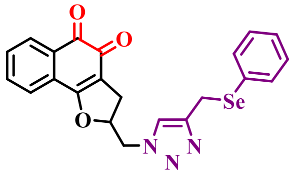

**(232)**

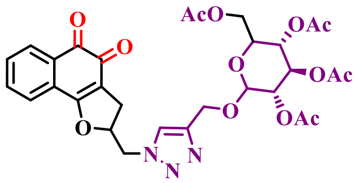

**(233)**

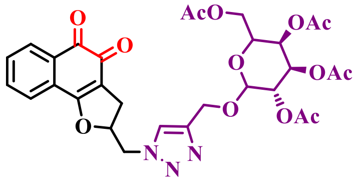

**(234)**

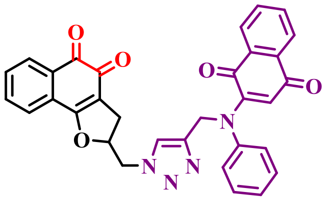

**(235)**

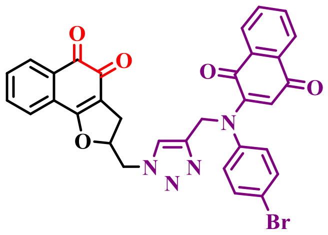

**(236)**

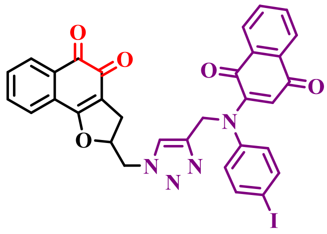

**(237)**

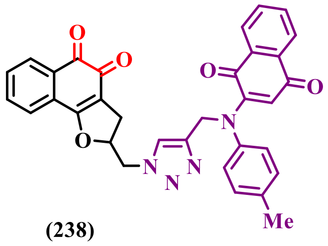

**(238)**

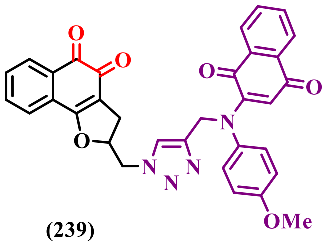

**(239)**

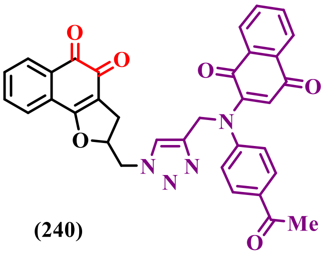

**(240)**

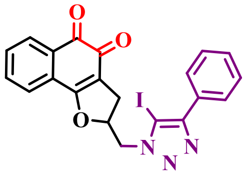

**(241)**

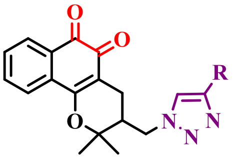

**8 compounds**

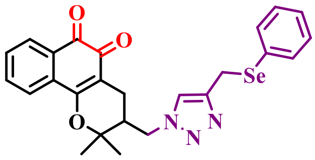

**(242)**

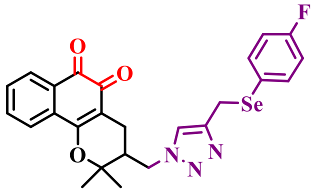

**(243)**

**Figure S14.** Group 3: *Ortho*-quinones-based 1,2,3-triazoles (part V).

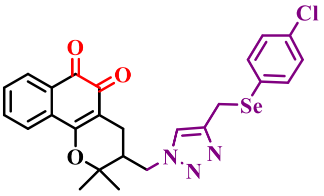

**(244)**

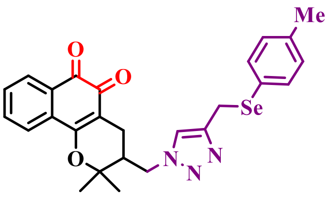

**(245)**

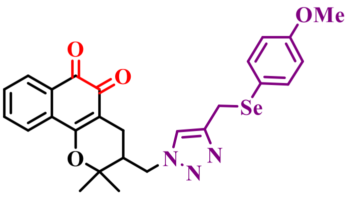

**(246)**

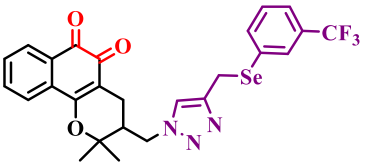

**(247)**

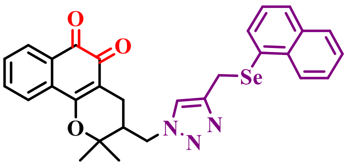

**(248)**

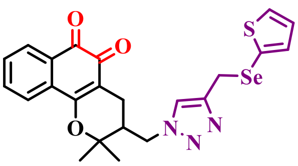

**(249)**

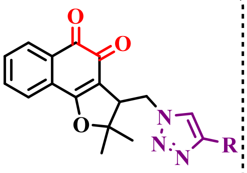

**8 compounds**

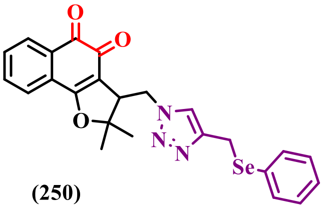

**(250)**

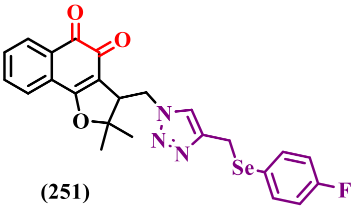

**(251)**

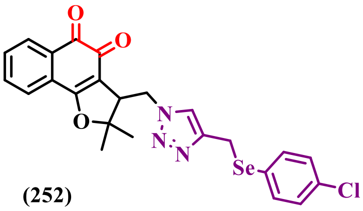

**(252)**

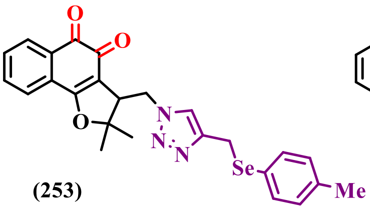

**(253)**

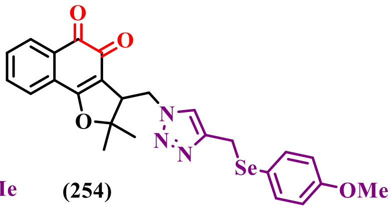

**(254)**

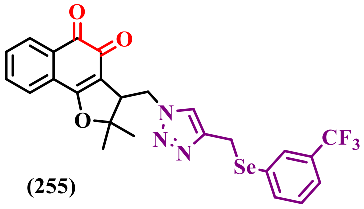

**(255)**

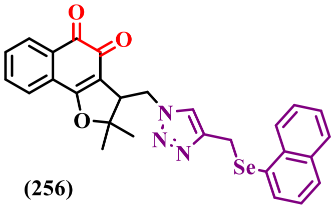

**(256)**

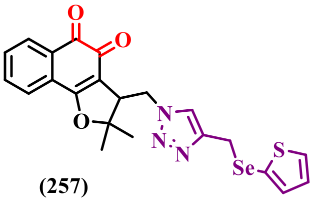

**(257)**

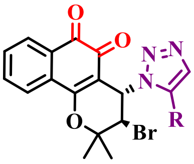

**1 compound**

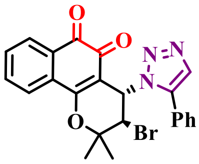

**(258)**

**Figure S15.** Group 3: *Ortho*-quinones-based 1,2,3-triazoles *(part VI)*.

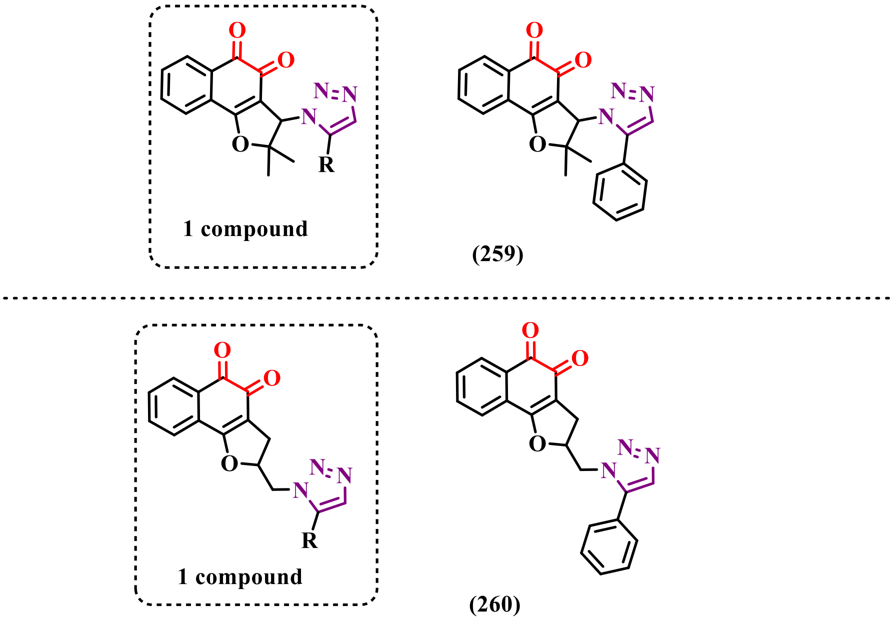

**Figure S16.** Group 3: *Ortho*-quinones-based 1,2,3-triazoles (part VII).

#### 1.5 Group 4: *Para*-quinones-based 1,2,3-triazoles

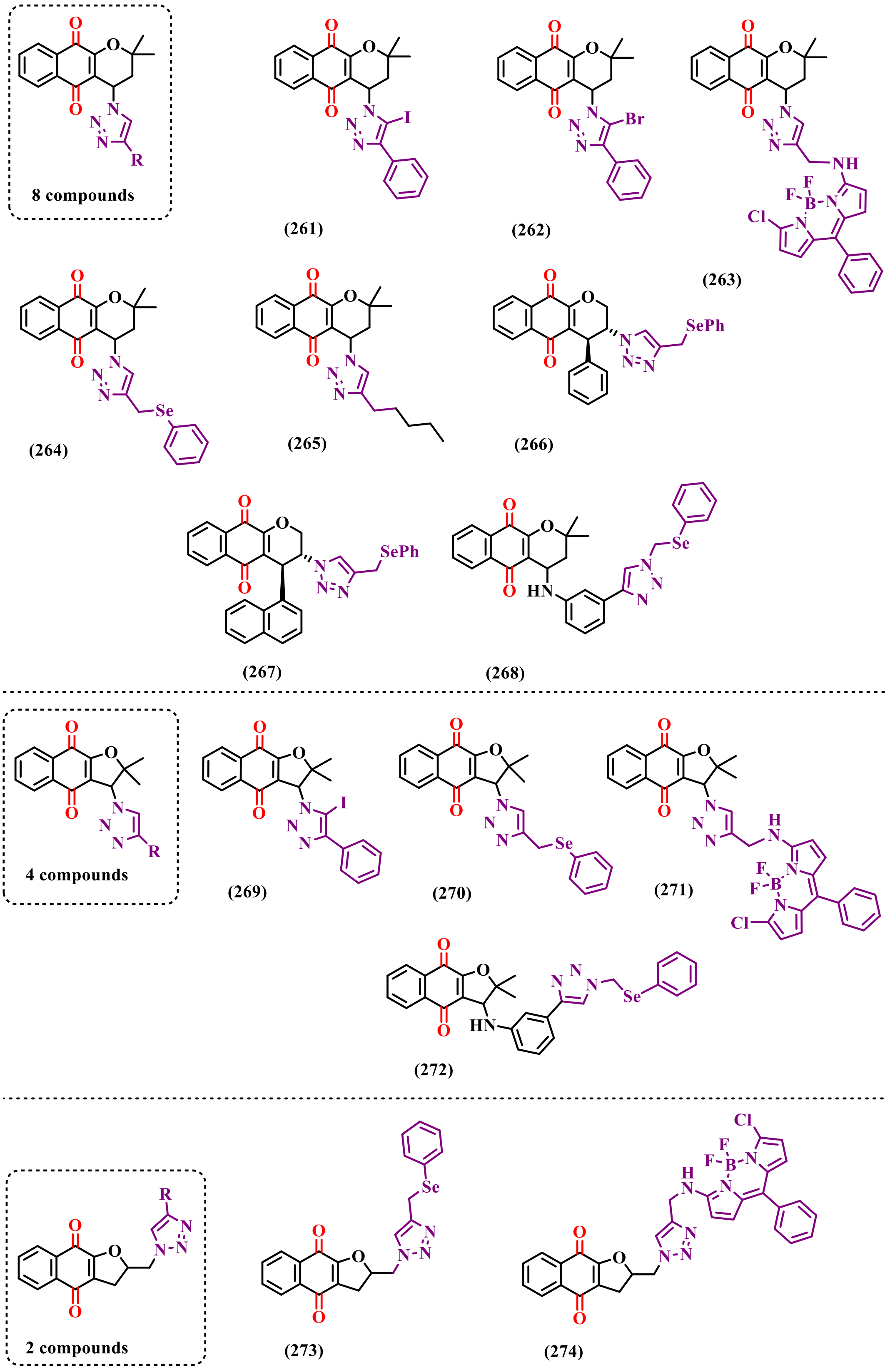

**Figure S17.** Para-quinones-based 1,2,3-triazoles (part I).

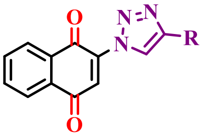

17 compounds

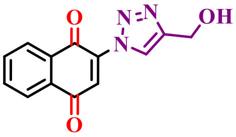

**(275)**

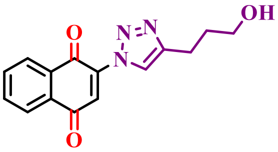

**(276)**

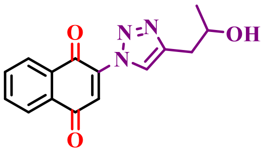

**(277)**

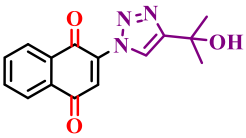

**(278)**

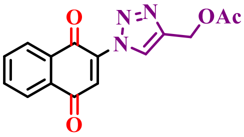

**(279)**

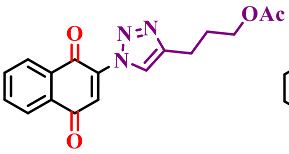

**(280)**

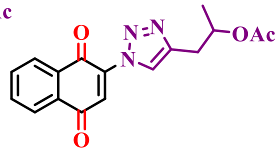

**(281)**

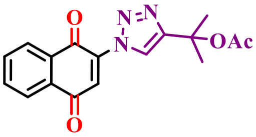

**(282)**

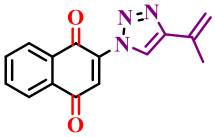

**(283)**

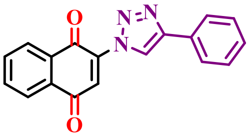

**(284)**

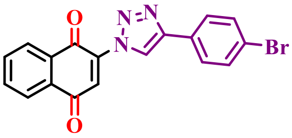

**(285)**

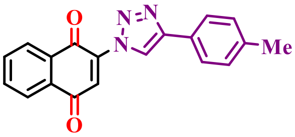

**(286)**

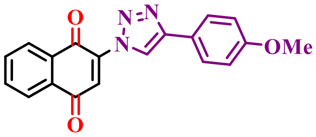

**(287)**

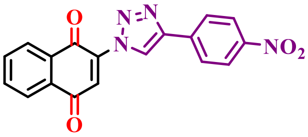

**(288)**

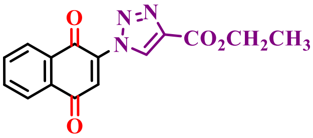

**(289)**

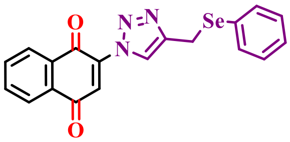

**(290)**

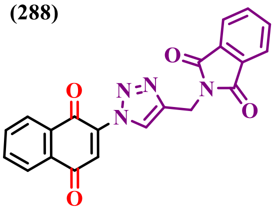

**(291)**

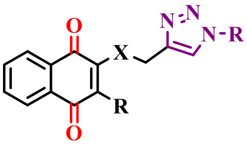

X= N, 30 compounds
X= O, 11 compounds

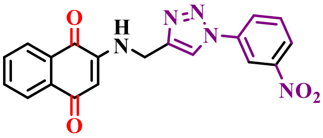

**(292)**

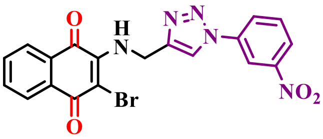

**(293)**

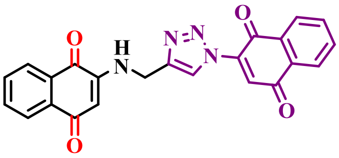

**(294)**

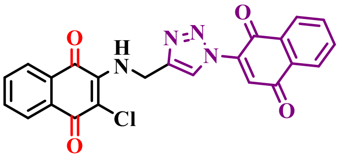

**(295)**

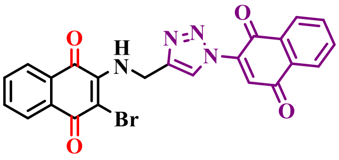

**(296)**

**Figure S18.** *Para*-quinones-based 1,2,3-triazoles (part II).

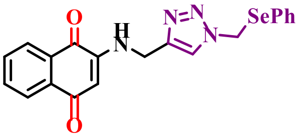

**(297)**

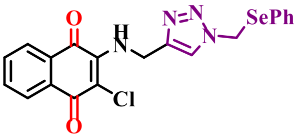

**(298)**

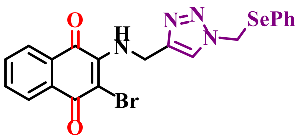

**(299)**

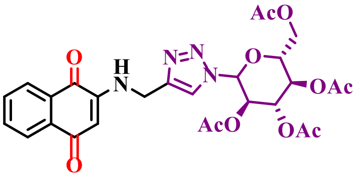

**(300)**

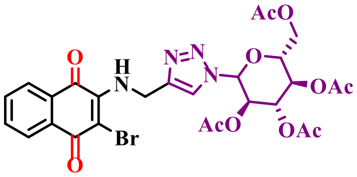

**(301)**

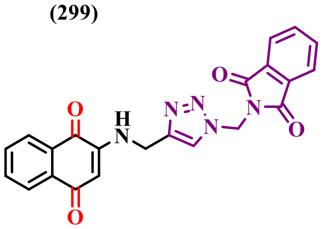

**(302)**

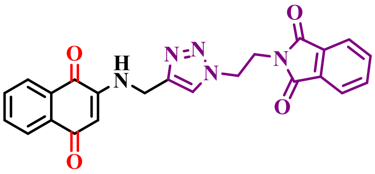

**(303)**

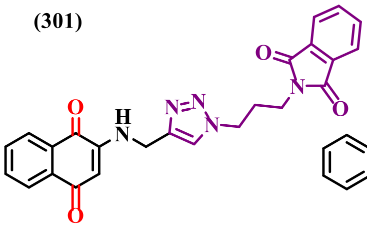

**(304)**

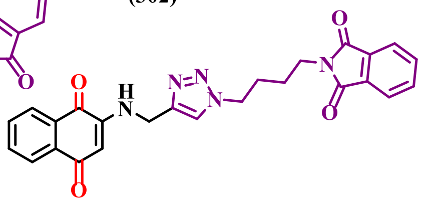

**(305)**

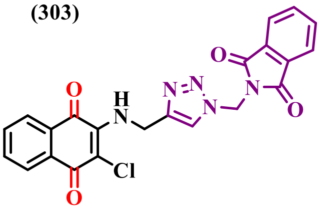

**(306)**

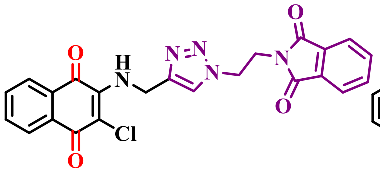

**(307)**

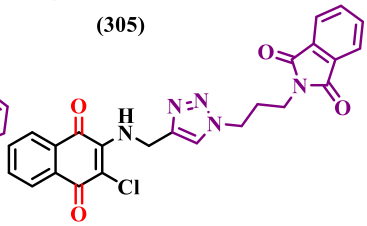

**(308)**

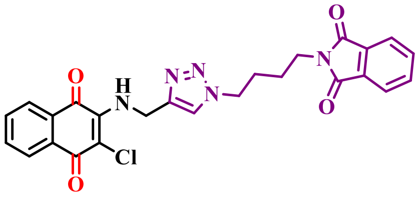

**(309)**

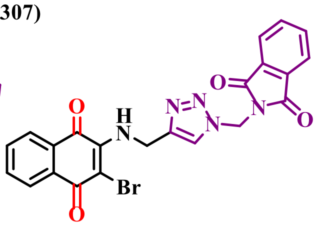

**(310)**

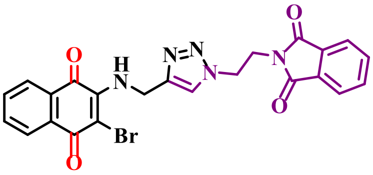

**(311)**

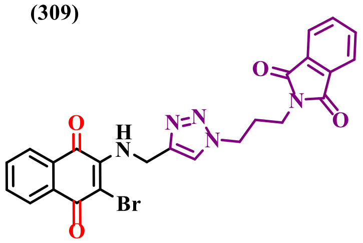

**(312)**

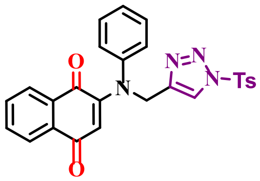

**(313)**

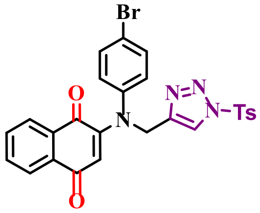

**(314)**

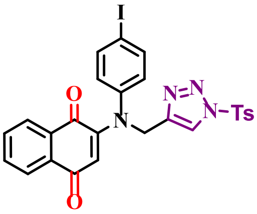

**(315)**

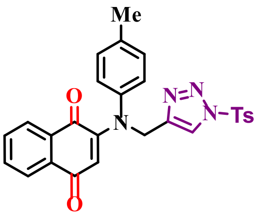

**(316)**

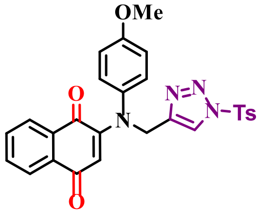

**(317)**

**Figure S19.** Para-quinones-based 1,2,3-triazoles (part III).

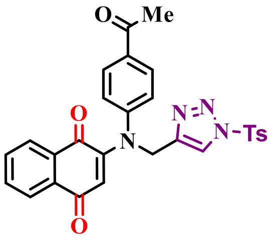

(318)

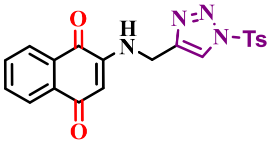

(319)

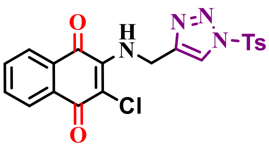

(320)

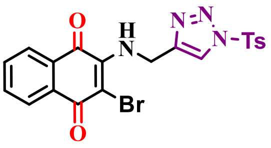

(321)

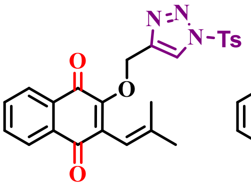

(322)

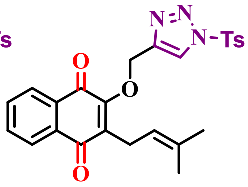

(323)

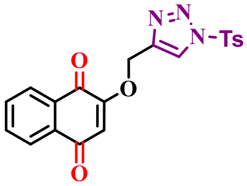

(324)

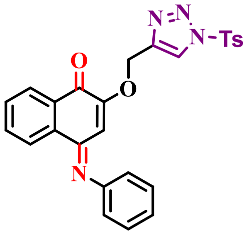

(325)

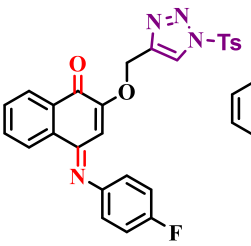

(326)

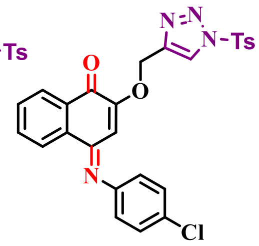

(327)

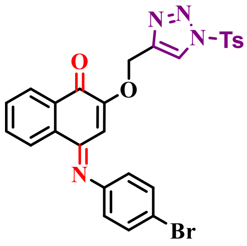

(328)

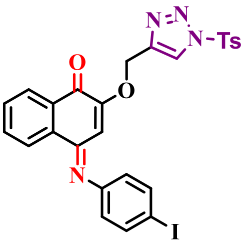

(329)

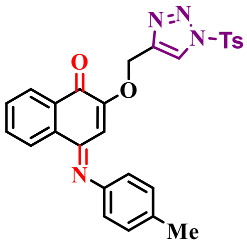

(330)

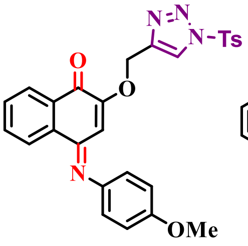

(331)

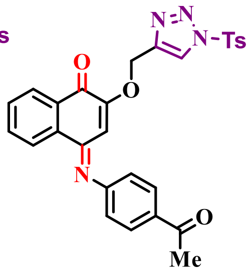

(332)

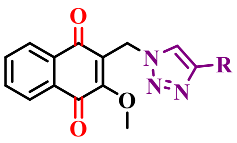

12 compounds

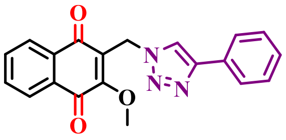

(333)

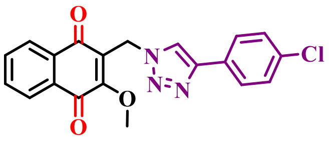

(334)

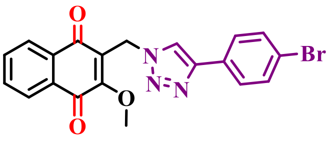

(335)

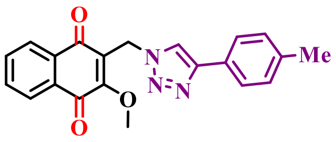

(336)

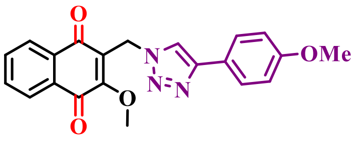

(337)

**Figure S20.** Para-quinones-based 1,2,3-triazoles (part IV).

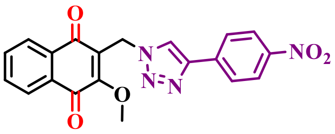

**(338)**

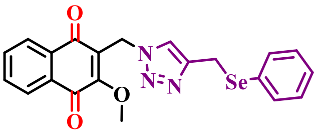

**(339)**

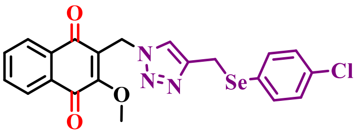

**(340)**

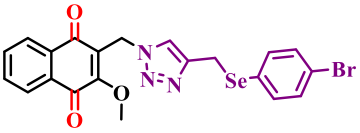

**(341)**

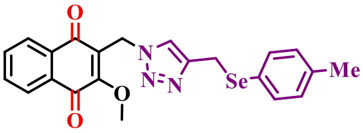

**(342)**

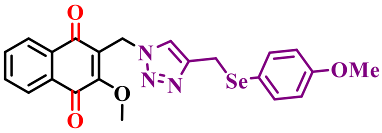

**(343)**

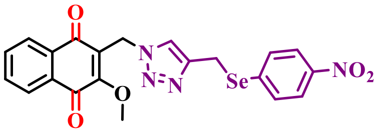

**(344)**

---

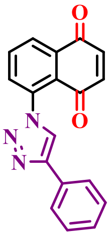

**(345)**

**Figure S21.** Para-quinones-based 1,2,3-triazoles (part V).

#### 1.6 Group 5: Phenazines derivatives

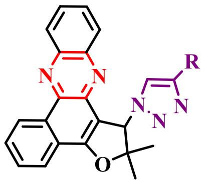

12 compounds

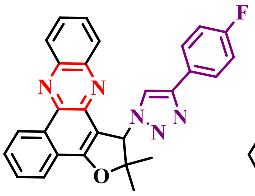

(346)

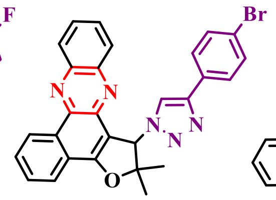

(347)

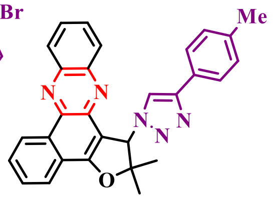

(348)

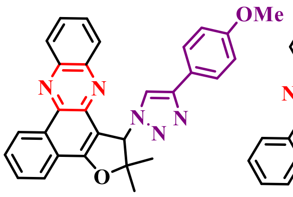

(349)

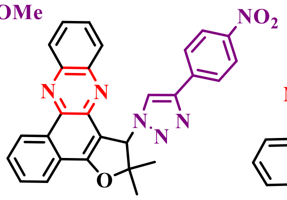

(350)

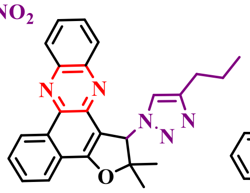

(351)

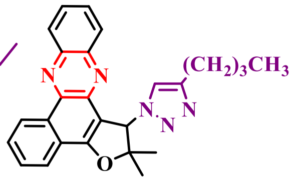

(352)

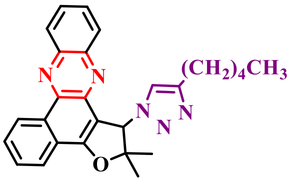

(353)

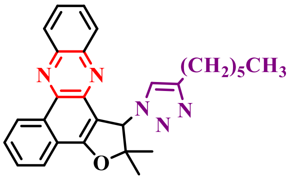

(354)

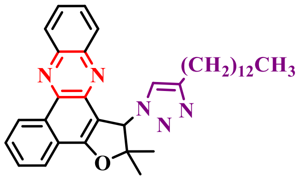

(355)

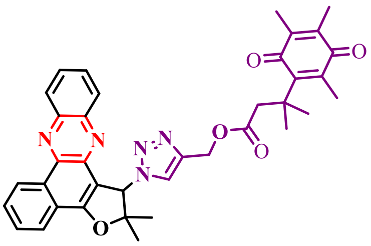

(356)

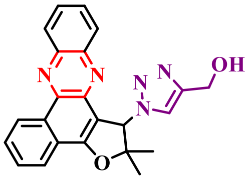

(357)

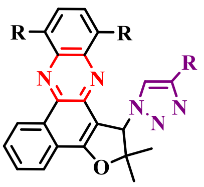

9 compounds

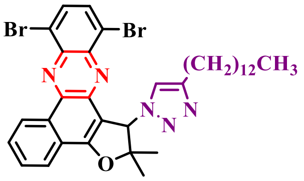

(358)

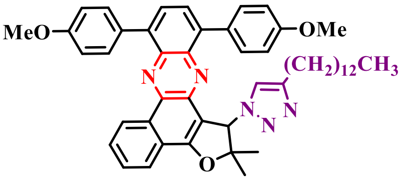

(359)

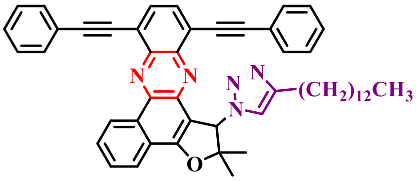

(360)

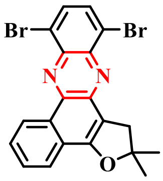

(361)

**Figure S22.** Group 5: Phenazines derivatives (part I).

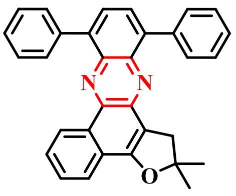

(362)

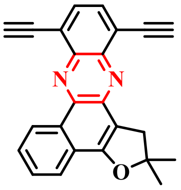

(363)

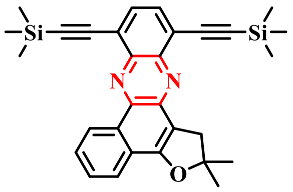

(364)

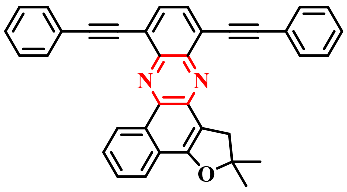

(365)

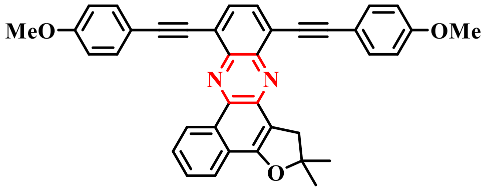

(366)

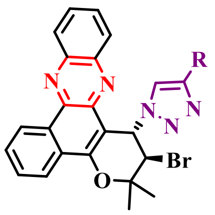

5 compounds

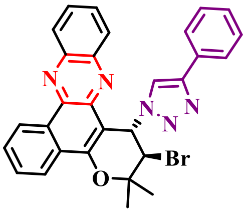

(367)

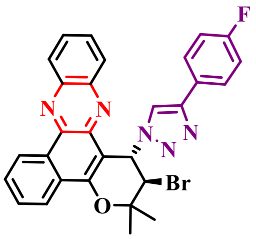

(368)

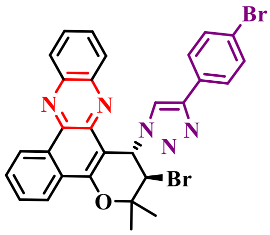

(369)

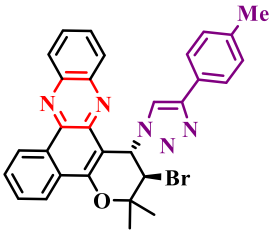

(370)

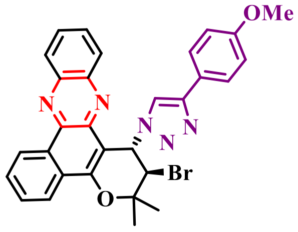

(371)

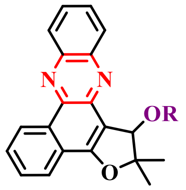

2 compounds

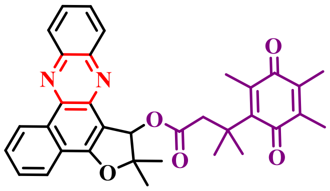

(372)

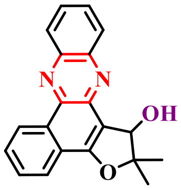

(373)

**Figure S23.** Group 5: Phenazines derivatives (part II).

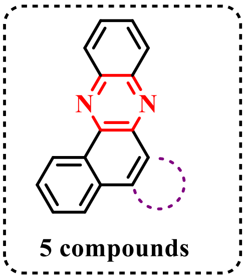

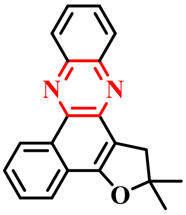

**(374)**

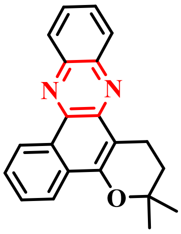

**(375)**

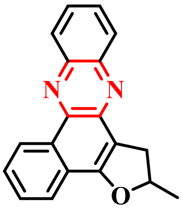

**(376)**

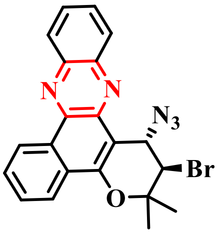

**(377)**

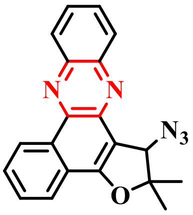

**(378)**

**Figure S24.** Group 5: Phenazines derivatives (part III).

#### 1.7 Group 6: 1,4-naphtoquinones and derivatives

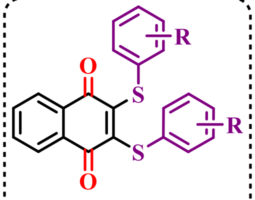

6 compounds

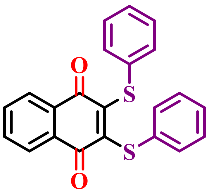

(379)

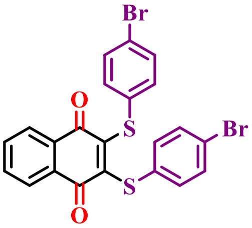

(380)

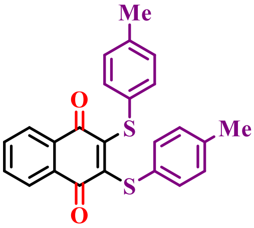

(381)

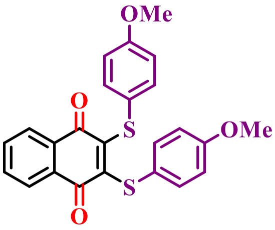

(382)

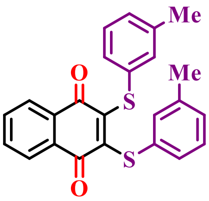

(383)

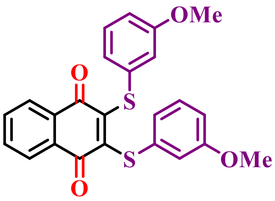

(384)

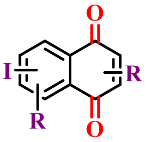

21 compounds

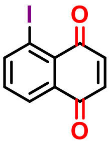

(385)

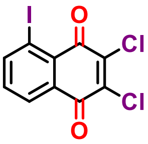

(386)

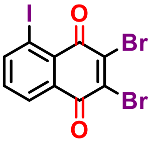

(387)

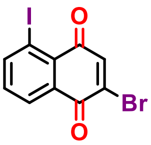

(388)

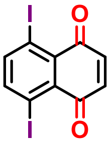

(389)

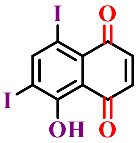

(390)

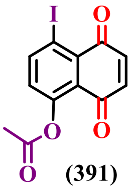

(391)

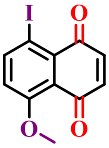

(392)

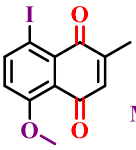

(393)

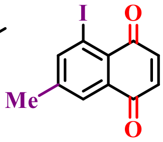

(394)

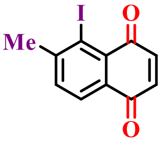

(395)

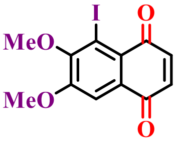

(396)

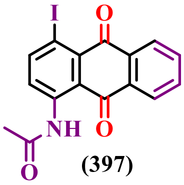

(397)

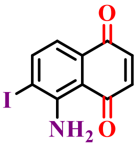

(398)

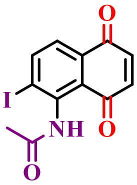

(399)

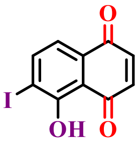

(400)

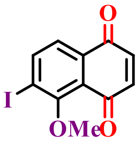

(401)

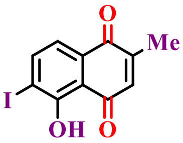

(402)

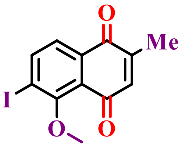

(403)

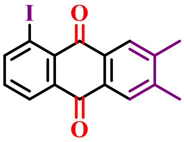

(404)

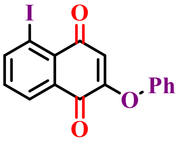

(405)

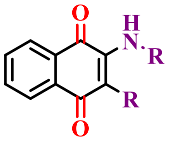

23 compounds

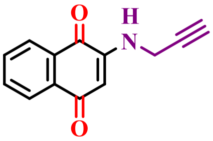

(406)

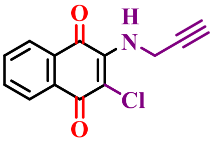

(407)

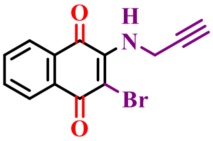

(408)

**Figure S25.** Group 6: 1,4-naphtoquinones and derivatives (part I).

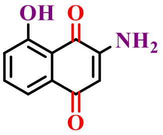

**(409)**

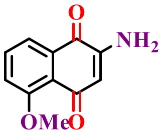

**(410)**

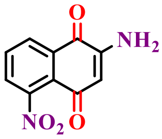

**(411)**

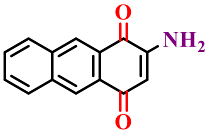

**(412)**

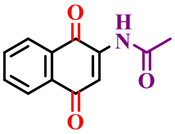

**(413)**

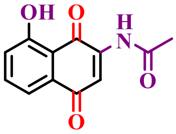

**(414)**

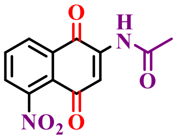

**(415)**

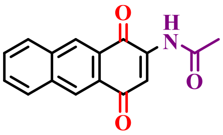

**(416)**

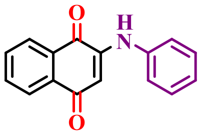

**(417)**

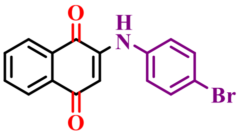

**(418)**

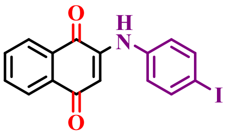

**(419)**

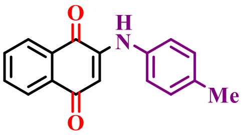

**(420)**

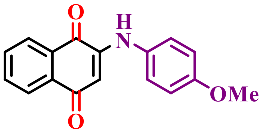

**(421)**

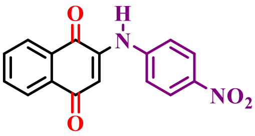

**(422)**

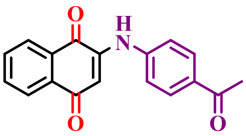

**(423)**

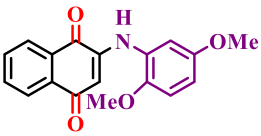

**(424)**

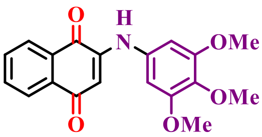

**(425)**

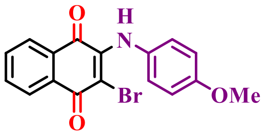

**(426)**

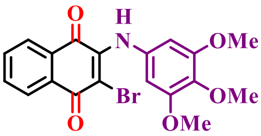

**(427)**

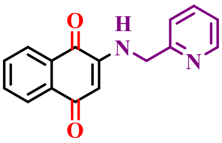

**(428)**

---

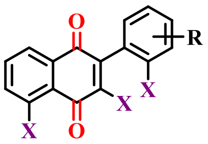

**X = H or OH**
**29 compounds**

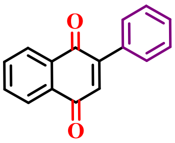

**(429)**

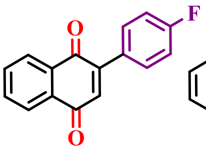

**(430)**

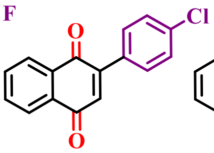

**(431)**

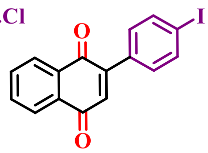

**(432)**

**Figure S26.** Group 6: 1,4-naphtoquinones and derivatives (part II).

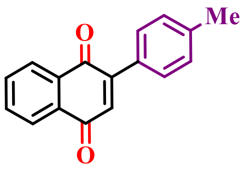

**(433)**

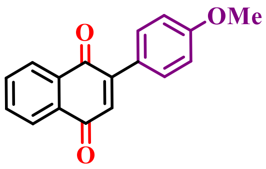

**(434)**

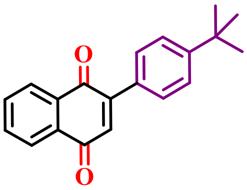

**(435)**

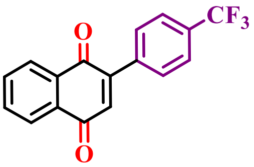

**(436)**

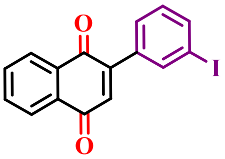

**(437)**

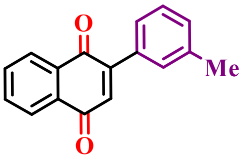

**(438)**

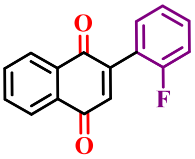

**(439)**

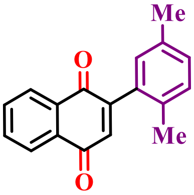

**(440)**

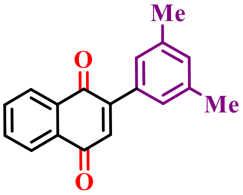

**(441)**

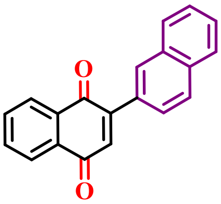

**(442)**

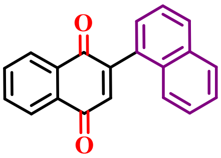

**(443)**

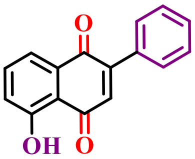

**(444)**

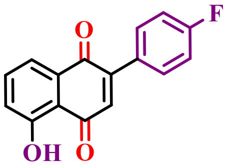

**(445)**

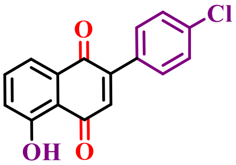

**(446)**

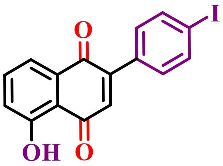

**(447)**

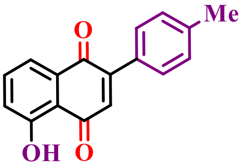

**(448)**

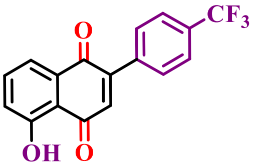

**(449)**

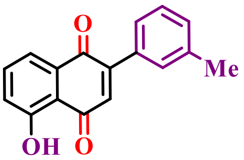

**(450)**

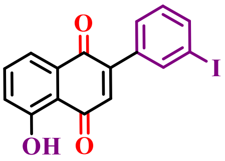

**(451)**

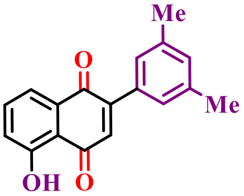

**(452)**

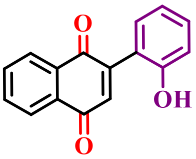

**(453)**

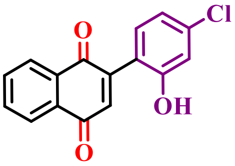

**(454)**

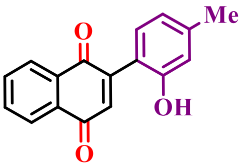

**(455)**

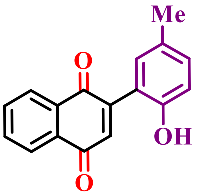

**(456)**

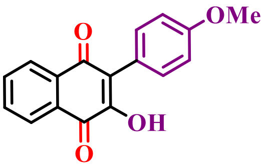

**(457)**

**Figure S27.** Group 6: 1,4-naphtoquinones and derivatives (part III).

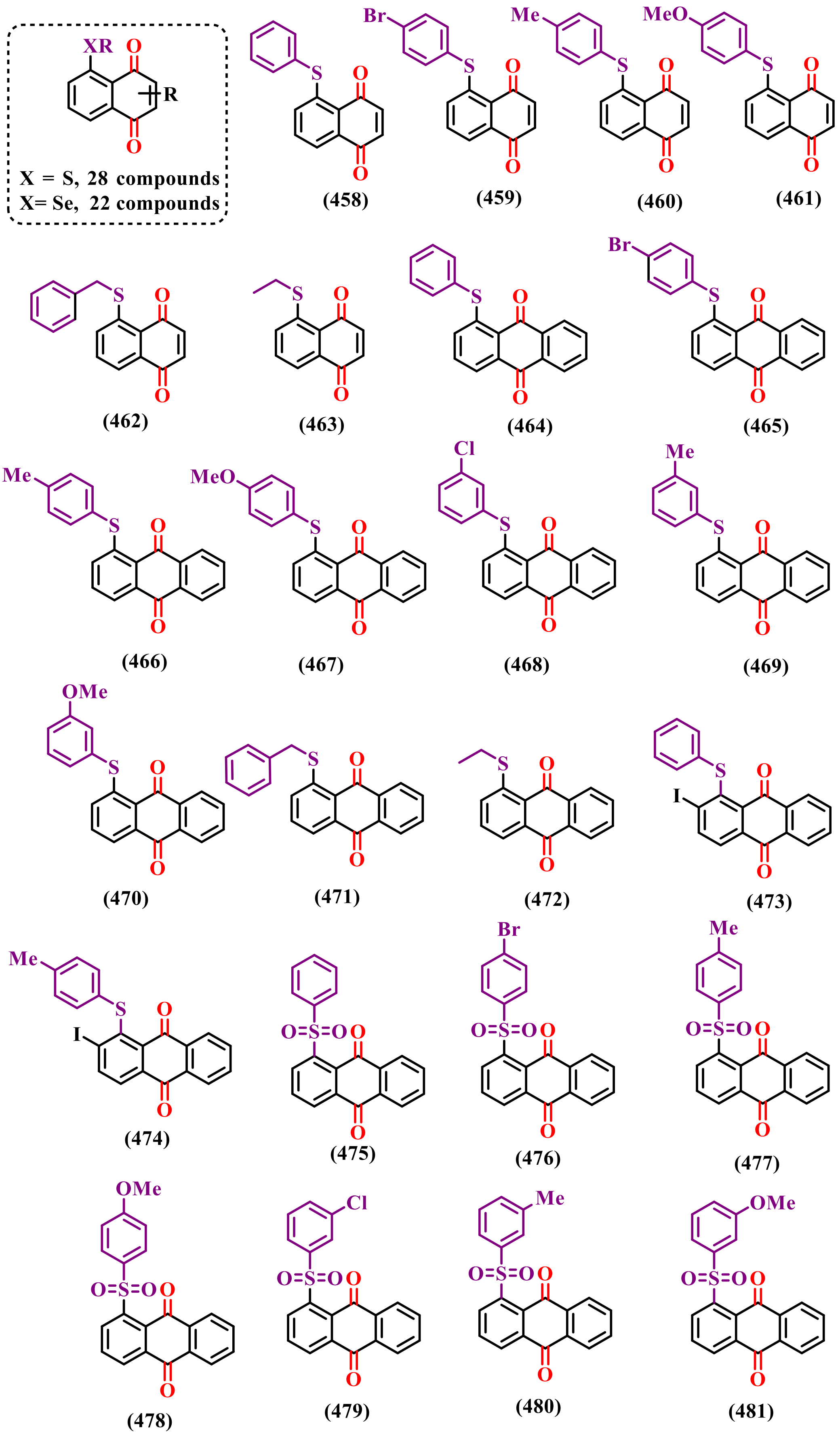

**Figure S28.** Group 6: 1,4-naphtoquinones and derivatives (part IV).

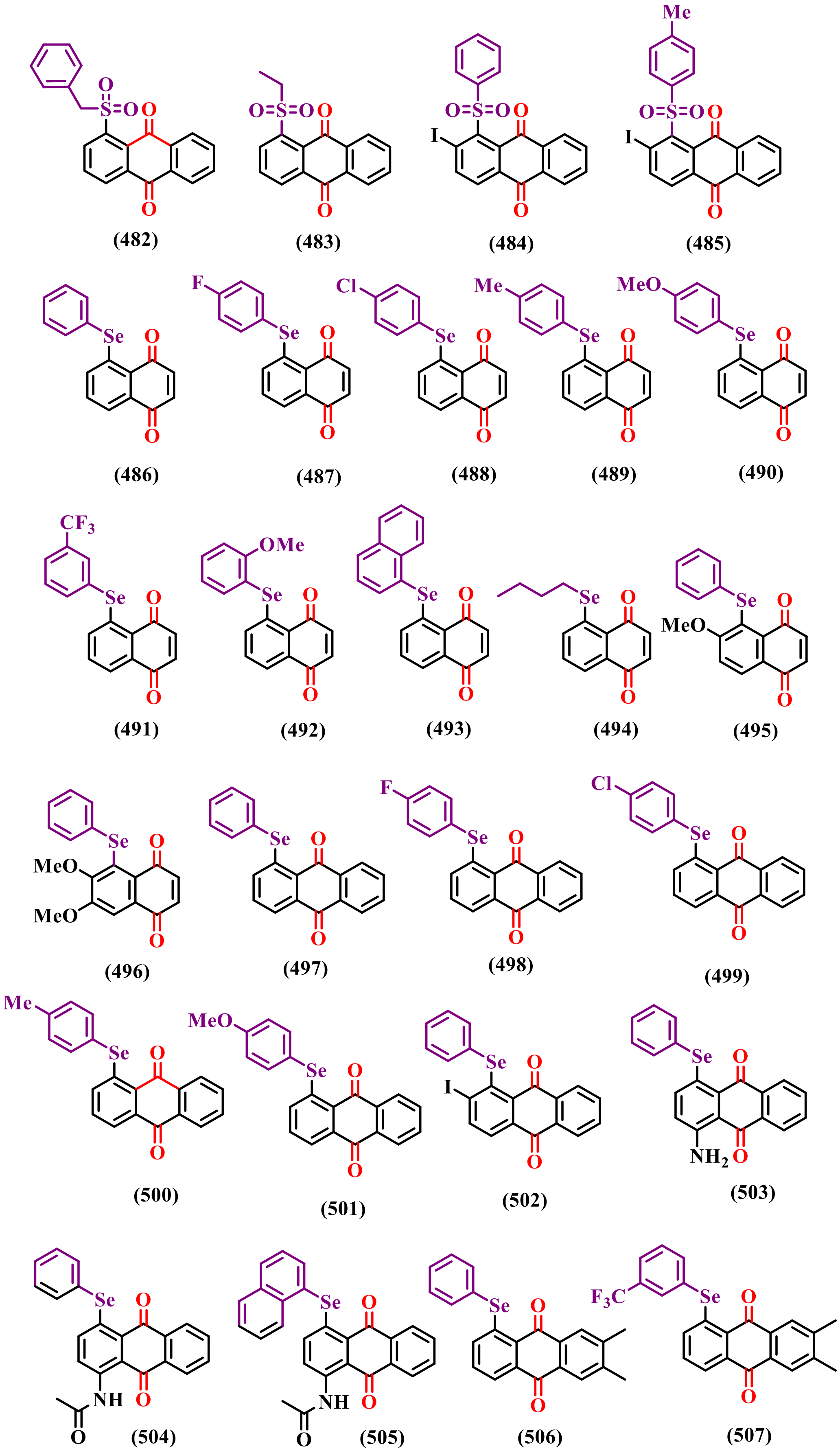

**Figure S29.** Group 6: 1,4-naphtoquinones and derivatives (part V).

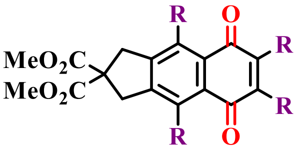

19 compounds

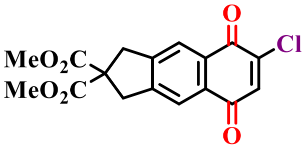

(508)

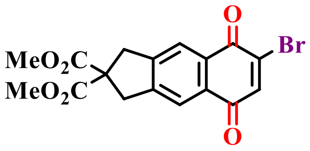

(509)

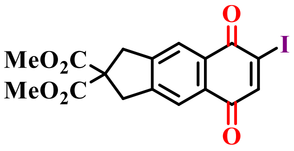

(510)

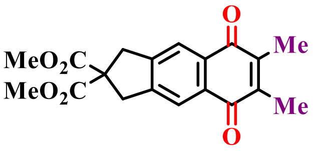

(511)

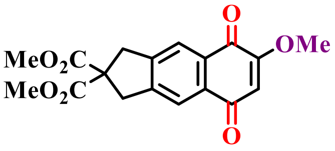

(512)

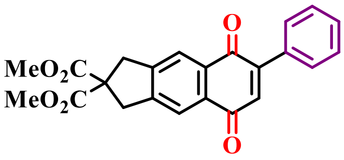

(513)

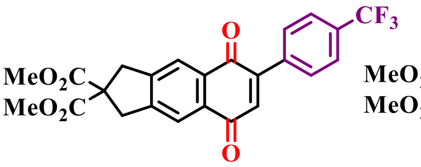

(514)

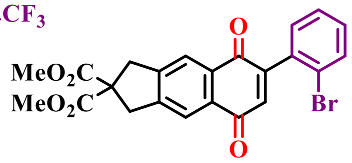

(515)

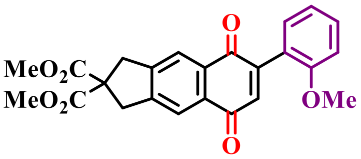

(516)

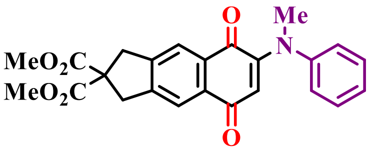

(517)

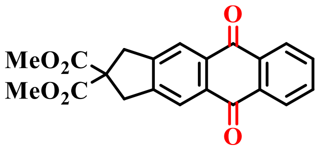

(518)

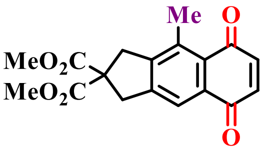

(519)

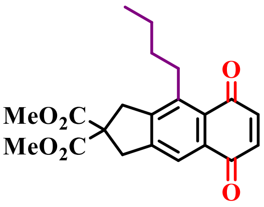

(520)

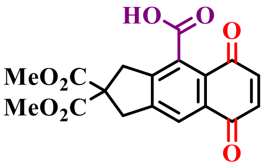

(521)

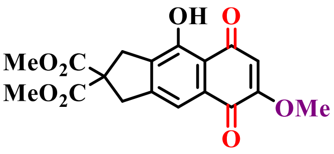

(522)

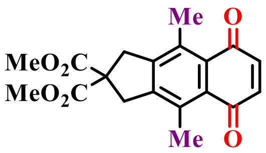

(523)

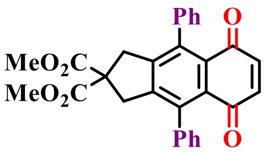

(524)

**Figure S30.** Group 6: 1,4-naphtoquinones and derivatives (part VI).

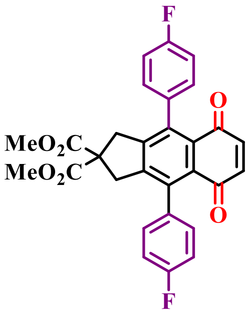

(525)

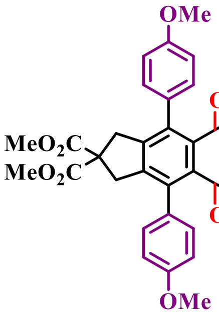

(526)

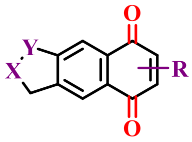

6 compounds

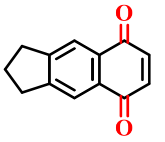

(527)

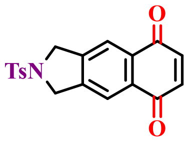

(528)

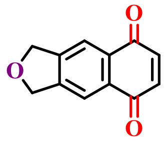

(529)

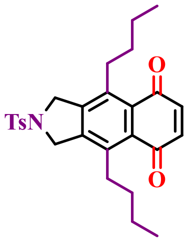

(530)

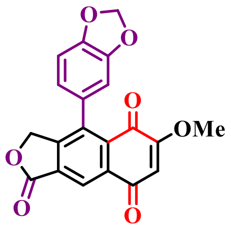

(531)

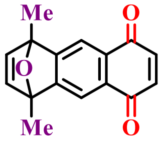

(532)

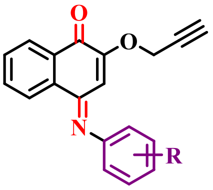

8 compounds

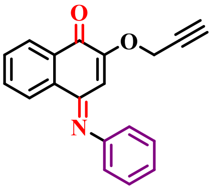

(533)

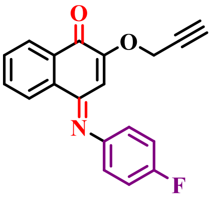

(534)

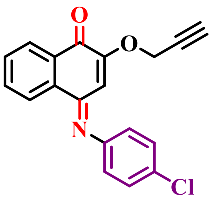

(535)

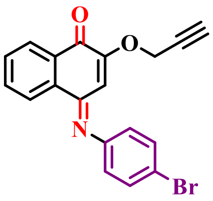

(536)

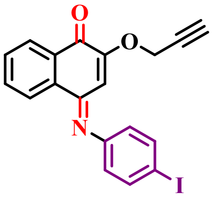

(537)

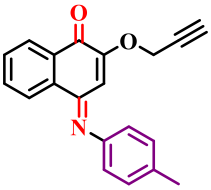

(538)

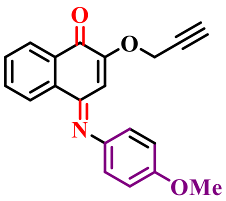

(539)

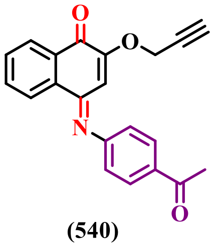

(540)

**Figure S31.** Group 6: 1,4-naphtoquinones and derivatives (part VII).

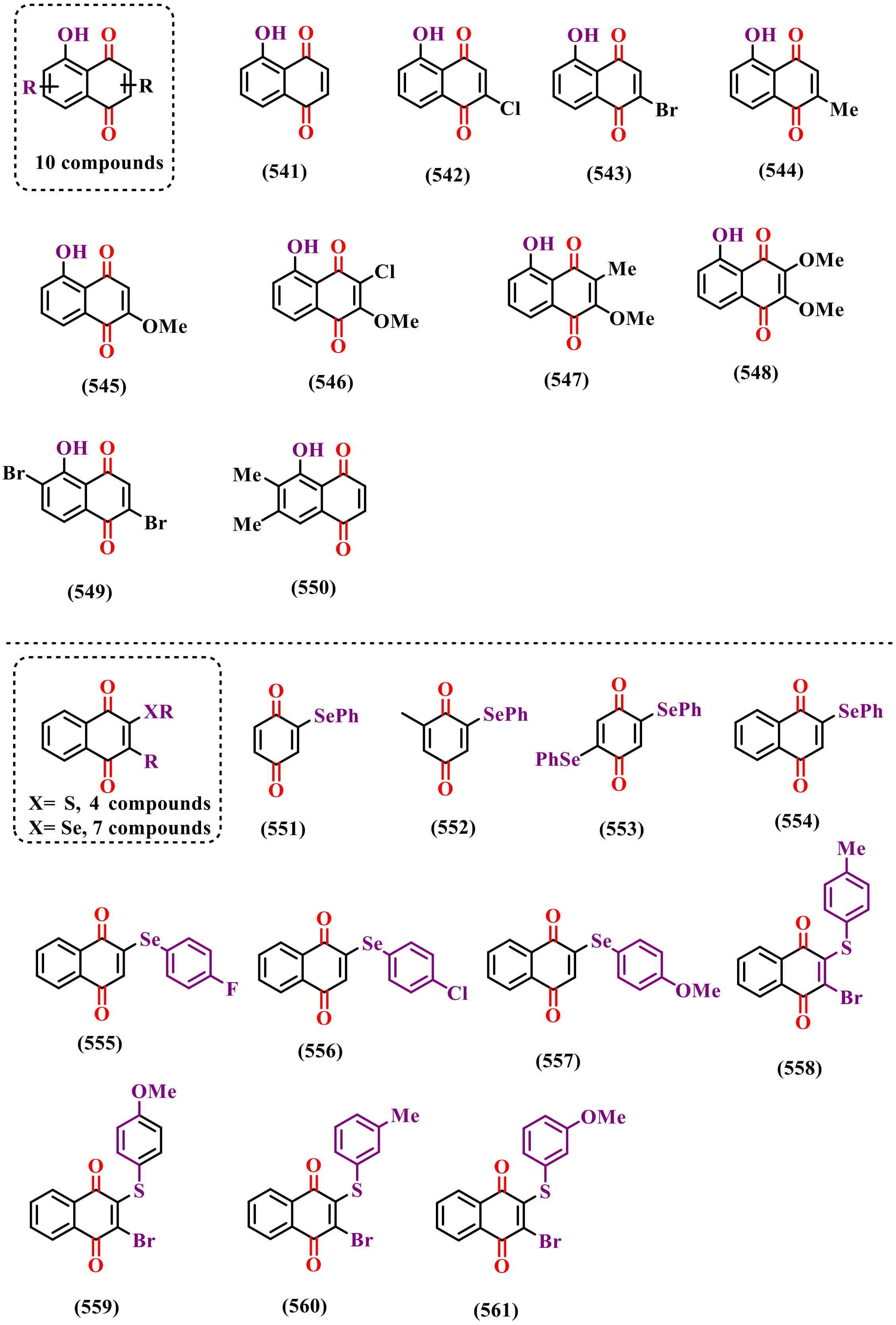

**Figure S32.** Group 6: 1,4-naphtoquinones and derivatives (part VIII).

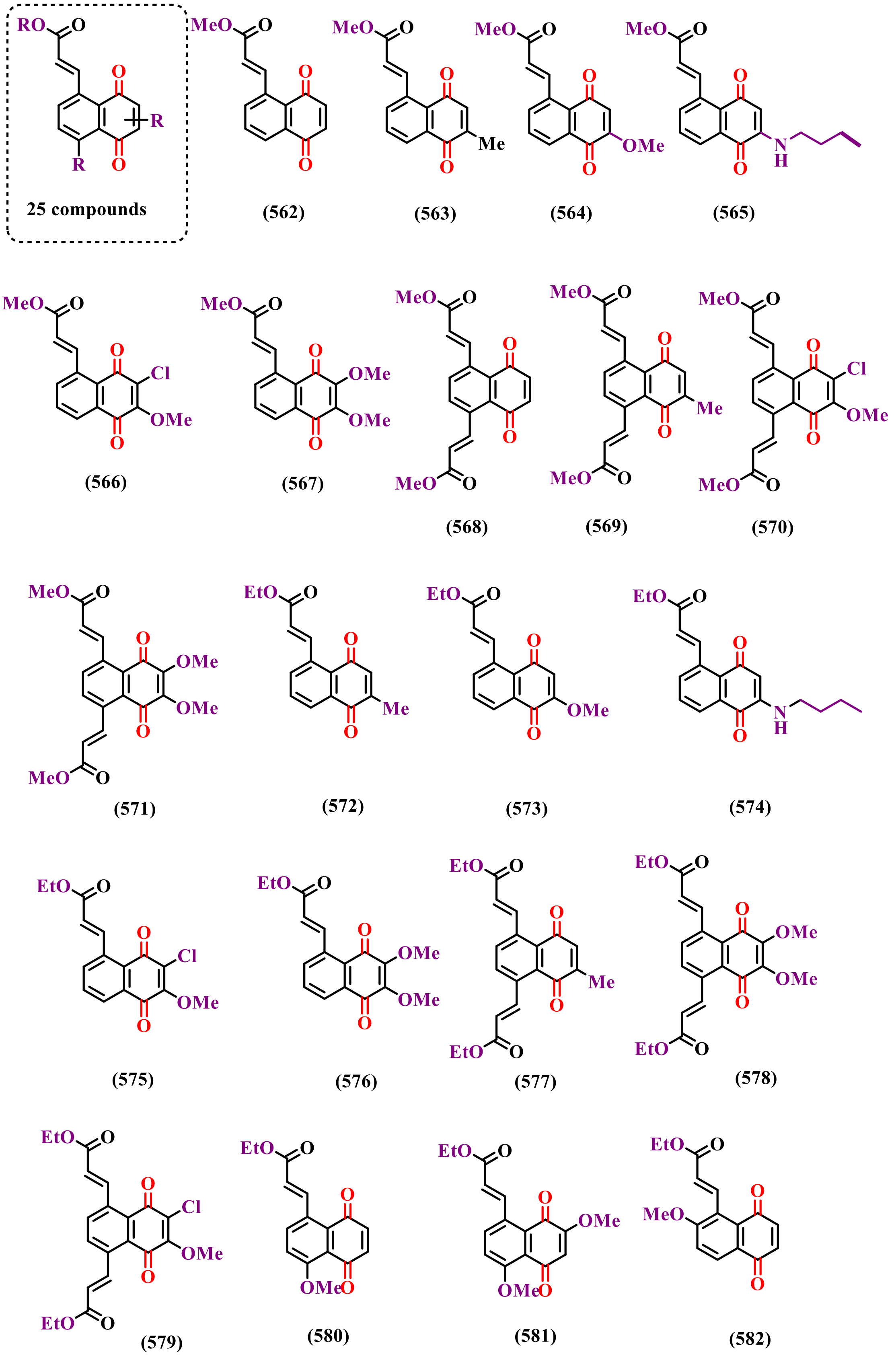

**Figure S33.** Group 6: 1,4-naphtoquinones and derivatives (part IX).

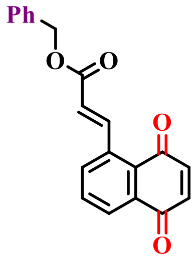
(583)

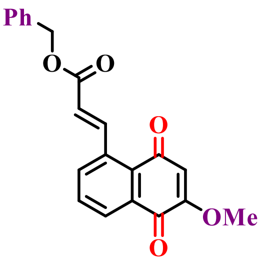
(584)

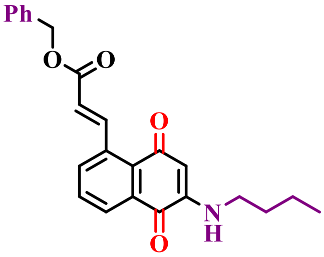
(585)

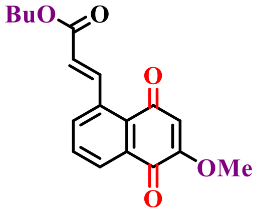
(586)

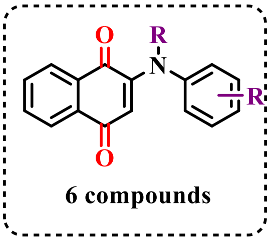

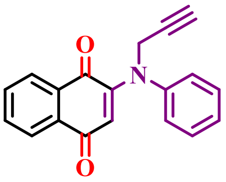
(587)

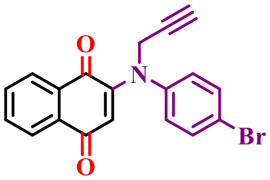
(588)

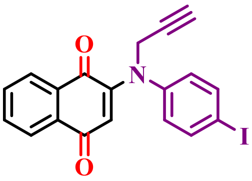
(589)

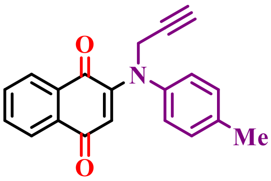
(590)

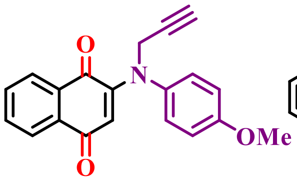
(591)

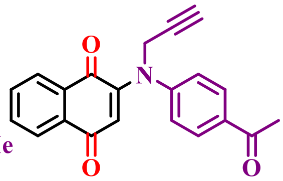
(592)

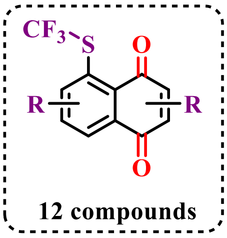

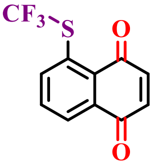
(593)

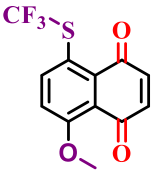
(594)

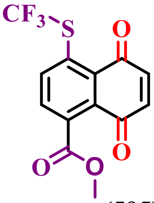
(595)

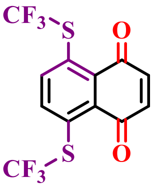
(596)

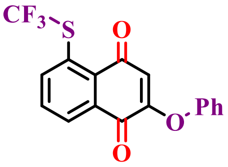
(597)

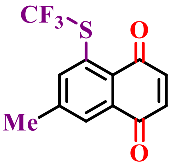
(598)

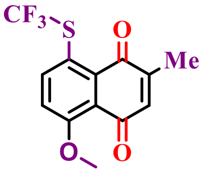
(599)

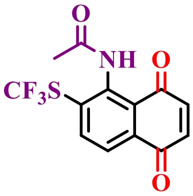
(600)

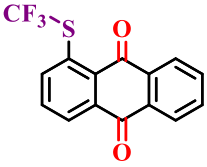
(601)

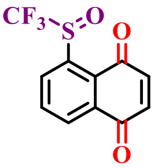
(602)

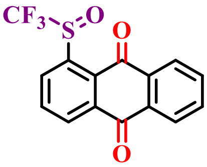
(603)

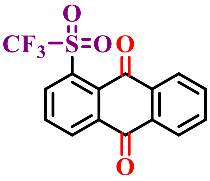
(604)

**Figure S34.** Group 6: 1,4-naphtoquinones and derivatives (part X).

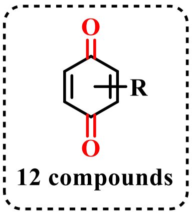

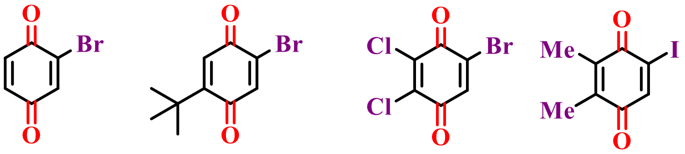

(605) (606) (607) (608)

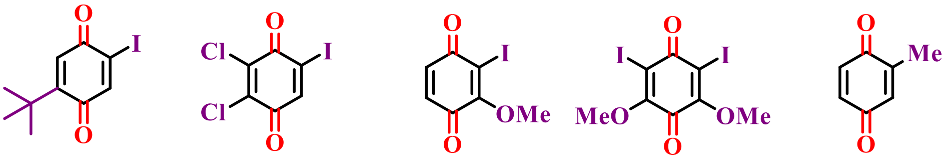

(609) (610) (611) (612) (613)

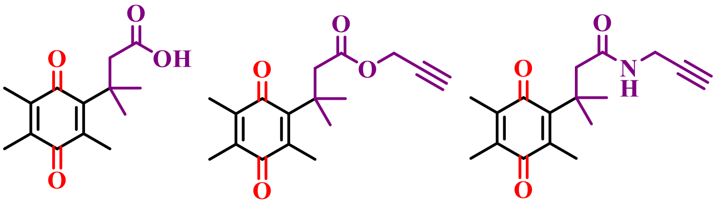

(614) (615) (616)

---

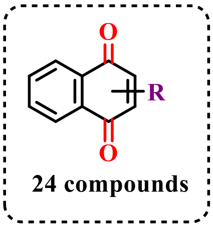

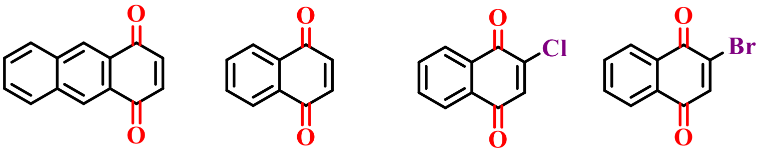

(617) (618) (619) (620)

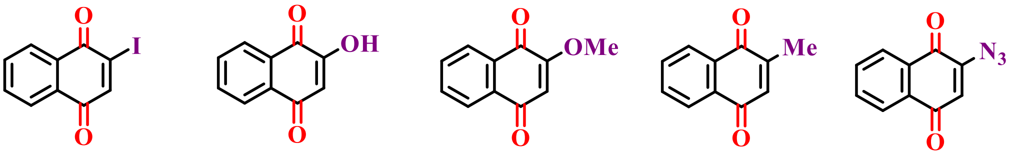

(621) (622) (623) (624) (625)

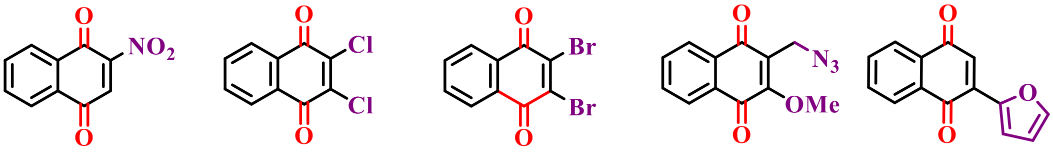

(626) (627) (628) (629) (630)

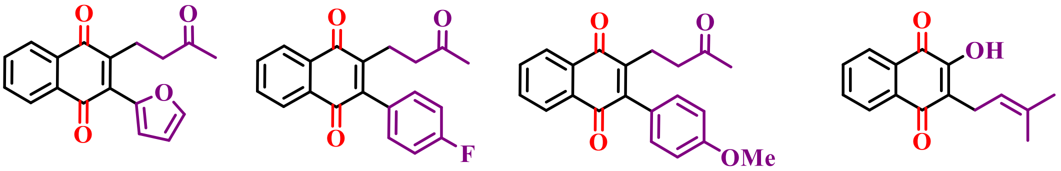

(631) (632) (633) (634)

**Figure S35.** Group 6: 1,4-naphtoquinones and derivatives (part XI).

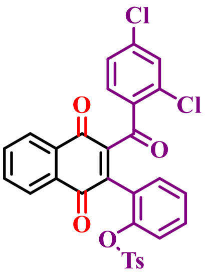

(635)

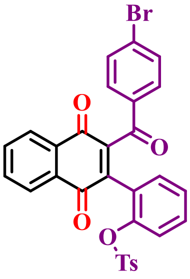

(636)

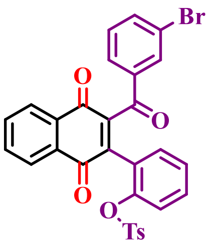

(637)

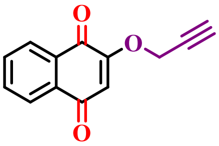

(638)

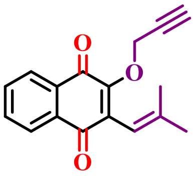

(639)

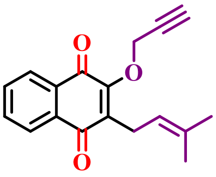

(640)

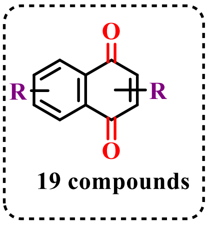

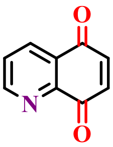

(641)

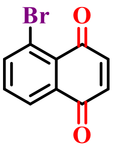

(642)

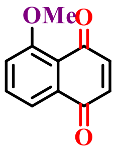

(643)

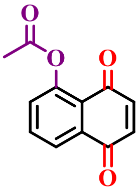

(644)

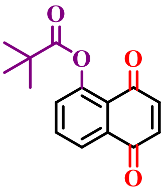

(645)

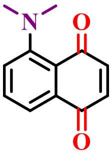

(646)

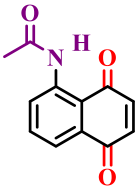

(647)

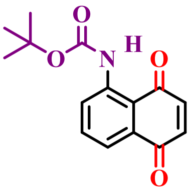

(648)

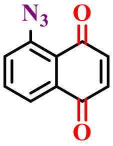

(649)

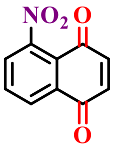

(650)

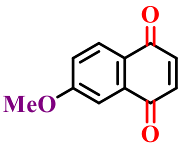

(651)

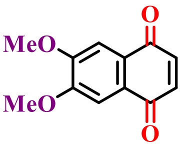

(652)

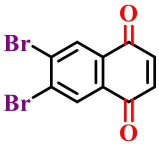

(653)

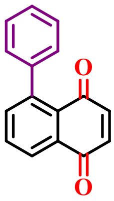

(654)

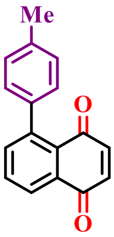

(655)

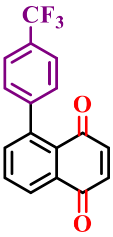

(656)

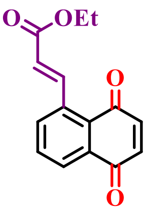

(657)

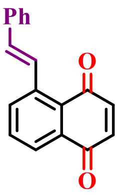

(658)

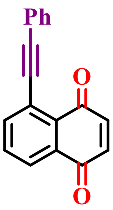

(659)

**Figure S36.** Group 6: 1,4-naphtoquinones and derivatives (part XII).

#### 1.8 Grupo 7: Hydrazo derivatives

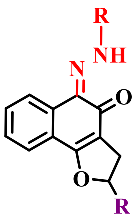

2 compounds

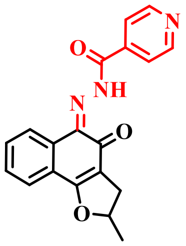

(660)

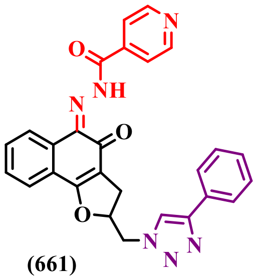

(661)

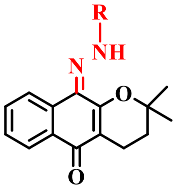

2 compounds

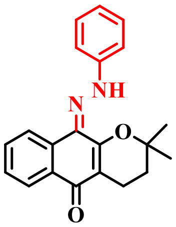

(662)

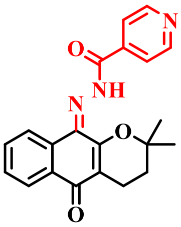

(663)

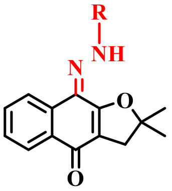

2 compound

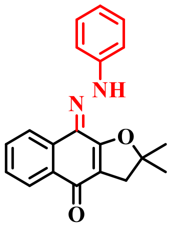

(664)

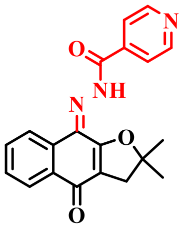

(665)

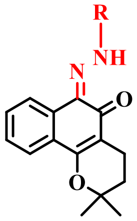

6 compounds

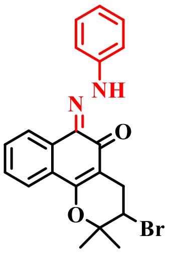

(666)

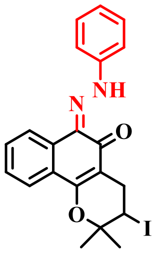

(667)

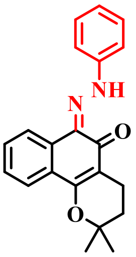

(668)

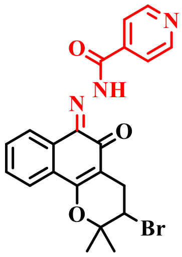

(669)

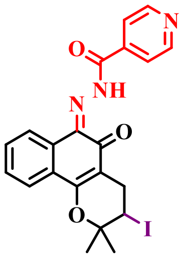

(670)

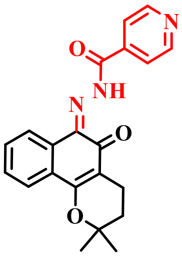

(671)

**Figure S37.** Grupo 7: Hydrazo derivatives (part I).

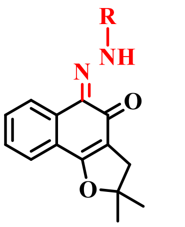

5 compounds

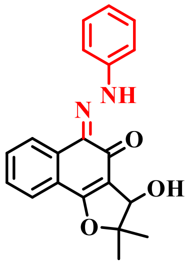

(672)

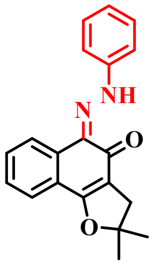

(673)

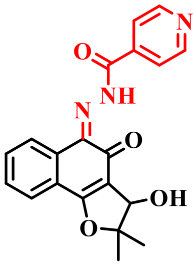

(674)

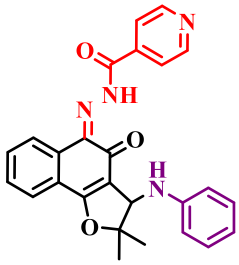

(675)

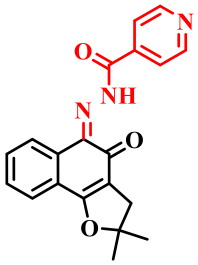

(676)

**Figure S38.** Grupo 7: Hydrazo derivatives (part II).

#### 1.9 Grupo 8: Imidazole and oxazole derivatives derivatives

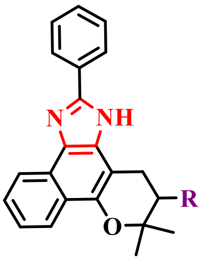

1 compound

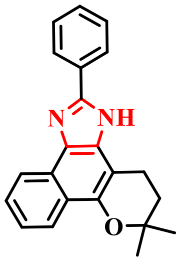

(677)

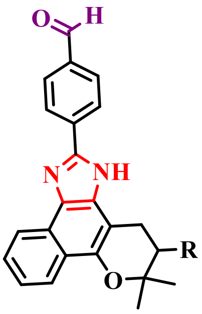

1 compound

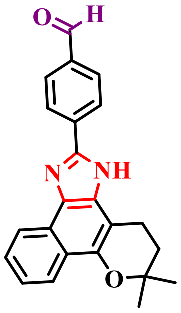

(678)

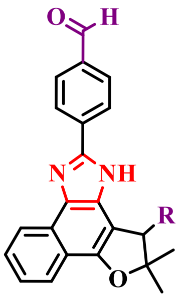

1 compound

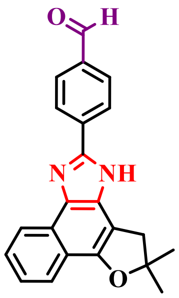

(679)

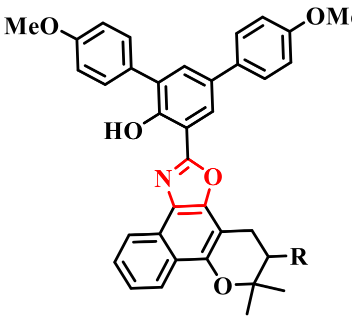

1 compound

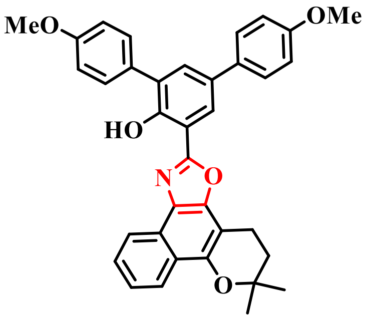

(680)

**Figure S39.** Grupo 8: Imidazole and oxazole derivatives derivatives (part I).

.

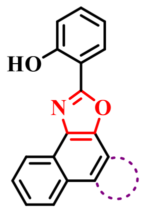

7 compounds

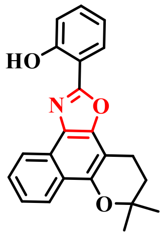

(681)

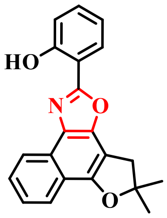

(682)

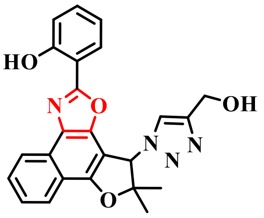

(683)

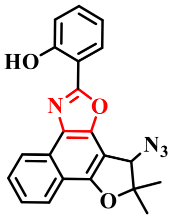

(684)

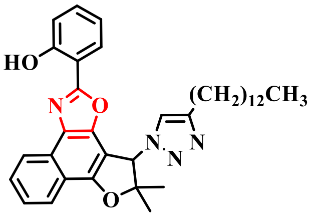

(685)

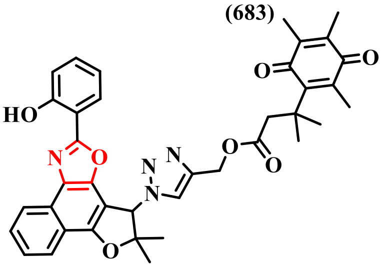

(686)

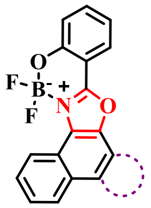

4 compounds

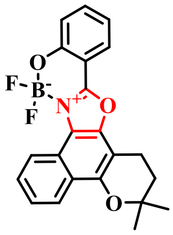

(687)

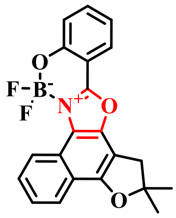

(688)

**Figure S40.** Grupo 8: Imidazole and oxazole derivatives derivatives (part II).

#### 2. Computational Approaches

##### 2.1 Available Mpro structures show conserved conformation, protein-ligand interactions, and location of waters molecules

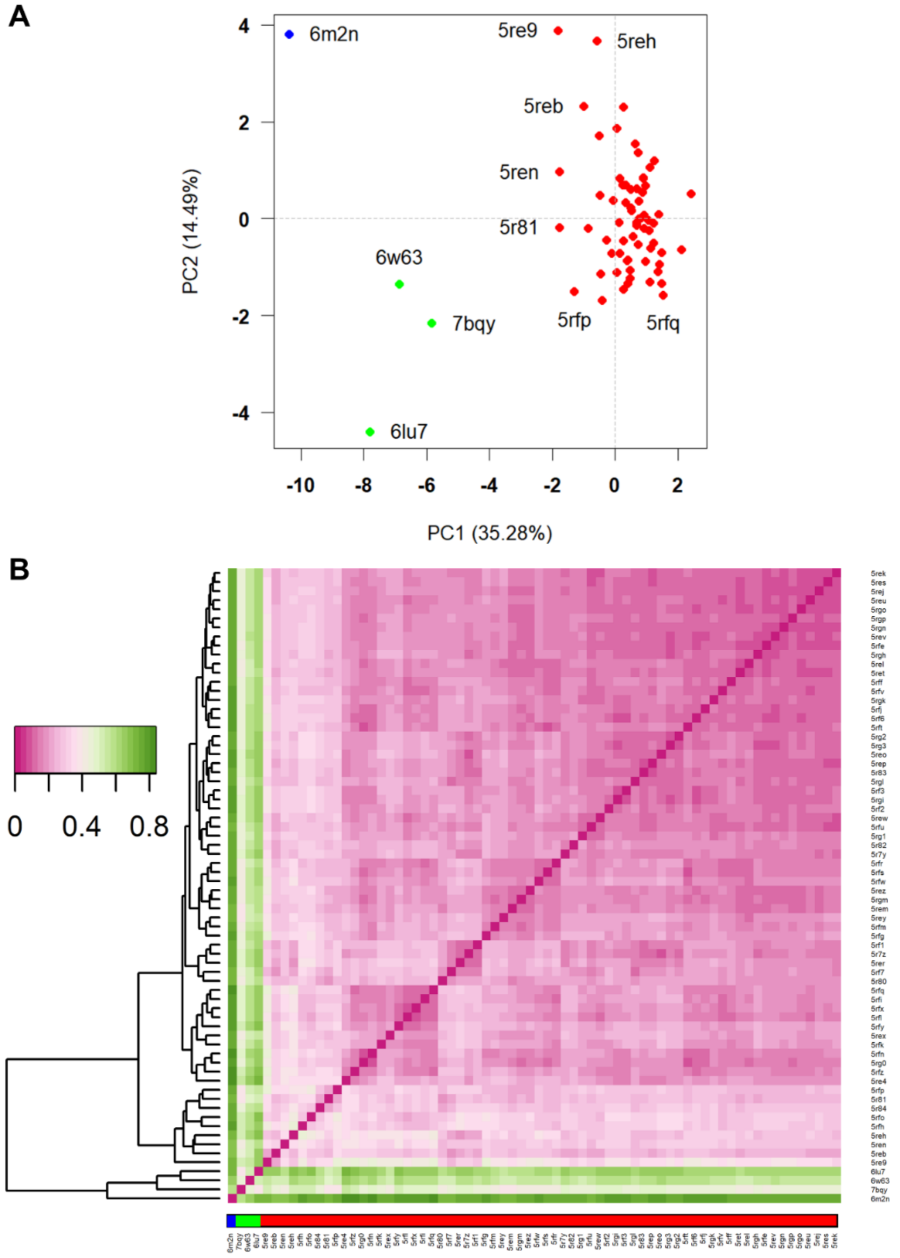

**Figure S41.** Structural analysis of 72 available SARS-CoV-2 Mpro structures in the PDB. A) Principal component analysis (PCA) represented by the first two components, PC1 and PC2, showed close to 50% structural variance using program R's package Bio3D. The PC1 versus PC2 plot displayed three structural clusters. One of the clusters (red dots) were the most populated showing the high similarity among most of Mpro structures evaluated. Two other clusters could be found (blue and green) and show structures that distances from the most populated cluster. B) Heatmaps and dendrograms representation of carbon alfa (Cα) root-mean-square deviation (RMSD) results between the structures. RMSD ranged from 0 (dark pink) to under 1.0 Å (dark green), with only the structures gather in the blue and green clusters showing a slightly elevated value (bottom rectangle), thus, conformation changes might be in specific regions for the different clusters.

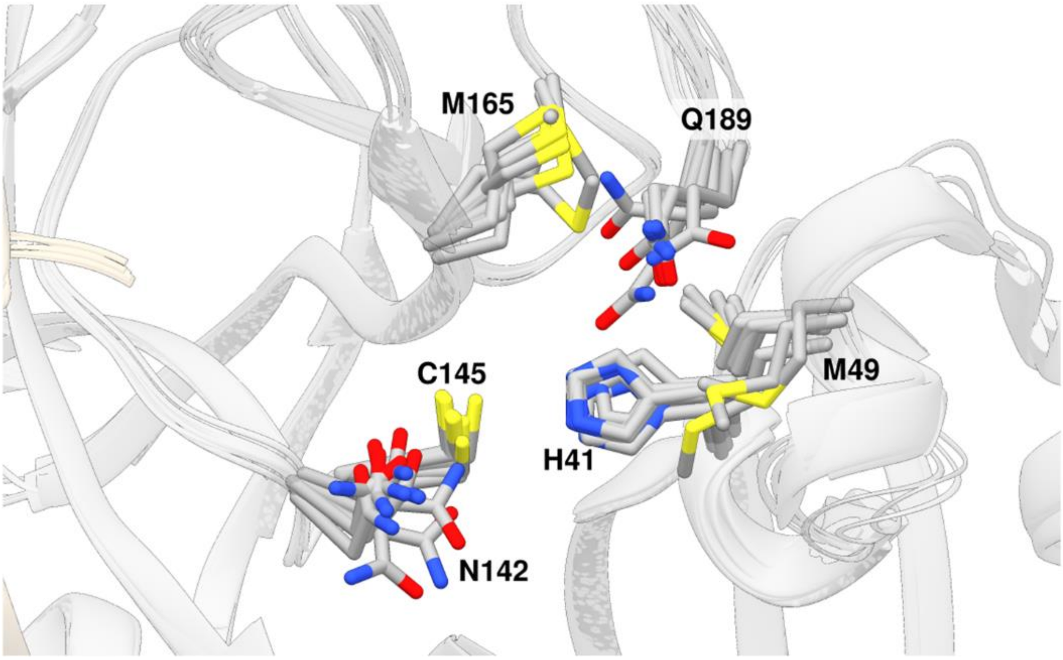

**Figure S42.** Superposition of ten Mpro PDB structures (PDB Codes: 5R82, 5RFW, 5RF6, 5RFE, 5RFV, 5RF3, 6M2N, 6W63, 6LU7, and 7BQY). The structures were selected by their high resolution and due to their distance to the other structures in the PCA analysis. The superposition revealed that most residues in the binding site adopt the same conformations, with the residues M49, N142, M165, and Q189 being more flexible among them.

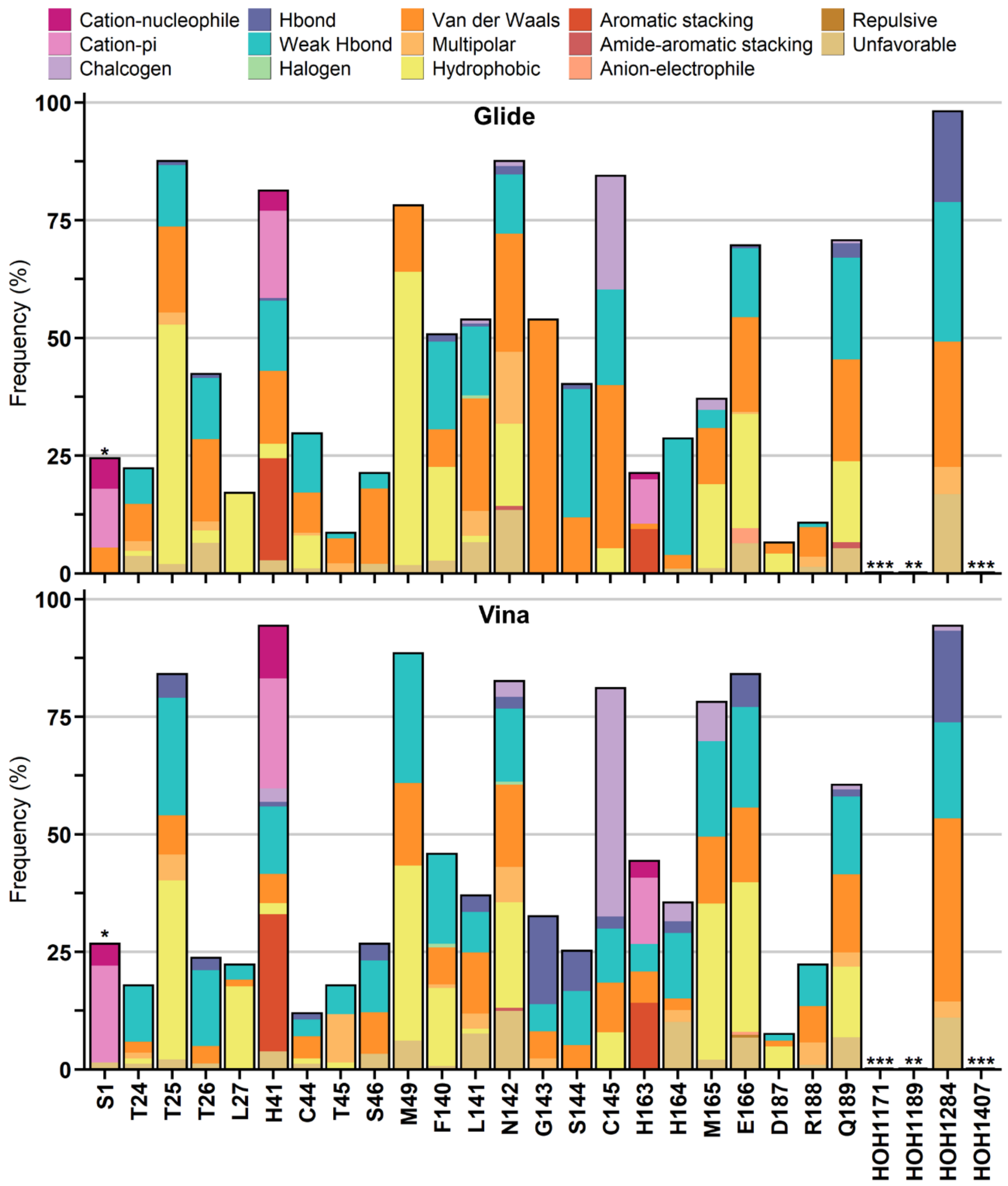

**Figure S43.** Protein-ligand interaction analysis of the selected 70 compounds from docking with Glide e Vina. Residues with (*) are from the other protomer. From docking results, the (**) highlights residues where no interaction was found, while (***) are residues that were not considered in the experiments.

#### 2.2 Virtual screening of naphthoquinoidal compounds against SARS-CoV-2 main protease

**Table S1.** Water molecules conserved in the SARS-CoV-2 main protease described by the ProBiS H2O plugin.

| Probis cluster | Frequency in PDBs | | Water number in PDB 5R82 | Retained for docking ? | Observations |
|---|---|---|---|---|---|
| | Structures | Conservation (%) | | | |
| **0** | 68 | 89 | 1284 | Yes | Could mediate h-bonds to Glu166 and Leu167 |
| **1** | 53 | 70 | Not present? | No | |
| **2** | 48 | 63 | 1189 | Yes | Involved in catalytic triad. Seems inaccessible to interact with ligands. |
| **3** | 41 | 54 | 1407 | No | Seems distant to the ligands |
| **4** | 40 | 53 | 1171 | No | H-bonds to water 1284 (cluster 0) |

**Table S2.** Stability of each compound evaluated by molecular dynamics simulation in each replicate in nanosecond.

| | Mpro | | | | PLpro | | |
|---|---|---|---|---|---|---|---|
| **System** | **415*** | | **382** | | **XR8-89** | **189** | **195** |
| **Rep/Chain** | **A** | **B** | **A** | **B** | **A** | **A** | **A** |
| **1** | 1000 | 0 | 1000 | 1000 | 1000 | 1000 | 1000 |
| **2** | 600 | 400 | 400 | 1000 | 1000 | 1000 | 1000 |
| **3** | 300 | 1000 | 800 | 1000 | 1000 | 1000 | 500 |
| **4** | 0 | 1000 | 1000 | 1000 | 1000 | 1000 | 500 |
| **5** | 850 | 150 | 1000 | 1000 | 1000 | 1000 | 500 |

*data concerning simulations in which compound **415** was not covalently bound to the enzyme. Compound **415** was also simulated covalently bound to Mpro, but the corresponding simulations are not included in the table, since the covalent linkage assures stability of the complex.

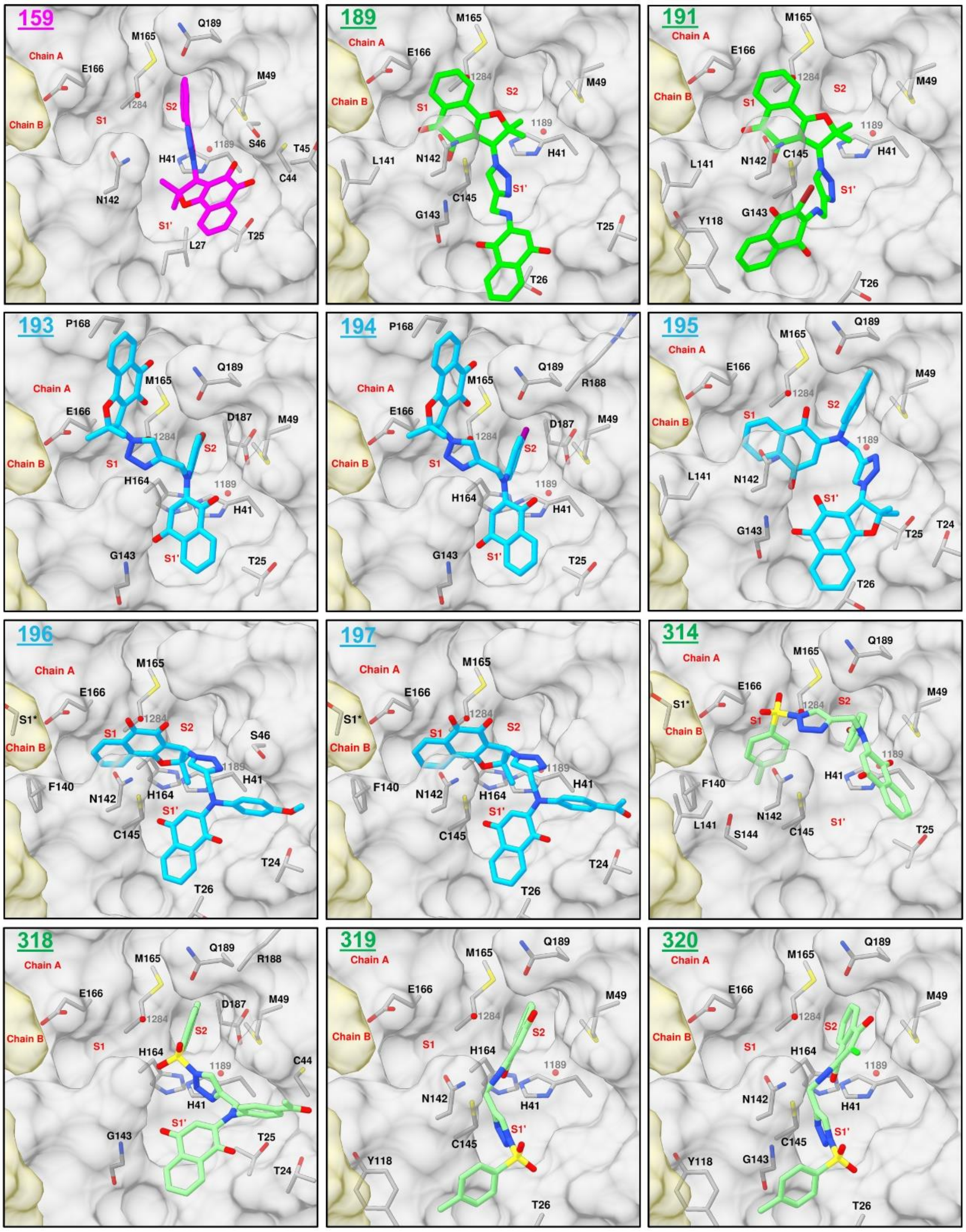

**Figure S44.** Docking binding modes from Glide of compounds **159**, **189**, **191**, **193**, **194**, **195**, **196**, **197**, **314**, **318**, **319**, and **320**, prioritized for evaluation in biochemical assays. The protein interacting residues are depicted as gray sticks.

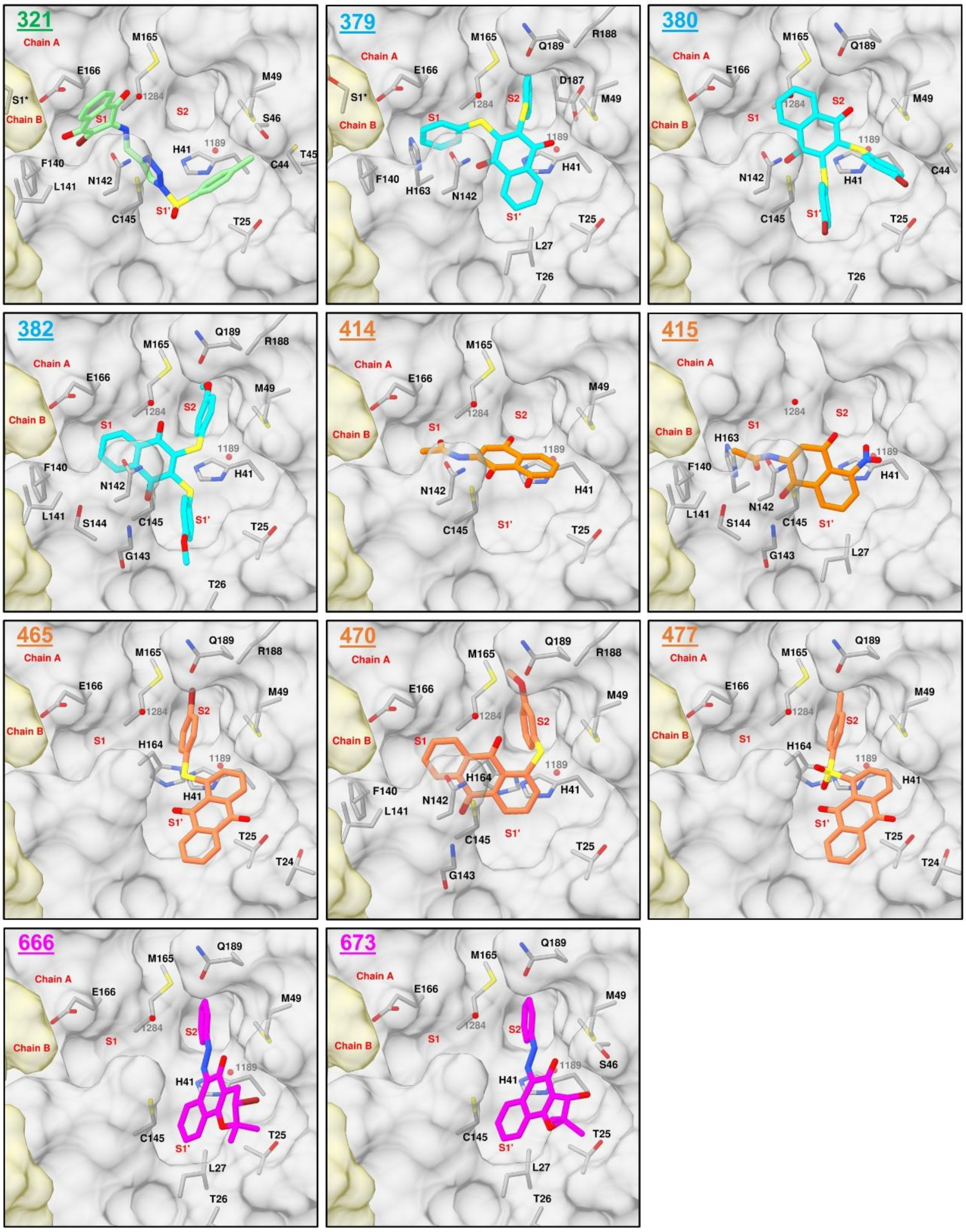

**Figure S45.** Docking binding modes from Glide of compounds **321**, **379**, **380**, **382**, **414**, **415**, **465**, **470**, **477**, **666**, and **673**, prioritized for evaluation in biochemical assays. The protein interacting residues are depicted as gray sticks.

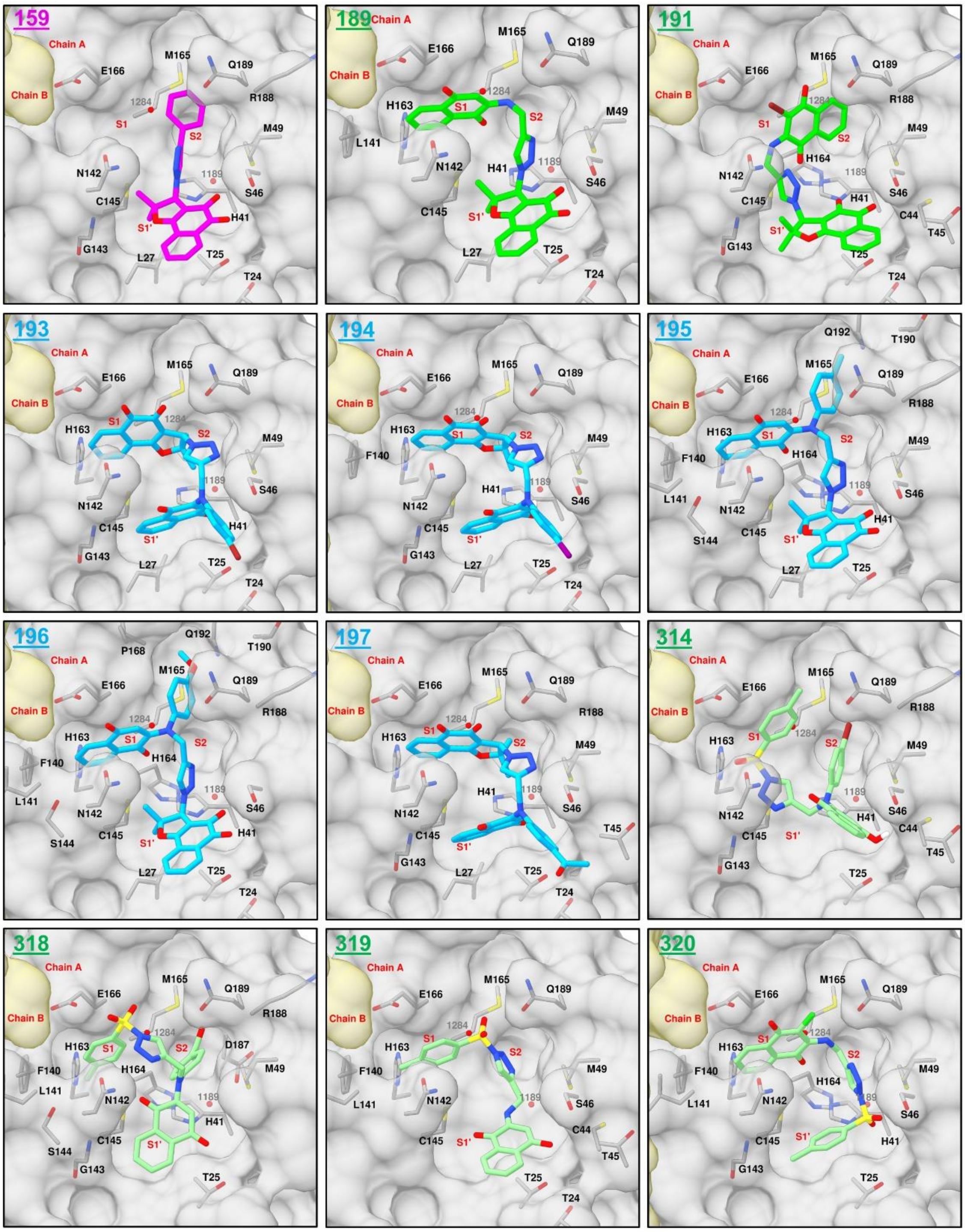

**Figure S46.** Docking binding modes from Vina of compounds **159**, **189**, **191**, **193**, **194**, **195**, **196**, **197**, **314**, **318**, **319**, and **320**, prioritized for evaluation in biochemical assays. The protein interacting residues are depicted as gray sticks.

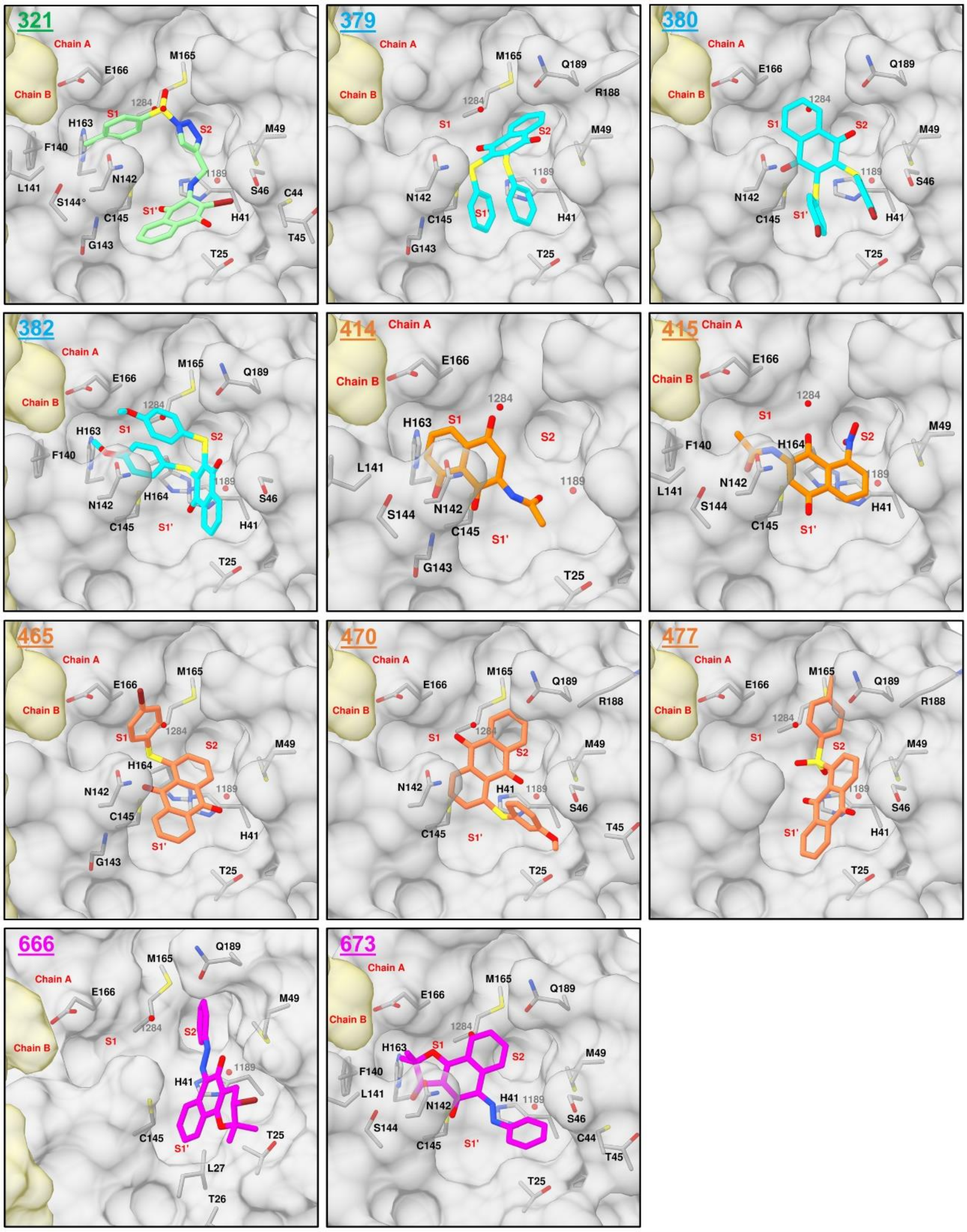

**Figure S47.** Docking binding modes from Vina of compounds **321**, **379**, **380**, **382**, **414**, **415**, **465**, **470**, **477**, **666**, and **673**, prioritized for evaluation in biochemical assays. The protein interacting residues are depicted as gray sticks.

#### 2.3 Validation of novel Mpro inhibitors

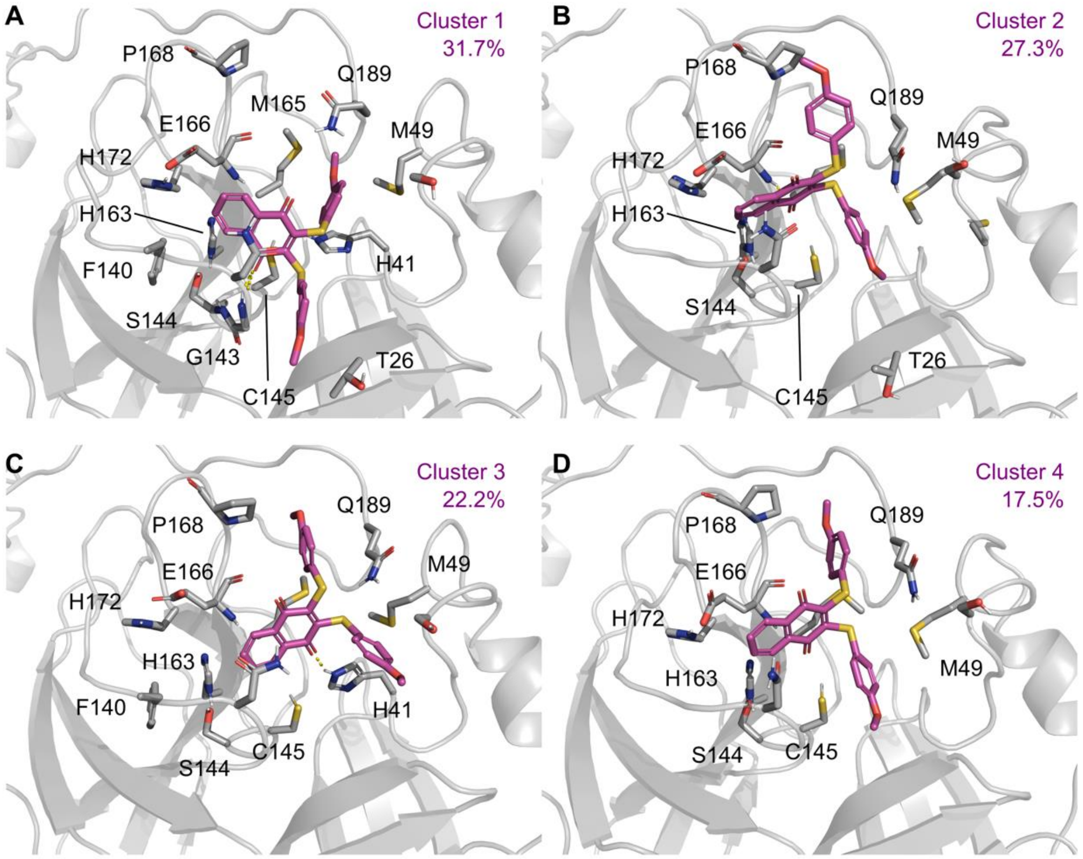

**Figure S48.** Clusters for compound 382 bound to Mpro. Clustering was performed using hierarchical clustering analyses, implemented in the trj_cluster.py script from Schrödinger, using the RMSD variation of the ligand with cut-off of 2.0 Å. Mpro residues are colored according to the types of atoms in the interacting amino acid residues (protein carbon, light gray; nitrogen, blue; oxygen, red), hydrogen bond interactions are represented as yellow dashed lines.

#### 2.4 Validation of novel PLpro inhibitors

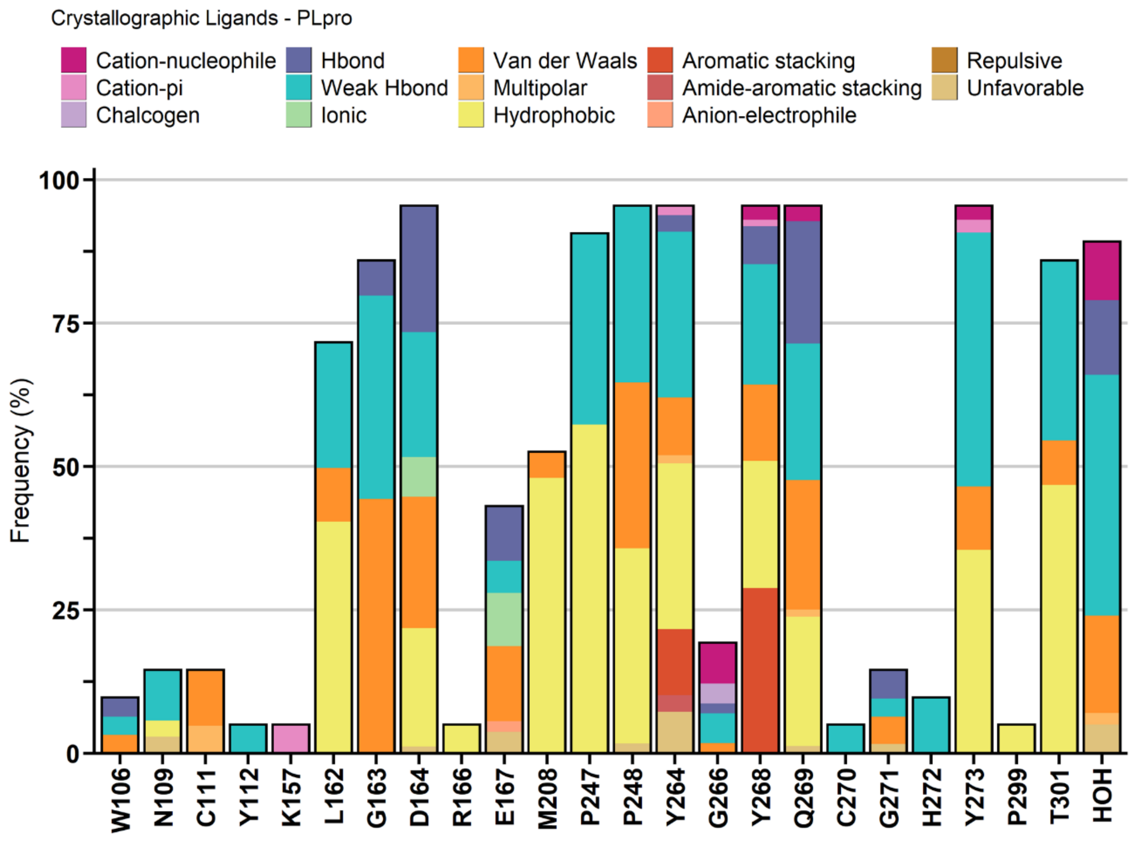

**Figure S49.** Protein-ligand interaction analysis from 21 PLpro complexes found in the PDB.

##### 2.4.1 Molecular dynamics simulations of PLpro bound to the crystallography ligand XR8-89

We performed simulations with the XR8-89 (PDB code 7LBR) as positive control. These simulations confirmed that the BL2 loop remained in the closed conformation, as originally proposed by the crystal structure. This BL2 loop conformation exposes a hydrophobic binding site by decreasing the solvent exposed surface. The binding mode of XR8-89 remained stable in all simulations, with its core structure being stabilized by hydrogen bond interactions between the carbonyl group and the Q269's backbone (during the entire simulation), as well as π-stacking interactions with Y268 (79% of the analyzed simulation time) (**Figure S50**). The water bridge between the amide's nitrogen and D164 (site S3), proposed in the original publication, was shown to be an intermittent in our simulations (present on average 22% of the analyzed trajectory). Interestingly, the amino-cyclopentanol moiety remained occupying the BL2 groove, sustained by hydrogen bond interactions with the G266 (present on average 81% of the analyzed simulation) and water-mediated interactions with N267 (32%). These characteristics were used to understand the results for our own ligands, since their proposed binding mode also initially relied on interactions with D164, Y268 and Q269, though with much greater variability in the simulations.

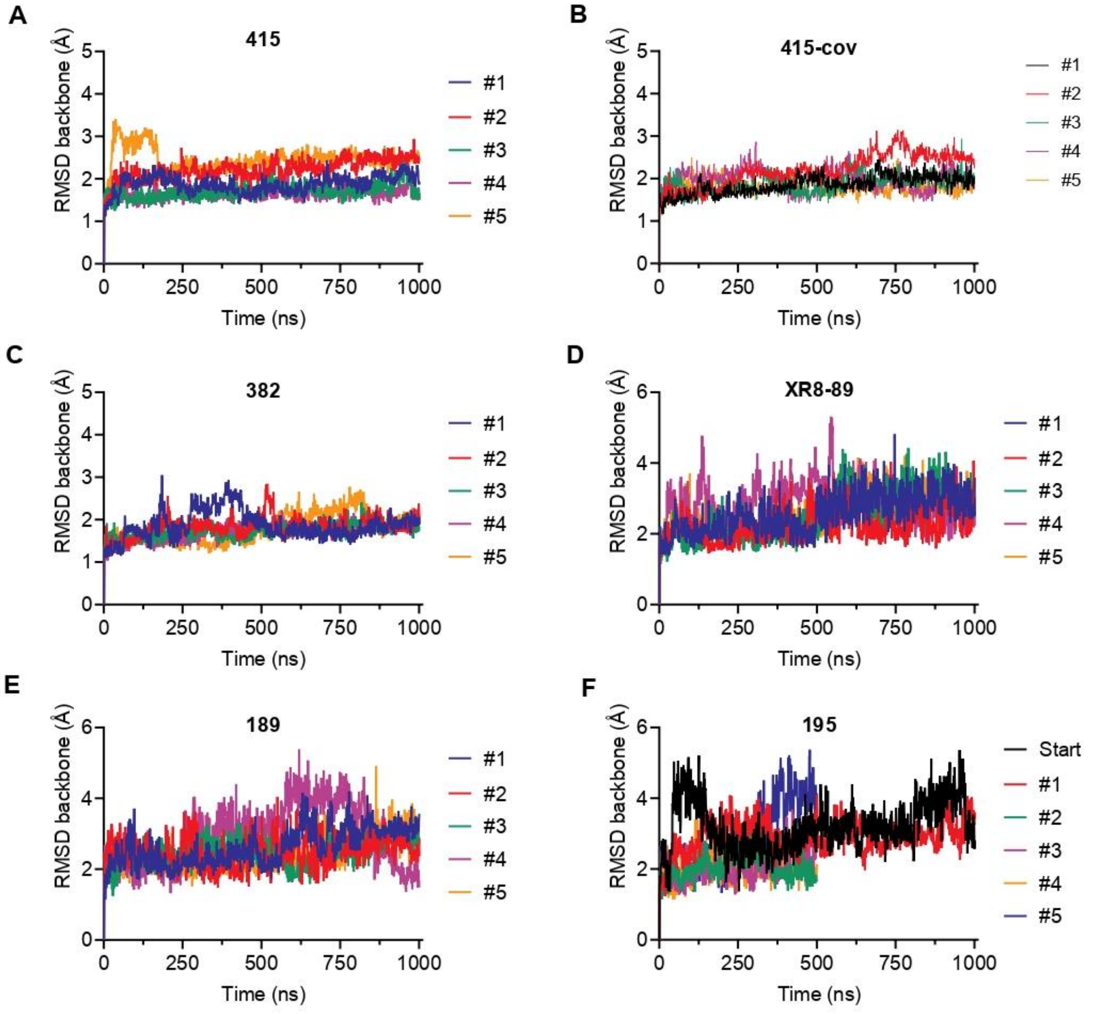

**Figure S50.** Root mean square deviation (RMSD) values long the simulation time for each replica (colored individually) for Mpro simulations with compound **415** (A), covalently bound 415 (B), and **382** (C), as well as for the co-crystallized ligand of PLpro (XR8-89, D) and our two identified hits **189** (E) and **195** (F). For the ligand **195** the initial simulation of 1 microsecond is depicted as a black line named Start, followed by five shorter replicas derived from the last frames on the start simulation.

#### 3. Evaluation of hit compounds in a SARS-CoV-2 viral infection assay

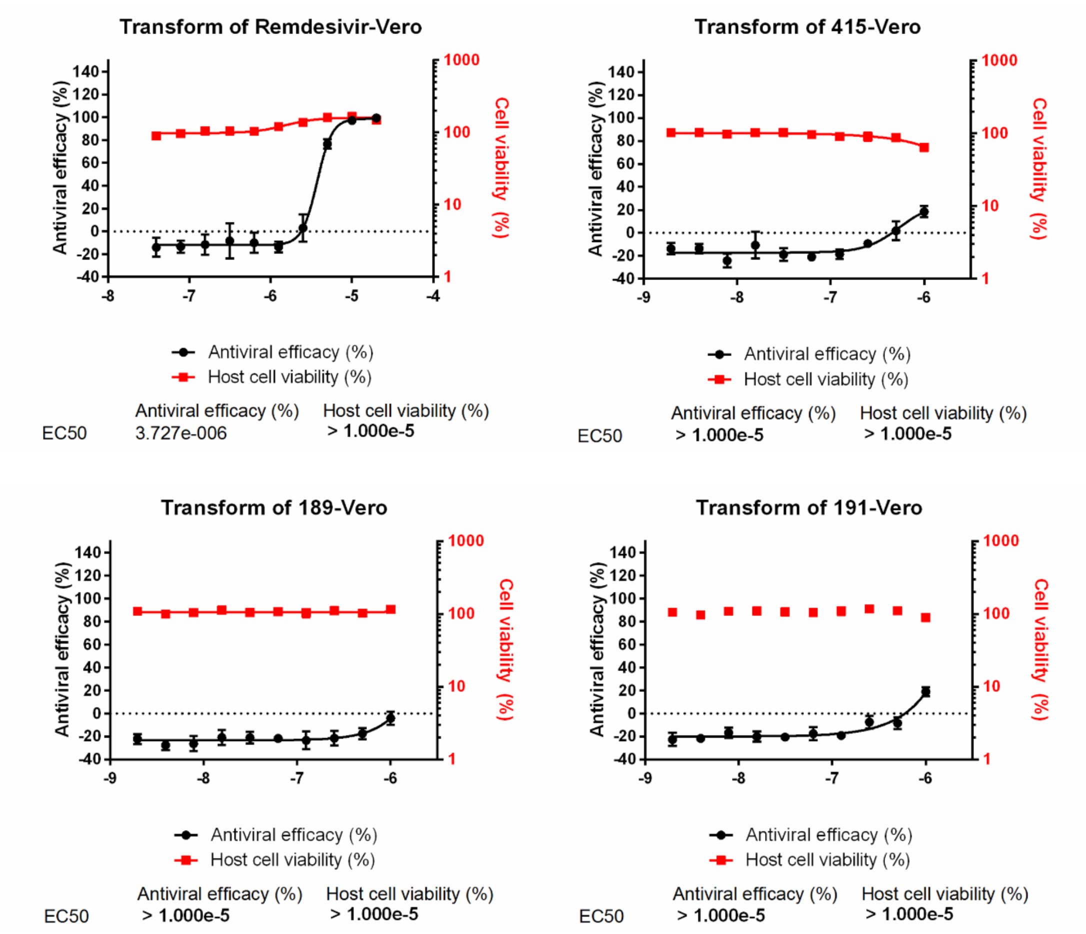

**Figure S51**. Evaluation of compounds **159**, **189**, **191** and **415** in a SARS-CoV-2 viral infection assay with Vero cells. Each compound was evaluated in 10 concentrations, in triplicate. Curves corresponding to antiviral efficacy (black) and Vero cell viability (red) are shown.

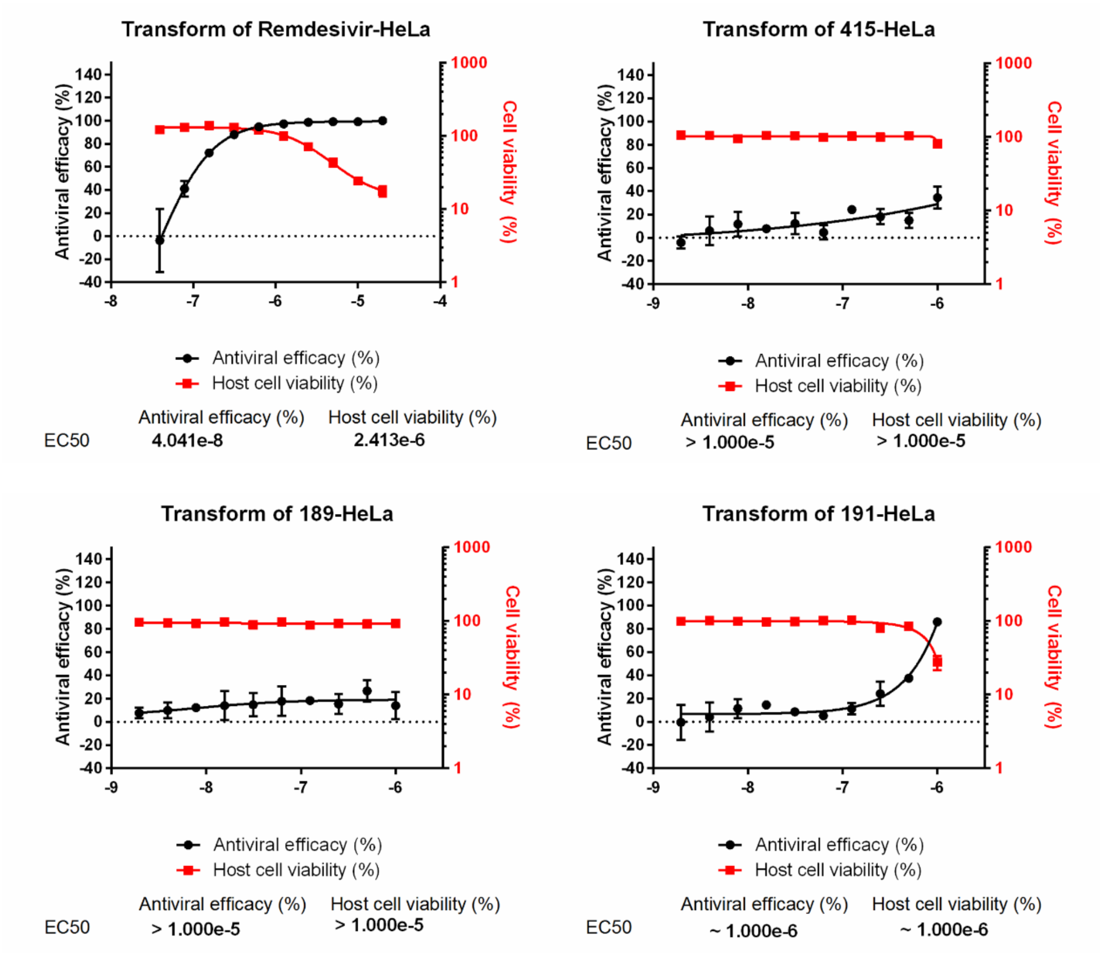

**Figure S52**. Evaluation of compounds **159**, **189**, **191** and **415** in a SARS-CoV-2 viral infection assay with HeLa cells. Each compound was evaluated in 10 concentrations, in triplicate. Curves corresponding to antiviral efficacy (black) and HeLa cell viability (red) are shown.

(1) da Silva Júnior, E. N.; de Souza, M. C. B. V.; Pinto, A. V.; Pinto, M. do C. F. R.; Goulart, M. O. F.; Barros, F. W. A.; Pessoa, C.; Costa-Lotufo, L. V.; Montenegro, R. C.; de Moraes, M. O.; Ferreira, V. F. Synthesis and potent antitumor activity of new arylamino derivatives of nor-β-lapachone and nor-α-lapachone. *Bioorg. Med. Chem.* **2007**, *15*, 7035-7041.

(2) da Silva Júnior, E. N.; de Souza, M. C. B. V.; Fernandes, M. C.; Menna-Barreto, R. F. S.; Pinto, M. do C. F. R.; de Assis Lopes, F.; de Simone, C. A.; Andrade, C. K. Z.; Pinto, A. V.; Ferreira, V. F.; de Castro, S. L. Synthesis and anti-Trypanosoma cruzi activity of derivatives from nor-lapachones and lapachones. *Bioorg. Med. Chem.* **2008**, *16*, 5030-5038.

(3) da Silva Júnior, E. N.; Guimarães, T. T.; Menna-Barreto, R. F. S.; Pinto, M. do C. F. R.; de Simone, C. A.; Pessoa, C.; Cavalcanti, B. C.; Sabino, J. R.; Andrade, C. K. Z.; Goulart, M. O. F.; de Castro, S. L.; Pinto, A. V. The evaluation of quinonoid compounds against *Trypanosoma cruzi*: Synthesis of imidazolic anthraquinones, nor-

β-lapachone derivatives and β-lapachone-based 1,2,3-triazoles. *Bioorg. Med. Chem*. **2010**, *18*, 3224-3230.

(4) da Silva, E. N.; De Deus, C. F.; Cavalcanti, B. C.; Pessoa, C.; Costa-Lotufo, L. V.; Montenegro, R. C.; De Moraes, M. O.; Pinto, M. D. C. F. R.; De Simone, C. A.; Ferreira, V. F.; Goulart, M. O. F.; Andrade, C. K. Z.; Pinto, A. V. 3-Arylamino and 3-alkoxy-nor-β-lapachone derivatives: Synthesis and cytotoxicity against cancer cell lines. *J. Med. Chem*. **2010**, *53*, 504-508.

(5) Souza, A. A. D.; De Moura, M. A. B.; Abreu, F. C. D.; Goulart, M. O.; da Silva Júnior, E. N.; Pinto, A. V., Ferreira, V. F; Moscoso, R.; Nunez-Vergara, L.; Squella, J. A. Electrochemical study, on mercury, of a Meta-nitroarylamine derivative of nor-β-lapachone, an antitumor and trypanocidal compound. *Quím. Nova*. **2010**, *33*, 2075-2079.

(6) da Cruz, E. H.; Silvers, M. A.; Jardim, G. A. M.; Resende, J. M.; Cavalcanti, B. C.; Bom, I. S.; Pessoa, C.; Simone, C. A.; de Botteselle, G. V.; Braga, A. L.; Nair, D. K.; Namboothiri, I. N. N.; Da Silva Júnior, E. N.; Boothman, D. A. Synthesis and antitumor activity of selenium-containing quinone-based triazoles possessing two redox centres, and their mechanistic insights. *Eur. J. Med. Chem*. **2016**, *122*, 1-16.

(7) Kharma, A.; Jacob, C.; Bozzi, Í. A. O.; Jardim, G. A. M.; Braga, A. L.; Salomão, K.; Gatto, C. C.; Silva, M. F. S.; Pessoa, C.; Stangier, M.; Ackermann, L.; da Silva Júnior, E. N. Electrochemical Selenation/Cyclization of Quinones: A Rapid, Green and Efficient Access to Functionalized Trypanocidal and Antitumor Compounds. *Eur. J. Org. Chem*. **2020**, *29*, 4474-4486.

(8) Jardim, G. A. M.; Reis, W. J.; Ribeiro, M. F.; Ottoni, F. M.; Alves, R. J.; Silva, T. L.; Goulart, M. O. F.; Braga, A. L.; Menna-Barreto, R. F. S.; Salomão, K.; de Castro, S. L.; da Silva Júnior, E. N. On the investigation of hybrid quinones: Synthesis, electrochemical studies and evaluation of trypanocidal activity. *RSC Adv*. **2015**, *5*, 78047-78060.

(9) Vieira, A. A.; Brandão, I. R.; Valença, W. O.; De Simone, C. A.; Cavalcanti, B. C.; Pessoa, C.; Carneiro, T. R.; Braga, A. L.; da Silva Júnior, E. N. Hybrid compounds with two redox centres: Modular synthesis of chalcogen-containing lapachones and studies on their antitumor activity. *Eur. J. Med. Chem*. **2015**, *101*, 254-265.

(10) Almeida, R. G.; Valença, W. O.; Rosa, L. G.; De Simone, C. A.; de Castro, S. L.; Barbosa, J. M. C.; Pinheiro, D. P.; Paier, C. R. K.; De Carvalho, G. G. C.; Pessoa, C.; Goulart, M. O. F.; Kharma, A.; da Silva Júnior, E. N. Synthesis of quinone imine and sulphur-containing compounds with antitumor and trypanocidal activities: Redox and biological implications. *RSC Med. Chem*. **2020**, *11*, 1145-1160.

(11) Jardim, G. A. M.; Guimarães, T. T.; Pinto, M. D. C. F. R.; Cavalcanti, B. C.; De Farias, K. M.; Pessoa, C.; Gatto, C. C.; Nair, D. K.; Namboothiri, I. N. N.; da Silva Júnior, E. N. Naphthoquinone-based chalcone hybrids and derivatives: Synthesis and potent activity against cancer cell lines. *MedChemComm*. **2015**, *6*, 120-150.

(12)Lima, D. J. B.; Almeida, R. G.; Jardim, G. A. M.; Barbosa, B. P. A.; Santos, A. C. C.; Valença, W. O.; Scheide, M. R.; Gatto, C. C.; de Carvalho, G. G. C.; Costa, P. M. S.; Pessoa, C.; Pereira, C. L. M.; Jacob, C.; Braga, A. L.; da Silva Júnior, E. N. It takes two to tango: synthesis of cytotoxic quinones containing two redox active centers with potential antitumor activity, RSC Med. Chem. (2021). https://doi.org/10.1039/d1md00168j

(13) Cavalcanti, B. C.; Cabral, I. O.; Rodrigues, F. A. R.; Barros, F. W. A.; Rocha, D. D.; Magalhães, H. I. F., Moura, D. J.; Saffi, J.; Henriques, J. A. P.; Carvalho, T. S. C.;

Moraes, M. O.; Pessoa, C.; De Melo, I. M. M.; da Silva Júnior, E. N. Potent antileukemic action of naphthoquinoidal compounds: Evidence for an intrinsic death mechanism based on oxidative stress and inhibition of DNA repair. *J. Braz. Chem. Soc.* **2013**, *24*, 145-163.

(14) de Castro, S. L.; Emery, F. S.; da Silva Júnior, E. N. Synthesis of quinoidal molecules: Strategies towards bioactive compounds with an emphasis on lapachones. *Eur. J. Med. Chem.* **2013**, *69*, 678-700.

(15) Dias, G. G.; Rogge, T.; Kuniyil, R.; Jacob, C.; Menna-Barreto, R. F. S.; da Silva Júnior, E. N.; Ackermann, L. Ruthenium-catalyzed C-H oxygenation of quinones by weak O-coordination for potent trypanocidal agents. *Chem. Commun.* **2018**, *54*, 12840-12843.

(16) Baiju, T. V.; Almeida, R. G.; Sivanandan, S. T.; de Simone, C. A.; Brito, L. M.; Cavalcanti, B. C.; Pessoa, C.; Namboothiri, I. N. N.; da Silva Júnior, E. N. Quinonoid compounds via reactions of lawsone and 2-aminonaphthoquinone with α-bromonitroalkenes and nitroallylic acetates: Structural diversity by C-ring modification and cytotoxic evaluation against cancer cells. *Eur. J. Med. Chem.* **2018**, *151*, 686-704.

(17) Wood, J. M.; Satam, N. S.; Almeida, R. G.; Cristani, V. S.; de Lima, D. P.; Dantas-Pereira, L.; Salomão, K.; Menna-Barreto, R. F. S.; Namboothiri, I. N. N.; Bower, J. F.; da Silva Júnior, E. N. Strategies towards potent trypanocidal drugs: Application of Rh-catalyzed [2+2+2] cycloadditions, sulfonyl phthalide annulation and nitroalkene reactions for the synthesis of substituted quinones and their evaluation against Trypanosoma cruzi. *Bioorg. Med. Chem.* **2020**, *28*, 115565.

(18) Suginome, H.; Konishi, A.; Sakurai, H.; Minakawa, H.; Takeda, T.; Senboku, H.; Tokuda, M.; Kobayashi, K. Photoinduced molecular transformations. Part 156. New photoadditions of 2-hydroxy-1, 4-naphthoquinones with naphthols and their derivatiyes. *Tetrahedron*, **1995**, *51*, 1377-1386.

(19) da Silva Júnior, E. N.; de Carvalho, R. L.; Almeida, R. G.; Rosa, L. G.; Fantuzzi, F.; Rogge, T.; Costa, P. M. S.; Pessoa, C.; Jacob, C.; Ackermann, L. Ruthenium(II)-Catalyzed Double Annulation of Quinones: Step-Economical Access to Valuable Bioactive Compounds. *Chem. Eur. J.* **2020**, *26*, 10981-10986

(20) da Silva, E. N.; Menna-Barreto, R. F. S.; Pinto, M. do C. F. R.; Silva, R. S. F.; Teixeira, D. V.; de Souza, M. C. B. V.; De Simone, C. A.; De Castro, S. L.; Ferreira, V. F.; Pinto, A. V. Naphthoquinoidal [1,2,3]-triazole, a new structural moiety active against Trypanosoma cruzi. *Eur. J. Med. Chem.* **2008**, *43*, 1774-1780.

(21) da Silva Júnior, E. N.; de Moura, M. A. B. F.; Pinto, A. V.; Pinto, M. do C. F. R.; de Souza, M. C. B. V.; Araújo, A. J.; Pessoa, C.; Costa-Lotufo, L. V.; Montenegro, R. C.; de Moraes, M. O.; Ferreira, V. F.; Goulart, M. O. F. Cytotoxic, trypanocidal activities and physicochemical parameters of nor-β-lapachone-based 1,2,3-triazoles. *J. Braz. Chem. Soc.* **2009**, *20*, 635-643.

(22) da Silva, E. N.; De Melo, I. M. M.; Diogo, E. B. T.; Costa, V. A.; De Souza Filho, J. D.; Valença, W. O.; Camara, C. A.; De Oliveira, R. N.; De Araujo, A. S.; Emery, F. S.; Dos Santos, M. R.; De Simone, C. A.; Menna-Barreto, R. F. S.; de Castro, S. L. On the search for potential anti-Trypanosoma cruzi drugs: Synthesis and biological evaluation of 2-hydroxy-3-methylamino and 1,2,3-triazolic naphthoquinoidal compounds obtained by click chemistry reactions. *Eur. J. Med. Chem.* **2012**, *52*, 304-312.

(23) Cardoso, M. F.; Rodrigues, P. C.; Oliveira, M. E. I.; Gama, I. L.; da Silva, I. M.; Santos, I. O.; Rocha, D. R.; Pinho, R. T.; Ferreira, V. F.; de Souza, M. C. B. V.; da Silva, F. C.; Silva-Jr, F. P. Synthesis and evaluation of the cytotoxic activity of 1, 2-furanonaphthoquinones tethered to 1, 2, 3-1H-triazoles in myeloid and lymphoid leukemia cell lines *Eur. J. Med. Chem.* **2014**, *84*, 708-717.

(24) Jardim, G. A. M.; Cruz, E. H. G.; Valença, W. O.; Resende, J. M.; Rodrigues, B. L.; Ramos, D. F.; Oliveira, R. N.; Silva, P. E. A.; da Silva Júnior, E. N. On the search for potential antimycobacterial drugs: Synthesis of naphthoquinoidal, phenazinic and 1,2,3-triazolic compounds and evaluation against Mycobacterium tuberculosis. *J. Braz. Chem. Soc.* **2015**, *26*, 1013-1027.

(25) dos Santos, F. S.; Dias, G. G.; de Freitas, R. P.; Santos, L. S.; de Lima, G. F.; Duarte, H. A.; de Simone, C. A.; Rezende, L. M. S. L.; Vianna, M. J. X.; Correa, J. R.; Neto, B. A. D.; da Silva Júnior, E. N. Redox Center Modification of Lapachones towards the Synthesis of Nitrogen Heterocycles as Selective Fluorescent Mitochondrial Imaging Probes. *Eur. J. Org. Chem.* **2017**, *2017*, 3763–3773.

(26) Bahia, S. B. B. B.; Reis, W. J.; Jardim, G. A. M.; Souto, F. T.; De Simone, C. A.; Gatto, C. C.; Menna-Barreto, R. F. S.; de Castro, S. L.; Cavalcanti, B. C.; Pessoa, C.; Araujo, M. H.; da Silva Júnior, E. N. Molecular hybridization as a powerful tool towards multitarget quinoidal systems: Synthesis, trypanocidal and antitumor activities of naphthoquinone-based 5-iodo-1,4-disubstituted-, 1,4- and 1,5-disubstituted-1,2,3-triazoles. *MedChemComm*, **2016**, *7*, 1555-1563.

(27) Jardim, G. A. M.; Da Cruz, E. H. G.; Valença, W. O.; Lima, D. J. B.; Cavalcanti, B. C.; Pessoa, C.; Rafique, J.; Braga A. L.; Jacob, C.; da Silva, E. N. Synthesis of selenium-quinone hybrid compounds with potential antitumor activity via Rh-Catalyzed C-H bond activation and click reactions. *Molecules.* **2018**, *23*, 1-16.

(28) Diogo, E. B. T.; Dias, G. G.; Rodrigues, B. L.; Guimarães, T. T.; Valença, W. O.; Camara, C. A.; De Oliveira, R. N.; da Silva, M. G.; Ferreira, V. F.; De Paiva, Y. G.; Goulart, M. O. F.; Menna-Barreto, R. F. S.; de Castro, S. L.; da Silva Júnior, E. N. Synthesis and anti-Trypanosoma cruzi activity of naphthoquinone-containing triazoles: Electrochemical studies on the effects of the quinoidal moiety. *Bioorg. Med. Chem.* **2013**, *21*, 6337-6348.

(29) Gontijo, T. B.; de Freitas, R. P.; Emery, F. S.; Pedrosa, L. F.; Vieira Neto, J. B.; Cavalcanti, B. C.; Pessoa, C.; King, A.; de Moliner, F.; Vendrell, M.; da Silva Júnior, E. N. On the synthesis of quinone-based BODIPY hybrids: New insights on antitumor activity and mechanism of action in cancer cells. *Bioorg. Med. Chem. Lett.* **2017**, *27*, 4446-4456.

(30) Gontijo, T. B.; De Freitas, R. P.; De Lima, G. F.; De Rezende, L. C. D.; Pedrosa, L. F.; Silva, T. L.; Goulart, M. O. F.; Cavalcanti, B. C.; Pessoa, C.; Bruno, M. P.; Corrêa, J. R.; Emery, F. S.; da Silva Júnior E. N. Novel fluorescent lapachone-based BODIPY: Synthesis, computational and electrochemical aspects, and subcellular localisation of a potent antitumour hybrid quinone. *Chem. Commun.* **2016**, *52*, 13281-13284.

(31) Melo, V. N.; Dantas, W. M.; Camara, C. A.; de Oliveira, R. N. Synthesis of 2, 3-unsaturated alkynyl O-glucosides from tri-O-acetyl-D-glucal by using montmorillonite K-10/iron (III) chloride hexahydrate with subsequent copper (I)-catalyzed 1, 3-dipolar cycloaddition. *Synthesis*, **2015**, *47*, 3529-3541.

(32) de Oliveira, R. N.; Xavier, A. D. L.; Guimaraes, B. M.; Melo, V. N.; Valenca, W. O.; Do Nascimento, W. S; Da Costa, P. F.; Camara, C. A. Combining clays and ultraso

und irradiation for an *O*-acetylation reaction of N-glucopyranosyl and other molecules. *J. Chil. Chem. Soc*, **2014**, *59*, 2610-2614.
(33) da Cruz, E. H.; Hussene, C. M.; Dias, G. G.; Diogo, E. B.; De Melo, I. M.; Rodrigues, B. L.; Silva, M. G.; Valença, W. O.; Camara, C. A.; de Oliveira R. N., Paiva, Y. G.; Goulart, M. O. F.; Cavalcanti, B. C.; Pessoa, C.; da Silva Júnior, E. N. 1, 2, 3-Triazole-, arylamino-and thio-substituted 1, 4-naphthoquinones: Potent antitumor activity, electrochemical aspects, and bioisosteric replacement of C-ring-modified lapachones. *Bioorg. Med. Chem*. **2014**, 22, 1608-1619.
(34) Valença, W. O.; Baiju, T. V.; Brito, F. G.; Araujo, M. H.; Pessoa, C.; Cavalcanti, B. C.; de Simone, C. A.; Jacob, C.; Namboothiri, I. N. N.; da Silva Júnior, E. N. Synthesis of Quinone-Based N -Sulfonyl-1,2,3-triazoles: Chemical Reactivity of Rh(II) Azavinyl Carbenes and Antitumor Activity. *ChemistrySelect.* **2017**, *2,* 4301-4308
(35) Cheng, Z.; Valença, W. O.; Dias, G. G.; Scott, J.; Barth, D. N.; Moliner, F.; Souza, G. B. P.; Mellanbyc, R. J.; Vendrell, M.; da Silva Júnior, E. N. *Bioorg. Med. Chem.* **2019**, *27*, 3938-3946.
(36) Rostovtsev, V. V.; Green. L. G.; Fokin, V. V.; Sharpless, K. B. *Angew. Chem. Int. Ed. Engl.* **2002**, *41*, 2599.
(37) Moliner, F.; King, A.; Dias, G. G.; de Lima, G. F.; de Simone, C. A.; da Silva Júnior, E. N.; Vendrell, M. Quinone-derived π-extended phenazines as new fluorogenic probes for live-cell imaging of lipid droplets. *Front.Chem*. **2018**, *6*, 339.
(38) Silva, R. S.; Amorim, M. B. D.; Pinto, M. D. C. F.; Emery, F. S.; Goulart, M. O.; Pinto, A. V. Chemoselective oxidation of benzophenazines by m-CPBA: n-oxidation vs. oxidative cleavage. *J. Braz. Chem. Soc.* **2007**, *18*, 759-764.
(39) Kulangiappar, K.; Anbukulandainathan, M., Raju; T. Synthetic communications: an international journal for rapid communication of synthetic organic chemistry. *Synth. Commun.* **2014**,*1*, 2494-2502.
(40) Errante, G.; La Motta, G.; Lagana, C.; Wittebolle, V.; Sarciron, M. É.; Barret, R. Synthesis and evaluation of antifungal activity of naphthoquinone derivatives. *Eur. J. Med. Chem.* **2006**, *41*, 773-778.
(41) Almeida, R. G.; De Carvalho, R. L.; Nunes, M. P.; Gomes, R. S.; Pedrosa, L. F.; De Simone, C. A.; Gopi, E.; Geertsen, V.; Gravel, E.; Doris, E.; da Silva Júnior, E. N. Carbon nanotube-ruthenium hybrid towards mild oxidation of sulfides to sulfones: Efficient synthesis of diverse sulfonyl compounds. *Catal. Sci. Technol*. **2019**, *9*, 2742-2748.
(42) Jardim, G. A. M.; Oliveira, W. X. C.; de Freitas, R. P.; Menna-Barreto, R. F. S.; Silva, T. L.; Goulart, M. O. F.; da Silva Júnior, E. N. Direct sequential C–H iodination/organoyl-thiolation for the benzenoid A-ring modification of quinonoid deactivated systems: a new protocol for potent trypanocidal quinones. *Org. Biomol. Chem.* **2018**, *16*, 1686-1691.
(43) Jardim, G. A. M.; Bozzi, Í. A. O.; Oliveira, W. X. C.; Mesquita-Rodrigues, C.; Menna-Barreto, R. F. S.; Kumar, R. A.; Gravel, E.; Doris, E.; Braga, A. L.; da Silva Júnior, E. N. Copper complexes and carbon nanotube-copper ferrite-catalyzed benzenoid A-ring selenation of quinones: An efficient method for the synthesis of trypanocidal agents. *New. J. Chem*. **2019**, *43*, 13751-13763.
(44) Jardim, G. A. M.; Silva, T. L.; Goulart, M. O. F.; de Simone, C. A.; Barbosa, Juliana M. C.; Salomao, K.; de Castro, S. L.; Bower, J. F.; da Silva Júnior, E. N. Rhodium-catalyzed C-H bond activation for the synthesis of quinonoid compounds: Significant

Anti- Trypanosoma cruzi activities and electrochemical studies of functionalized quinones. *Eur. J. Med. Chem.* **2017**, *136,* 406-419.
(45) Jardim, G. A. M.; Bower, J. F.; da Silva Júnior, E. N. Rh-Catalyzed Reactions of 1,4-Benzoquinones with Electrophiles: C–H Iodination, Bromination, and Phenylselenation. *Org. Lett.* **2016**, *18*, 4454-4457.
(46) Dias, G. G.; Nascimento, T. A. d.; de Almeida, A. K. A.; Bombaça, A. C. S.; Menna-Barreto, R. F. S.; Jacob, C.; Warratz, S.; da Silva Júnior, E. N.; Ackermann, L. Ruthenium(II)-Catalyzed C–H Alkenylation of Quinones: Diversity-Oriented Strategy for Trypanocidal Compounds. *Eur. J. Org*. Chem. **2019**, *2019*, 2344-2353.
(47) Sunassee, S. N.; Veale, C. G. L.; Shunmoogam-Gounden, N.; Osoniyi, O.; Hendricks, D. T.; Caira, M. R.; De La Mare, J. A.; Edkins, A. L.; Pinto, A. V.; da Silva Júnior, E. N.; Davies-Coleman, M. T. Cytotoxicity of lapachol, β-lapachone and related synthetic 1,4-naphthoquinones against oesophageal cancer cells. *Eur. J. Med. Chem.* **2013**, *62*, 98-110.
(48) Kumar, T.; Satam, N.; Namboothiri, I.N.N. Hauser–Kraus Annulation of Phthalides with Nitroalkenes for the Synthesis of Fused and Spiro Heterocycles. *Eur. J. Org. Chem*. **2016**, 3316-3321.
(49) Suresh, A.; Baiju, T. V.; Kumar, T.; Namboothiri, I. N. N. Synthesis of Spiro- and Fused Heterocycles via (4+4) Annulation of Sulfonylphthalide with o-Hydroxystyrenyl Derivatives. *J. Org. Chem*. **2019**, *84*, 3158-3168.
(50) Wood, J. M.; da Silva, E. N.; Bower, J. F. Rh-Catalyzed [2+2+ 2] Cycloadditions with Benzoquinones: De Novo Access to Naphthoquinones for Lignan and Type II Polyketide Synthesis. *Org. Lett*. **2020**, *22*, 265-269.
(51) de Carvalho, R. L.; Jardim, G. A. M.; Santos, A. C. C.; Araujo, M. H.; Oliveira, W. X. C.; Bombaça, A. C. S.; Menna-Barreto, R. F. S.; Gopi, E.; Gravel, E.; Doris, E.; da Silva Júnior, E. N. Combination of Aryl Diselenides/Hydrogen Peroxide and Carbon-Nanotube/Rhodium Nanohybrids for Naphthol Oxidation: An Efficient Route towards Trypanocidal Quinones. *Chem. Eur. J*. **2018**, *24*, 15227-15235.
(52) Jardim, G. A.; da Silva Júnior, E. N.; Bower, J. F. Overcoming naphthoquinone deactivation: rhodium-catalyzed C-5 selective C–H iodination as a gateway to functionalized derivatives. *Chem. Sci*. **2016**, *7*, 3780-3784.
(53) Kalt, M. M.; Schuehly, W.; Saf, R.; Ochensberger, S.; Solnier, J.; Bucar, F.; Presser, A. Palladium-catalysed synthesis of arylnaphthoquinones as antiprotozoal and antimycobacterial agents. *Eur. J. Med. Chem*. **2020**, *207*, 112837.
(54) Chen, L.; Xie, Y. Z.; Luo, Z. Y.; Liu, L. J.; Zou, Z. Z.; Liu, H. D.; Liu, S. Y. Synthesis and biological evaluation of novel isothiazoloquinoline quinone analogues. *Bioorg. Med. Chem. Lett.* **2020** *30*, 127286.
(55) Reis, W. J.; Bozzi, Í. A.; Ribeiro, M. F.; Halicki, P. C.; Ferreira, L. A.; da Silva, P. E. A.; Ramos, D. F.; de Simone C. A.; da Silva Júnior, E. N. Design of hybrid molecules as antimycobacterial compounds: Synthesis of isoniazid-naphthoquinone derivatives and their activity against susceptible and resistant strains of Mycobacterium tuberculosis. *Bioorg. Med. Chem*. **2019**, *27*, 4143-4150.
(56) Moura, K. C.; Carneiro, P. F.; Pinto, M. D. C. F.; da Silva, J. A.; Malta, V. R.; de Simone, C. A.; Dias, G. G.; Jardim, G. A. M.; Cantos, J.; Coelho, T. S.; Almeida, P. E.; da Silva Júnior, E. N. 1,3-Azoles from ortho-naphthoquinones: synthesis of aryl substituted imidazoles and oxazoles and their potent activity against Mycobacterium tuberculosis. *Bioorg. Med. Chem*. **2012**, *20*, 6482-6488.

(57) Dias, G. G.; Pinho, P. V.; Duarte, H. A.; Resende, J. M.; Rosa, A. B.; Correa, J. R.; Neto, B. A. D.; da Silva Júnior, E. N. Fluorescent oxazoles from quinones for bioimaging applications. *RSC Adv*. **2016**, *6*, 76056-76063.

(58) Dias, G. G.; Rodrigues, B. L.; Resende, J. M.; Calado, H. D.; de Simone, C. A.; Silva, V. H.; Neto, B. A. D. Goulart, M. O. F.; Ferreira, F. R.; Meira, A. S.; Pessoa, C.; Correa, J. R.; da Silva Júnior, E. N. Selective endocytic trafficking in live cells with fluorescent naphthoxazoles and their boron complexes. *Chem Commun.* **2015**, *51*, 9141-9144.
